# Systematic *in silico* discovery of novel solute carrier-like proteins from proteomes

**DOI:** 10.1101/2021.11.19.469292

**Authors:** Gergely Gyimesi, Matthias A. Hediger

## Abstract

Solute carrier (SLC) proteins represent the largest superfamily of transmembrane transporters. While many of them play key biological roles, their systematic analysis has been hampered by their functional and structural heterogeneity. Based on available nomenclature systems, we hypothesized that many as yet unidentified SLC transporters exist in the human genome, which await further systematic analysis. Here, we present criteria for defining “SLC-likeness” to curate a set of “SLC-like” protein families from the Transporter Classification Database (TCDB) and Protein families (Pfam) databases. Computational sequence similarity searches surprisingly identified ∼120 more proteins in human with potential SLC-like properties compared to previous annotations. Interestingly, several of these have documented transport activity in the scientific literature. To complete the overview of the “SLC-ome”, we present an algorithm to classify SLC-like proteins into protein families, investigating their known functions and evolutionary relationships to similar proteins from 6 other clinically relevant experimental organisms, and pinpoint structural orphans. We envision that our work will serve as a stepping stone for future studies of the biological function and the identification of the natural substrates of the many under-explored SLC transporters, as well as for the development of new therapeutic applications, including strategies for personalized medicine and drug delivery.

## Introduction

Membrane transporters and channels are the main entry routes for nutrients, ions, xenobiotics and serve as major exit routes for waste products and metabolites. The solute carrier (SLC) protein superfamily accounts for over 50% of all transport-related proteins and about 10% of all membrane proteins encoded by the human genome. With more than 400 annotated members, it is the largest superfamily of membrane transporter proteins [1,2]. The roles of SLC transporters as cellular gatekeepers, determinants of nutrient homeostasis and facilitators of drug metabolism and drug targeting has recently been revisited [3]. Around ∼50% of currently annotated SLCs are predicted to be associated with human disease phenotypes and many SLCs are considered to represent promising drug targets or drug delivery systems or to affect drug ADMET (absorption, distribution, metabolism, extrusion, toxicity). It has recently become evident that the SLC superfamily offers an enormous unexplored therapeutic treasure. But while the list of approved drugs that target transporter proteins is increasing, many promising SLCs still remain unexplored, uncharacterized and underrepresented in the literature.

The SLC nomenclature system has traditionally been used to classify mammalian secondary active and facilitative transporters, including exchangers and antiporters, into families based on sequence identity per enquiry by the Human Gene Nomenclature Committee (HGNC) starting in the early 1990s [1]. Originally, SLCs assignments have been made for all membrane transport proteins that are not channels, ATP-driven pumps, aquaporins, porins of the outer mitochondrial membrane or ATP-binding cassette (ABC) transporters, while usually having multiple transmembrane spanning segments and exhibiting transmembrane transport of a solute, or showing homology to membrane proteins having such features. Due to this construction, the SLC superfamily is structurally and functionally highly heterogeneous and thus most likely evolutionarily polyphyletic in origin. Because of these properties and the lack of common sequence patterns in the different SLC transporters, it has been difficult to assess how many SLC transporters exist in the human genome or which proteins could be predicted as SLC transporters. Proteins were typically added to the SLC system on a case-by-case basis. However, in view of recent requests to add new members into the SLC nomenclature, we suspected that the SLC system might be incomplete.

Despite their heterogeneity, SLC transporters seem to share common properties that probably were evolutionarily selected based on their suitability as facilitative transporters, secondary active transporters or exchangers. A remarkable property is that they generally have a symmetric inverted repeat architecture, which can be observed based on the available structures [4] and can sometimes also be detected at the sequence level [5,6]. Another important aspect is that the currently well-studied SLC transporters seem to follow an alternating access mechanism, meaning that the substrate-binding site is exposed on either one or the other side of the membrane, but not on both sides simultaneously [7,8]. Consequences of these properties are that SLC transporters typically contain many transmembrane helices (TMHs), and functionally exhibit saturable transport activity. In addition, most but not all SLC carriers transport water-soluble small molecules. These properties could be used as criteria to identify additional SLC transporter proteins.

In fact, there have been earlier attempts to gather additional SLCs from the human genome [9–11], as well as to classify them using automatic methods [12,13]. In this regard, one study [9] used BLAST searches to find SLC transporters that have local sequence similarities and found that 15 of the known SLC families fall into 4 phylogenetic clusters, which were termed α, β, γ, and δ groups. In addition, they have found 19 sequences that have previously not been described as SLCs. A later study [12] used a more sensitive HMM-HMM (hidden Markov-model) comparison-based method to identify locally similar regions in known SLC proteins. Visualization of the similarity network revealed visible protein clusters that correlated with existing SLC families. In addition, they identified two unannotated protein sequences that showed similarity to existing SLC proteins. A common limitation of these studies, however, is that they only searched for proteins that were similar to proteins already annotated as human SLC transporters. Nevertheless, these efforts using sequence similarity-based approaches have highlighted that there are additional as yet unannotated SLC transporters in human protein databases.

In our current study, we aimed to identify missing SLC transporters that may differ from those currently annotated in human. To do this, we turned to sequence databases and annotation systems that are phylogenetically broader and not limited to human proteins, and we developed criteria to define “SLC-like” proteins. Our method is thus more complete and general than previous approaches and tackles the task of identifying SLC transporters starting from first principles. For this reason, we turned to the Transporter Classification DataBase (TCDB) to enable the extraction of missing “SLC-like” proteins.

The Transporter Classification DataBase (TCDB) is an alternative classification system that was created in the 1990s in parallel to the SLC nomenclature series [14,15]. It collects transport-related membrane proteins, including membrane receptors, transporters, ion channels, and membrane-anchored enzymes from all kingdoms of life, with a particular focus on proteins from lower organisms. The proteins in the TCDB are organized hierarchically into subfamilies, families and superfamilies based on phylogenetic and functional considerations, and each member in the database is given a five-segmented TC# similar to the EC# that is used for enzyme classification. In addition, a brief description is provided for each family that introduces identified members and contains links to the most important relevant papers.

However, the TCDB dataset is not directly applicable for creating an overview of the collection of SLC transporters encoded in the human genome (the human SLC-ome). One of the reasons is that the TCDB is set up as a “representative database”, which means that it only contains certain representative sequences from each family. In addition, there is no particular focus on human proteins, and in fact several annotated human SLC transporters are not present in the database.

It also seems problematic to consider all proteins in the TCDB that are annotated as part of the secondary transporter superfamily TC# 2.A as being SLC-like. Indeed, many of the TCDB families annotated as part of TC# 2.A exhibit structural or sequence features that do not match the characteristics of existing SLCs. Examples of this are the Trk K^+^ transporters (#2.A.38), which display an ion-channel like structural fold [16], the GUP glycerol uptake proteins (#2.A.50), which exhibit enzymatic activity based on follow-up studies [17], and the Twin Arginine Targeting (Tat) family (#2.A.64), which are actually protein secretion complexes [18,19]. Since none of these families correspond to structural or functional characteristics of currently known SLC transporters, it is likely that not all proteins annotated under the TC# 2.A superfamily are “SLC-like”. Thus, while the TCDB could be a rich source of information for finding new transporters, it is clear that the perception of what a “transporter” should be according to TCDB does not always correspond to the typical properties of well-characterized SLC proteins. Additional filtering of TCDB data is therefore necessary.

As part of the TransportDB project, there were parallel efforts to collect secondary transporters from human and several other organisms [20–22]. Within the TransportDB project, the authors have built an automatic transporter annotation pipeline (TransAAP), which relies on BLAST searches, the Clusters of Orthologous Groups (COG) database [23], “selected HMMs for transporter protein families” [22] from the TIGRfam and Pfam databases [24,25] and hydropathy predictions of TMHMM [26]. This pipeline was used as a semiautomatic tool to annotate transporter-like proteins from the NCBI RefSeq database. However, based on the currently available TransportDB website, the resulting protein hits are neither linked to protein annotation databases, such as UniProt [27], nor are their official gene symbols or SLC names displayed. Therefore, no correlation with the existing SLC nomenclature is provided, and it is not trivial to say whether or not an existing SLC protein is included in the database.

We would also like to mention the Protein families (Pfam) database [28], which aims to maintain a curated set of protein families, often represented by functional domains. Notably, Pfam provides curated HMMs for each Pfam family to facilitate sequence similarity searches for the occurrence of those domains. In addition, Pfam groups protein families into higher-order groups called clans. Pfam clans contain evolutionarily related families whose relationships are supported either by sequence similarity, structural similarity or other orthogonal biological evidence [29]. While many known Pfam models correspond to the functional regions of known SLC transporters, Pfam neither attaches special importance to transport-related domains nor to membrane-spanning domains. For this reason, while the Pfam database could be a rich source of information about transporters, extracting Pfam families that encode transporter-like domains is non-trivial.

Therefore, there is a clear need in the field to define the criteria of “SLC-likeness” and to identify and classify all proteins in humans and other species that exhibit “SLC-likeness”. Thus, in our work, we interpret the term “SLC-likeness” by defining the essential criteria for it, and we carry out an exhaustive search for proteins that potentially meet these criteria, both with manual curation of datasets and with automatic sequence similarity-based approaches.

## Results

### Elaboration of criteria for “SLC-likeness”

Since the TCDB takes a very inclusive approach to collecting membrane transport-related proteins from a broad range of biological organisms, we have selected it as the source database for our endeavors. However, as outlined in the introduction, selecting SLC-like proteins from the TCDB is non-trivial. Therefore, we have introduced a set of criteria based on current knowledge of SLC transporters in order to select SLC-like protein families from the TCDB. We believe that these criteria represent the most important properties of currently known SLC transporters. The criteria used to infer SLC-likeness were as follows.

1. Structure of the protein should be α-helical, with at least three transmembrane helices (TMHs). Proteins with a β-barrel architecture, mostly β-structure proteins, membrane-anchored proteins, cyclic peptides and proteins consisting only of soluble domains (based on predictions or structural data) were excluded.
2. The size of the transported substrate should fall within the small-molecule range (i.e. oligopeptides might be accepted as substrates but protein secretion systems are excluded). Also excluded are DNA-, RNA- and polysaccharide-transporting systems.
3. Proteins with a channel-like mechanism were excluded, except in some rare cases. In particular, holins, toxins and other pore-forming proteins, and proteins bearing similarity to them, have been excluded.
4. Nucleotide-driven transporters (e.g. ATP-binding cassette/ABC families) were excluded.
5. Receptors that trigger endocytosis upon substrate binding were excluded. Only receptors were included where the receptor protein itself mediates the translocation of the substrate through the membrane, or the insertion of the substrate in the membrane, if that is its final location.
6. Proteins with enzymatic activity, or similarity to known enzymes were excluded. In some cases, where the protein was believed to contain both a transport domain and a soluble enzyme domain, the proteins have been included.
7. Proteins where transport activity was used as a synonym for trafficking (i.e. protein or vesicle translocation within the cell) but otherwise seemingly having no small-molecule transmembrane transport activity were excluded. On this basis, chaperones and other proteins helping the insertion of nascent proteins into a cellular membrane were also excluded.
8. For some (mostly putative) transporter families, TCDB does not give an explanation why the proteins would be considered as transporters. Families with no resemblance to known transporters and no indication or argument as to why they would be transporters were excluded.

We must emphasize that for specific proteins or protein families, the verification of some of the criteria for SLC-likeness as described above requires extensive and detailed information about the nature of the transport phenomenon. For example, deciding whether the substrate is a small molecule requires the identification of the transported solute, while ruling out a channel-like mechanism requires extensive information about the transport mechanism itself. However, due to missing information, these criteria could not always be verified while filtering the TCDB for “SLC-like” protein families. In certain cases, even the identity of the protein that performs the actual membrane translocation within a transporter complex can be unclear. For similar reasons, the historical discrepancy that the naming and classification of a gene/protein generally precludes the detailed analysis of its structure and function has also caused the currently known (“classic”) SLC nomenclature to contain transport proteins with a channel-like mechanism (e.g., the SLC41 family), single TMH (SLC27 family) as well as auxiliary proteins to actual transporters (SLC3 family). Some of these classic SLC families have been included in our selection in order to provide consistency with the existing SLC nomenclature despite the fact that they do not meet some of our criteria for SLC-likeness. Nevertheless, we believe that our criteria are broad enough to allow the identification of all putative SLC-like transporters, while also being specific enough to distinguish them from other well-known, non transport-related transmembrane protein families. Our criteria enabled us to initiate, for the first time, an attempt to set up guidelines defining SLC-likeness, in the sense that explicit criteria are established to disambiguate “transmembrane solute transport” and distinguish from other related membrane protein activities such as channel-like transport, receptor-mediated endocytosis or protein secretion. In our current study, we designate proteins and protein families that potentially meet the above criteria as “SLC-like”.

### Search for novel “SLC-like” proteins

The above-mentioned criteria were applied to manually select protein families in the TCDB that either fulfill or can potentially fulfill the criteria for SLC-likeness based on the description of each third-level family from the TCDB database. Throughout this manuscript, we use the term “TCDB family” to refer to third-level groupings in the classification hierarchy (i.e. TC# x.y.z), while “subfamily” and “superfamily” refer to fourth-level (TC# x.y.z.w) and second-level (TC# x.y) classes, respectively. In the work presented herein, we have analyzed superfamilies TC# 1.A, 2.A, 9.A and 9.B, as well as the families and in certain cases the subfamilies within them (see Table 1).

**Table 1.**
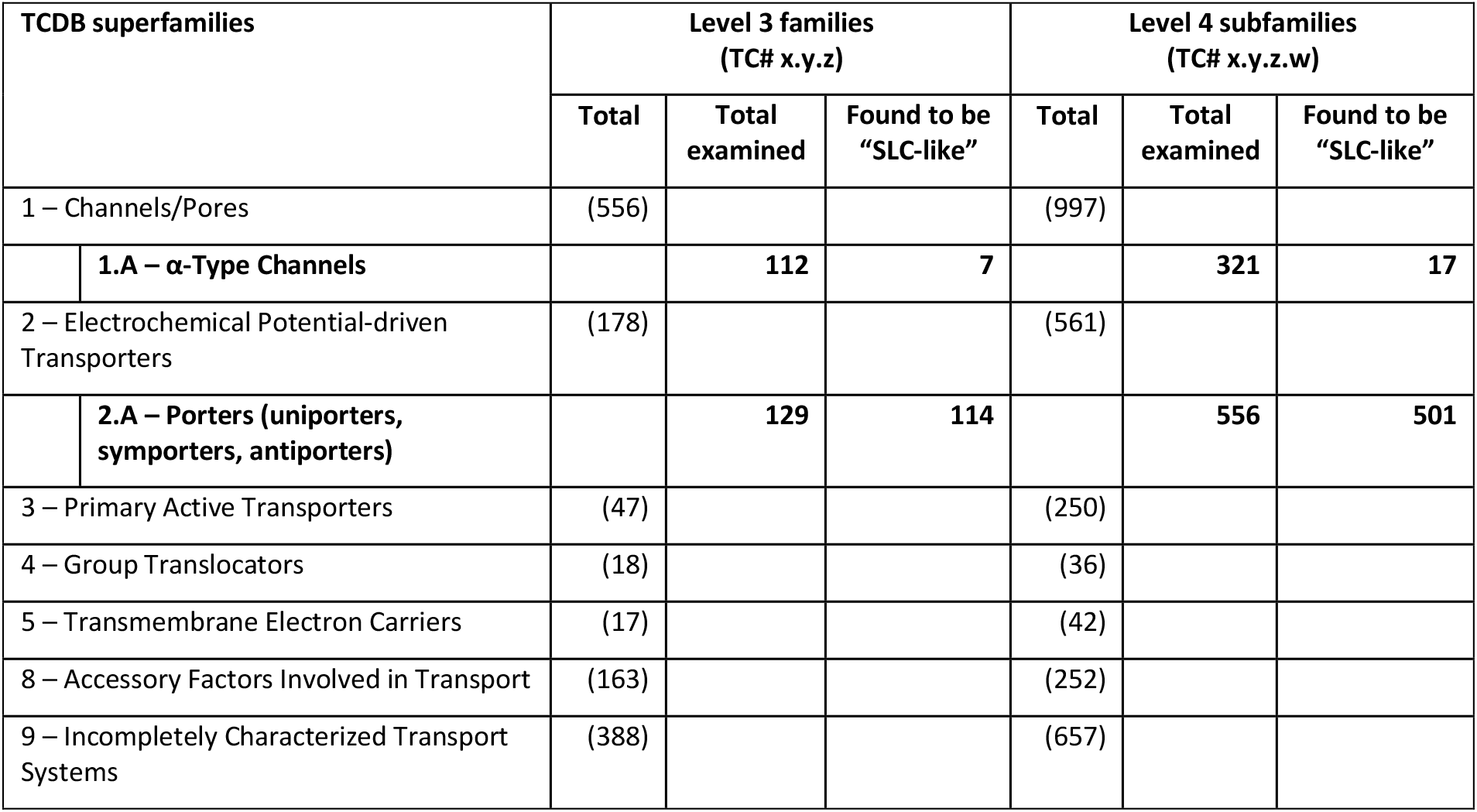
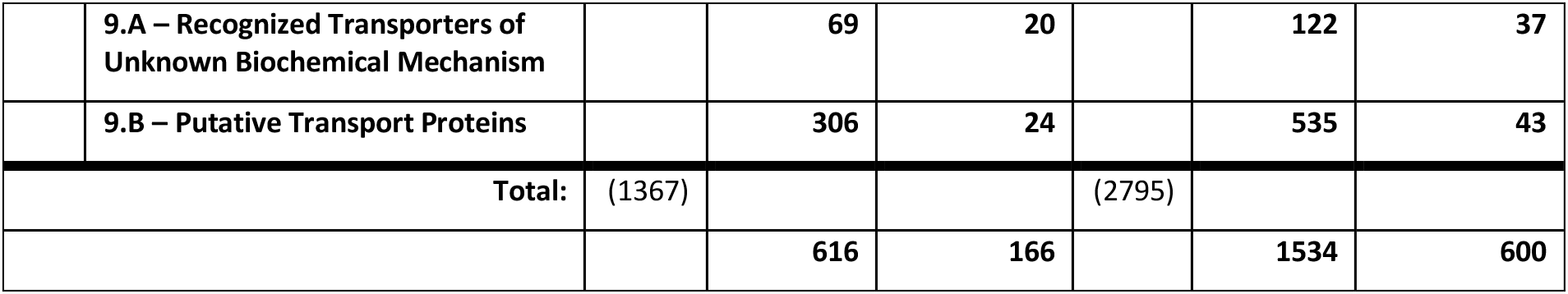
Families and subfamilies examined from the TCDB for SLC-like proteins. Superfamilies marked in boldface have been analyzed. The superfamily #1.A was examined since several existing SLC families are classified here, while superfamily #2.A was expected to contain most known SLC proteins. Superfamilies #9.A and #9.B were also examined. Total number of families and subfamilies in the first-level classes are indicated in parentheses. Numbers show the total number of level 3 families and level 4 subfamilies in each superfamily, as well as those found to be “SLC-like” (see text).

It was a significant curation effort to manually assess the 1534 subfamilies within 616 families in the above-mentioned superfamilies of the TCDB, which were expected to contain SLC transporter-like proteins (Table 1). To streamline our curation efforts, we have also expanded our analysis to Pfam protein families. To this end, we used HMMER [30] to search for all Pfam models in the sequences of all TCDB members within the four superfamilies analyzed, as shown in Table 1. The resulting Pfam families found in TCDB sequences were manually analyzed in tandem with the TCDB families and subfamilies in which they occur whether they also meet the criteria for SLC-likeness as defined above. The SLC-likeness criteria have been assessed according to the following guidelines:

- As an inclusive approach, we mainly searched for information that can be used to exclude (sub)families based on one or more criteria. Missing information (e.g. about transport mechanism, structure, potential other functions, exact chemical identity of substrate) was not interpreted as a reason for exclusion, unless it could be replaced by predictions (see below). Therefore, we strived to include (sub)families in our list that potentially fulfill the criteria for being SLC-like.
- When a (sub)family could be excluded from being SLC-like based on one criterion, the remaining criteria were not assessed.
- The functional criteria were assessed based on information presented on the TCDB web site in the family description text and in the description of certain individual members, where this was present, as well as in the textual descriptions of Pfam families. Families that claim their members to be “channels” in their descriptions were assumed to function with a channel-like mechanism and were therefore excluded, unless there was evidence for saturable transport or alternating access mechanism for certain members. TCDB and Pfam families showing similarity to other transporter families, either as mentioned in the family descriptions in text or *via* SCOOP-similarity [31], have been chosen to potentially fulfill the functional criteria, unless specific functional annotations indicated otherwise.
- The criterion about the size of the substrate was judged based on the known or putative substrate according to TCDB family descriptions. The assumption that certain proteins are transporters, whether based on experimental observations or purely on prediction, always entails the presence of either a tested or an assumed class of substrates. Families where no indication was available why they would function as transporters were thus excluded for the moment until further information on their function becomes available.
- The structural criterion was assessed based on the “number of transmembrane segments” annotations in the TCDB for each protein. (Sub)families where their TCDB descriptions mentioned a β-barrel structure or relationship with such proteins were excluded. Care was taken to identify outliers, i.e. proteins with less TMHs as the functional unit, which could potentially be fragments, and proteins with extra TMHs in addition to the functional core. In this regard, we based our decision on Pfam models conserved in the (sub)family and the number of THMs the model spans according to predictions at the Pfam web site for the most characterized members of the (sub)family. For certain Pfam families, structures and annotations from the Protein Data Bank (PDB) linked with the family on the Pfam web site were used to decide whether or not the Pfam model encodes soluble proteins, provided that the family model has sufficient coverage by the structure available.
- In some TCDB families with multicomponent transport systems, the membrane-spanning core components were identified by the presence of conserved Pfam models in all members of the family. Whether Pfam domains were membrane-spanning was deduced by their descriptions at the Pfam web site and Pfam prediction of TMHs. The TCDB description of the family also helped in deciding which components were essential and membrane-spanning. If multiple such Pfam models were found, the proteins were assumed to function as an obligate heteromultimer, and such proteins were considered as a single entity to assess the overall number of TMHs for the structural criterion.

Strikingly, of the 1534 subfamilies examined, only 600 in 166 families were found to be SLC-like according to our criteria. In particular, of the 556 subfamilies within superfamily TC# 2.A (“porters”), only 501 appeared to meet our selection criteria. This underlines once more that the term “porter” in relation to solute transport is ambiguous in this field and a clearer definition of the perception of a solute transporter protein is needed.

Our curation efforts also yielded 209 Pfam families bearing SLC-like properties, which likely contain the membrane-spanning regions of SLC-like proteins selected from the 336 Pfam families present in total in the TCDB sequences analyzed. Special care has been taken to consistently include or exclude TCDB (sub)families and their corresponding Pfam family, if applicable, from our selection. Notably, many of those Pfam families that have been excluded represented soluble structural or regulatory domains. Following our initial round of selection, we took advantage of clan groupings in Pfam and extended our selection efforts to analyze Pfam families belonging to the same clan as the selected 209 SLC-like Pfam families. This was based on the observation that many SLC-like transporter families in Pfam appear grouped into clans, so that members of these clans might represent SLC-like transporters themselves. As noted before, Pfam clans represent remotely related protein families, and while protein function is not always conserved across different families within a Pfam clan, the inspection of these families whether they meet our SLC-likeness criteria is validated by their relatedness or similarity to SLC-like families. Such a “clan expansion” procedure resulted in 12 additional Pfam families that are evolutionarily related to SLC-like protein families, of which 8 Pfam families potentially meet our criteria as SLC-like, as evaluated according to the guidelines and criteria above. Interestingly, these 8 SLC-like Pfam families currently have no representatives with a modest score (bit score > 25) in the TCDB, while 4 of the 8 families are annotated in Pfam as domains of unknown function (DUF). Thus, a total of 217 SLC-like Pfam families were identified in our search (see S1 Table).

We would like to note that our selection of SLC-like TCDB families and subfamilies as well as Pfam families is limited by the availability of information on less characterized protein families and may therefore need to be revised as more information becomes available. Our criteria are objective, however, and can easily be used to revise the decision as to whether a particular family is SLC-like in the light of new information.

As a next step, we wanted to know whether the selected families either from the TCDB or from Pfam have representatives in the proteomes of human and other clinically relevant organisms. For this analysis, we selected 7 organisms due to their clinical relevance or scientific utility (*Homo sapiens*, *Rattus norvegicus*, *Mus musculus*, *Gallus gallus*, *Danio rerio*, *Drosophila melanogaster*, *Caenorhabditis elegans*), for which we downloaded all sequences from the UniProt database [27], including Swiss-Prot (curated) and TrEMBL (predicted) [32] entries. Sequences of proteins in the TCDB have been aligned within each SLC-like family and subfamily and the alignments converted to HMMs for sensitive sequence similarity searches (see Methods). In addition, HMMs of SLC-like Pfam families were downloaded from the Pfam database. HMM-based similarity searches were then performed on the sequences downloaded for the 7 organisms to find proteins similar to any of the SLC-like TCDB families or subfamilies, or SLC-like Pfam domains, followed by the clustering of sequence fragments to arrive at one representative protein sequence per gene (see Methods).

The results of our search for SLC-like proteins are summarized in numbers in Table 2. Briefly, 59-67 of the 166 TCDB families have representatives in the 7 organisms (66 in human), and the organisms seem to contain 434-673 SLC-like proteins (549 in human). In total, 3733 proteins have been found in the 7 organisms studied. Notably, the number of SLC-like transporters found in human in our search is ∼130 higher than previously reported [2], indicating that the human SLC-ome may be significantly larger than previously thought.

**Table 2.** Results of the initial search for “SLC-like” proteins. The search was performed using HMM-based sequence similarity analysis based on selected families and subfamilies from the TCDB, as well as selected Pfam models that likely encode transmembrane domains of “SLC-like” transporters (see text for details). The table shows the number of families, subfamilies from the TCDB and the number of Pfam models that had representative sequences in each organism. In addition, the initial number of “SLC-like” proteins found is shown.

| Species | Level 3 families<br>(TC# x.y.z) | Level 4 subfamilies<br>(TC# x.y.z.w) | Pfam models | Proteins |
| --- | --- | --- | --- | --- |
| Human (*H. sapiens*) | 66 | 308 | 79 | 549 |
| Mouse (*M. musculus*) | 67 | 314 | 79 | 532 |
| Rat (*R. norvegicus*) | 67 | 316 | 79 | 537 |
| Zebrafish (*D. rerio*) | 65 | 312 | 80 | 673 |
| Chicken (*G. gallus*) | 63 | 305 | 77 | 473 |
| Fruitfly (*D. melanogaster*) | 60 | 273 | 66 | 434 |
| Roundworm (*C. elegans*) | 59 | 283 | 69 | 535 |

After arriving at this initial set of SLC-like proteins, we proceeded by a performing a first round of sanitization based on human proteins. First, we manually investigated the UniProt records of all human hits that are non Swiss-Prot-validated sequences, which revealed that 19 out of 23 are likely non-human or fragment sequences (S2 Table). After excluding these, we investigated the number of helical transmembrane (TM) segments annotated in UniProt for 144 of all remaining 530 human hits that do not correspond to classic SLC transporters (families SLC1-SLC52). In total, 17 of these proteins, showing similarity to 8 different TCDB families, contained less than 3 TMHs based on UniProt annotations and thus did not satisfy our structural criterion (1). These proteins have been investigated individually based on their coverage of HMMs used in the search and the location of TMHs according to UniProt annotations in other SLC-like proteins covering the same HMMs, as well as available literature on oligomeric state and function. This way, we have retained 8 sequences where the functional unit contains 3 or more TMHs, or the protein spans at least 3 TMHs based on HMM coverage, or for other reasons (see S4 Table). These proteins have been included in Table 3 and S4 Table. In contrast, 10 of the 17 proteins seemed to cover regions of HMMs where other members did not seem to contain TMHs, while in addition containing less than 3 TMHs or known non-transporter TM domains. Notably, certain Pfam models seem to cover both TM and non-TM domains, and several TCDB families and their corresponding HMMs encode multi-domain proteins, leading to hits that show similarity to the model in the non-TM region, but not in the TM region that would be crucial for SLC-likeness. These 10 proteins have been excluded from further analyses and included in S2 Table. Furthermore, initial SLC-like hits that only match with such non-TM regions of these HMMs (#2.A.19.3 positions 250-725, #9.B.64.1 positions above 200, “OST3_OST6” positions below 200, “RBP_receptor” positions above 400) in all other organisms have also been excluded from further analyses. Thus, in this initial sanitization step, 29 human proteins, as well as 8, 10, 2, 9, and 6 proteins from *M. musculus*, *R. norvegicus*, *C. elegans*, *D. rerio*, and *G. gallus*, respectively, have been excluded, leaving a remaining 3669 proteins for further analysis, including 520 from human.

**Table 3.**
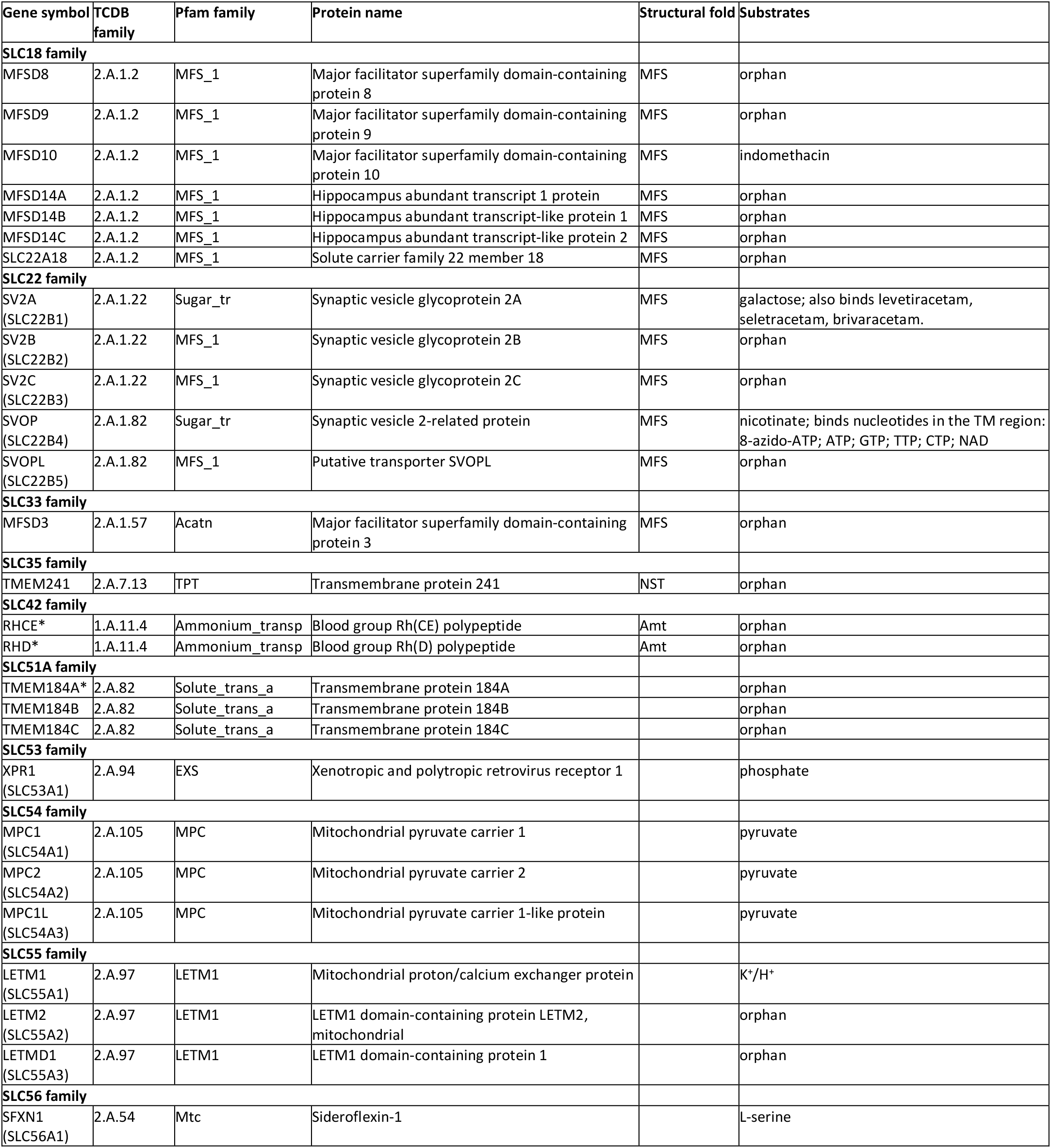
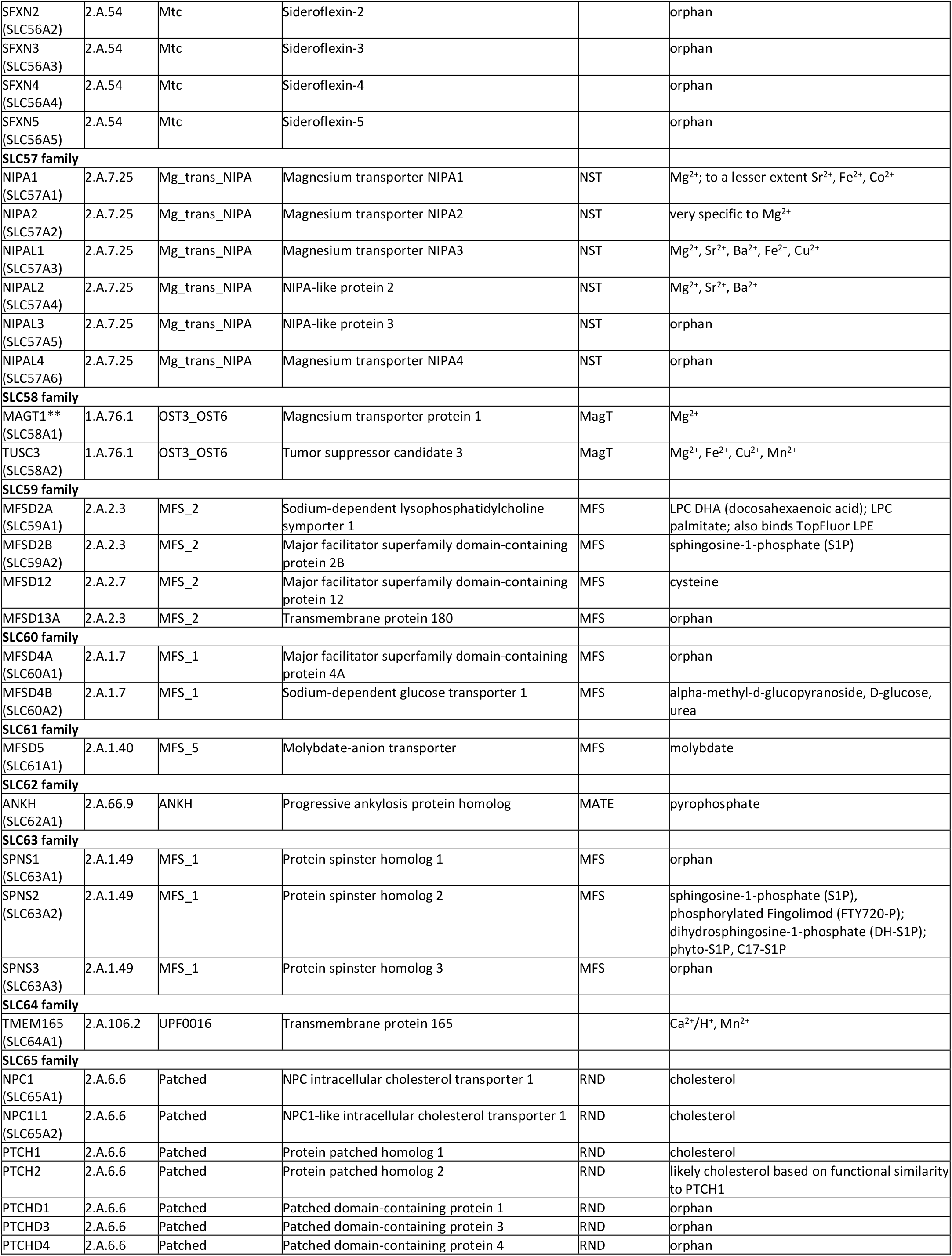
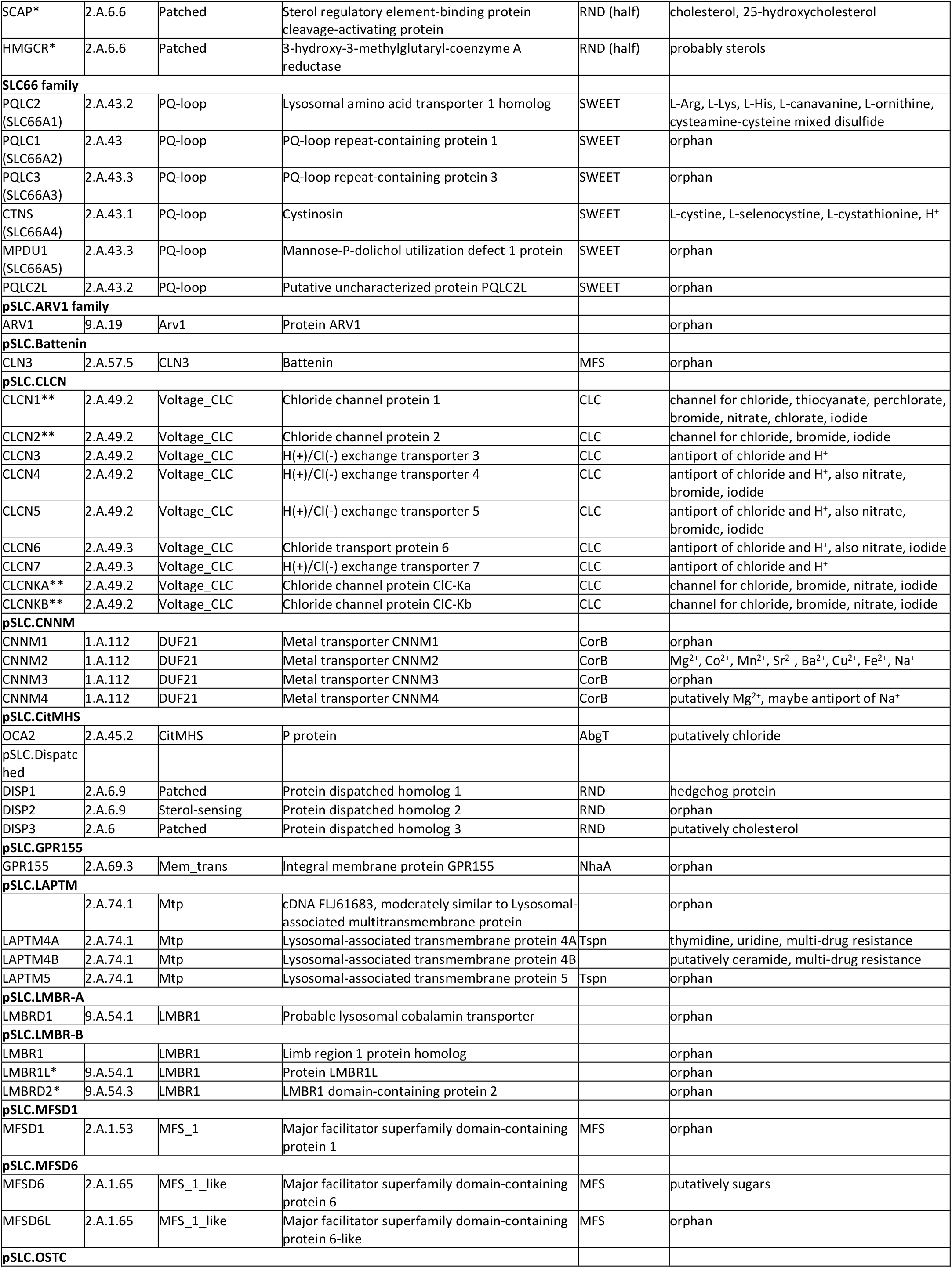
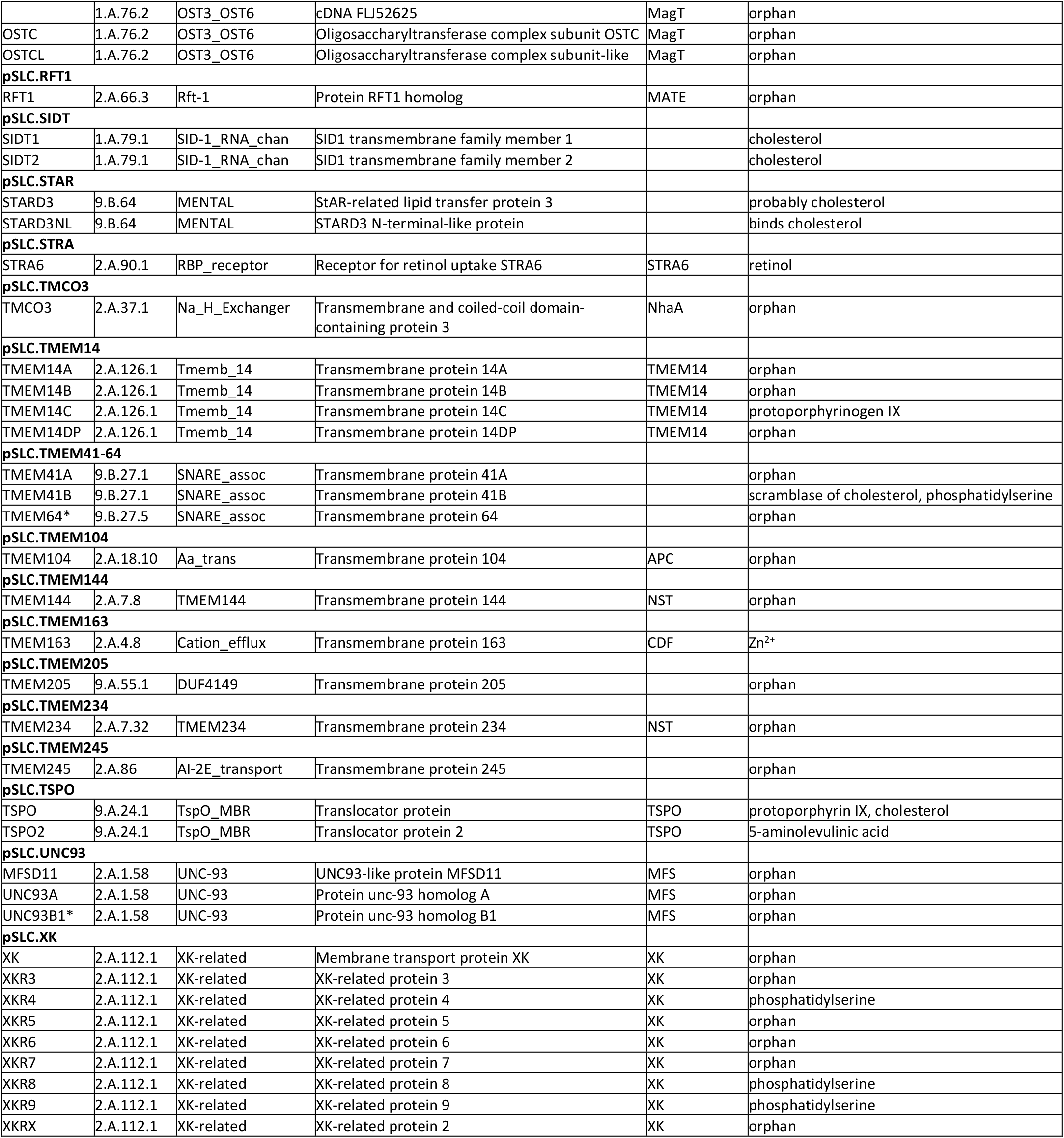
Novel SLC-like proteins. The table shows SLC-like human proteins from our search that are not in the classic group of SLCs (SLC1-52 families). The most similar (highest-scoring) TCDB family/subfamily and Pfam family are shown, as well as the most similar (highest-scoring) protein with structural information based on the pdb70 dataset (see Methods). Substrate information, where available, was retrieved from available literature. Based on our search, SLC families SLC53-66 have recently been incorporated into the nomenclature in collaboration with the HUGO/HGNC. Proteins marked with a single asterisk are included due to sequence similarity but may have functions other than transporter. Proteins marked with double asterisks might have a channel-like transport mechanism. For additional details, see S4 Table.

As a next step, we proceeded by attempting to classify the remaining proteins into protein families, followed by manual database and literature searches focused on novel human proteins, as detailed below.

### Classification into families

As mentioned in the introduction, SLC carriers are likely of polyphyletic origin, and individual families can be so diverse that even sensitive sequence similarity-based methods may have difficulties grouping related SLCs [12]. In our experience, multiple sequence alignment-based methods were not able to cluster the identified SLC-like sequences and to reproduce known SLC families, so we devised a custom method for clustering distantly related sequences into proteins families based on the introduction of “HMM fingerprints”. An HMM fingerprint is a mathematical vector of numbers assigned to a protein sequence, where the numbers represent the similarity scores of that protein sequence against each of the TCDB families, subfamilies and Pfam families that we have selected to be SLC-like. Thus, two protein sequences that show a similar pattern in their HMM fingerprints indicate their similarity. The usefulness of an HMM fingerprint depends on a meaningful definition of HMMs used in the fingerprint, whereby we capitalize on the evolutionary principle in the construction of TCDB families, subfamilies as well as Pfam families. Nevertheless, our goal was not to reconstruct the evolutionary history of a set of proteins, but to group proteins that share similar sequence features. However, due to the transitivity of homology and since similarity to a group of proteins suggests homology, clusters derived using HMM fingerprinting are likely to contain homologous proteins.

The HMM fingerprint-based classification of the SLC-like proteins found in our search yielded 102 protein families in total, 94 of which had representatives in human (Fig 1). For existing SLC transporters, the generated families corresponded well to classical SLC families. Interestingly, outlier proteins were found in several families, which did not cluster with their families at the threshold we used. Examples include SLC5A7, SLC10A7, SLC25A46, SLC30A9, SLC39A9, MPDU1/SLC66A5 as well as the SLC9B family and SLC35 subfamilies. This shows that the HMM fingerprints, and thus likely the sequences of these outlier proteins diverge from those of other members of their families, and the sequences of subfamilies seem to diverge in certain cases. Several classic SLC families also clustered together, such as SLC32-SLC36-SLC38, SLC2-SLC22, and SLC17-SLC18-SLC37 proteins, likely due to their high sequence similarity. To ensure optimal correlation with the preexisting classification of classic SLC proteins, we have introduced split and join constraints to keep 1) closely related families separated, 2) outlier proteins merged with their families and 3) heterogenous families merged (see Methods). These constraints did not affect the classification of novel SLC-like proteins. The number of families proposed by our method is somewhat dependent on the threshold of clustering used. Raising it to 0.8 lowers the number of families to 98 (90 in human), joining families SLC58-pSLC.OSTC, SLC8-SLC24, SLC45-SLC59, and SLC65-pSLC.Dispatched, as well as pSLC.TSPO to a 2-membered subfamily of unannotated proteins from *D. rerio*. Since our clustering method is based on shared HMMs, these families are likely related. Indeed, SLC8 and SLC24 proteins constitute a superfamily of Na^+^/Ca^2+^ antiporters [33], while NPC1/SLC65A1 and Dispatched proteins share structural similarity [34]. Lowering the clustering threshold to 0.6 generates 107 families (96 in human), splitting the pSLC.TMEM41-64 family to the TMEM41 and TMEM64 subfamilies, the MFSD3 proteins off the SLC33 family, an unannotated protein from *D. melanogaster* off the XK protein family, as well as splitting an unannotated protein family from *D. melanogaster* into two subfamilies.

**Figure 1.**
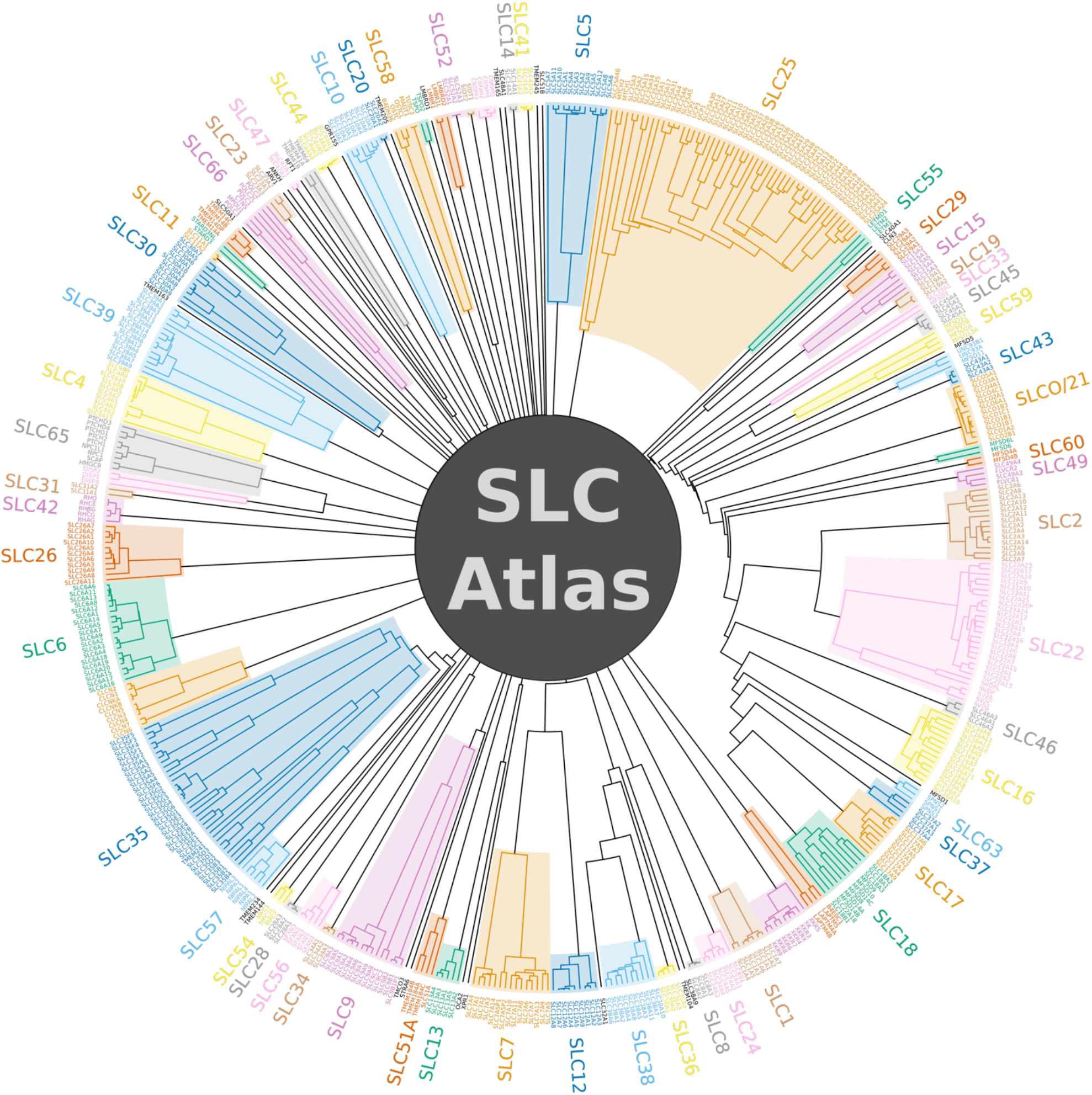
Dendrogram of human SLC-like protein families based on HMM fingerprint-based clustering. Classic and newly incorporated SLC families are shown with colors. Since SLC proteins are polyphyletic, which manifests itself in mathematically orthogonal HMM fingerprints that cannot be further clustered, the dendrogram does not join into a single branch. Branch lengths have been transformed for better visibility (see Methods).

Our analysis revealed 42 new proteins families in total in human, containing proteins that have not yet been annotated as SLCs. Interestingly, our search has also found new proteins that clustered into existing SLC families (Table 3, S4 Table).

### Structural homologues

For polytopic transmembrane proteins, structural similarity can support evolutionary relatedness, while at the same time evolutionary relatedness can provide a basis for homology-based model building efforts [35,36]. On the other hand, the lack of predicted similarity to any protein of known structure could pin down interesting targets for structural biology efforts by highlighting proteins likely belonging to new fold families.

We performed HMM-based searches on the pdb70 database (sequences of the proteins represented in the Protein Data Bank clustered to 70% sequence identity, see Methods) to assess whether structural homologues are available for the proteins found. In total, for 79 of the 102 families, at least one similar protein was found with a corresponding structure in the Protein Data Bank. Importantly, 477 human SLC-like proteins likely have a homologue whose structure has been solved. On the other hand, 43 human SLC-like proteins belonging to 19 different families do not seem to have homologues with a known structure and thus are likely to constitute novel fold families. For the classical SLC proteins, their best-scoring similar proteins from the pdb70 dataset and the corresponding structural fold families are summarized in S3 Table. Based on this, it appears that classical SLCs from families SLC34, SLC44, SLC48 and SLC51 are still “structural orphans”.

Detecting remote homology to proteins with a known transporter fold can also support their transporter function as well as give clues about their transport mechanism. Fold families that are well-characterized and typically host transporters with an alternating access mechanism encode potential transporter-like structures. Out of the 18 fold families to which novel SLC-like proteins show similarity to (S4 Table), representatives of 7 have been observed in various conformational states that hint at an alternating access mechanism, while for a further 7 fold families, a transport mechanism has been suggested (S4 Table). These fold families cover 87 of the novel SLC-like transporters, suggesting a possible transporter-like transport mechanism for these proteins (S4 Table).

### Phylogenetic analysis

Model organisms can be useful to study the biological function of various proteins, including solute transporters. In order to relate the results to human, however, knowledge of orthologous gene pairs is necessary. To this end, we performed phylogenetic analysis on each family of SLC-like proteins corresponding to our clustering. In brief, unrooted phylogenetic trees for each SLC-like protein family from all organisms have been generated and reconciled with the species tree of the 7 organisms in our study to identify gene duplication and speciation events in their evolutionary histories (see Methods). The resulting evolutionary trees are deposited in S1 File. Based on these trees, we carried out orthology analysis focusing on human proteins and the human lineage. The resulting data is presented in Fig 2, showing relationships between human genes and their orthologs in the other 6 organisms in our study. In addition, gene clusters are presented that have arisen through gene duplication events in the evolutionary history of the human lineage, but where the corresponding human genes have likely been lost.

**Figure 2.**
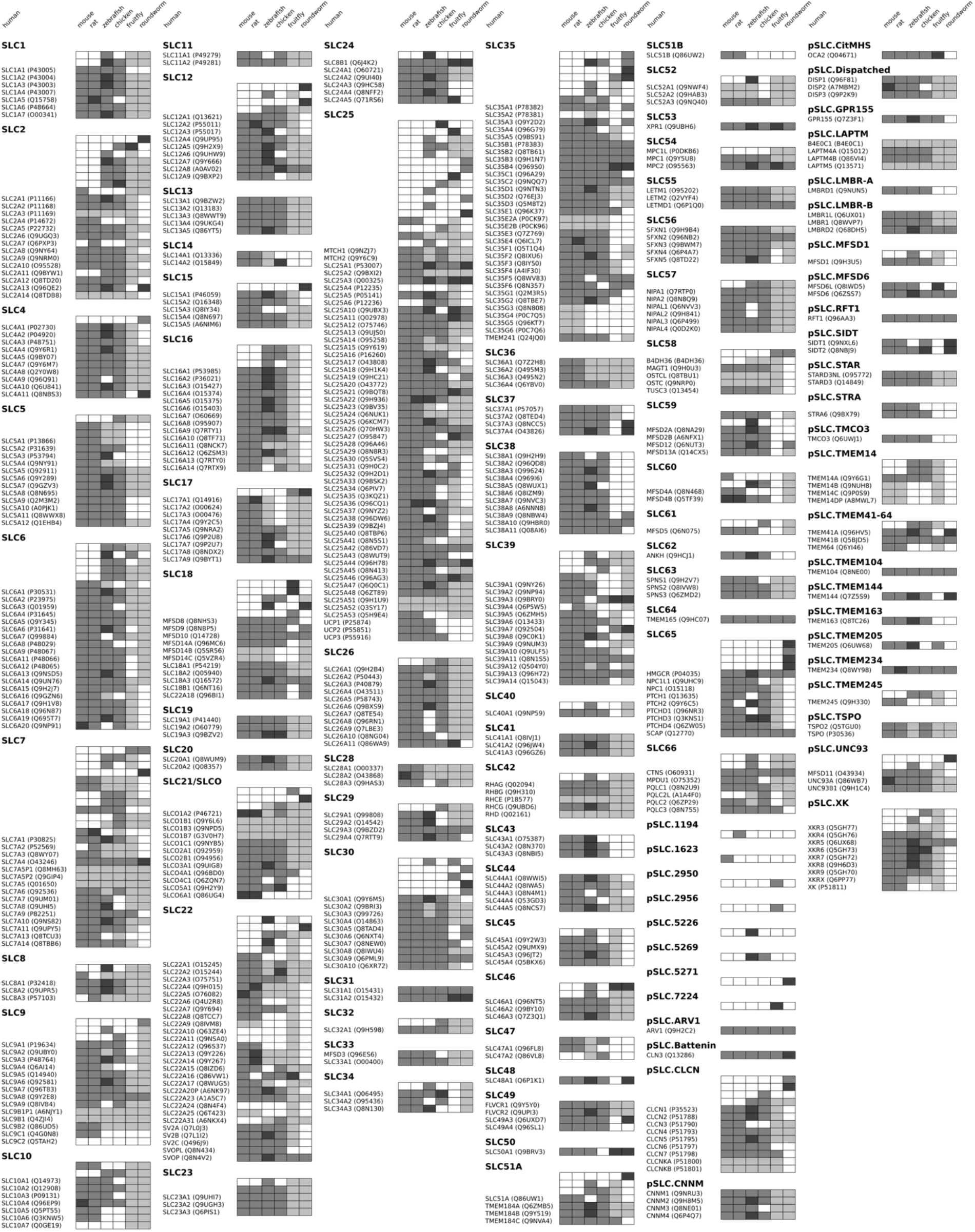
Orthology analysis of SLC-like proteins based on the human lineage. Each line shows a gene cluster that has evolved through gene duplication events along the human lineage. The human gene in the cluster, if present, is noted in the text label. Normal grey boxes in each column indicate a 1:1 orthology relationship between the human gene and a corresponding gene in the organism specified. Dark grey boxes indicate that the human gene has several orthologs, and light grey boxes indicate that several human genes share a common ortholog in that organism. White boxes denote no ortholog in the organism specified, while lines without a human gene name correspond to genes that have been lost in human.

### Literature search on newly found SLC-like proteins

As alluded to above, our search was followed by thorough investigation of the available literature to check whether a description of transport activity for the newly found proteins is available. Surprisingly, our search has revealed human proteins for which transported substrates are known. These data have been included in Table 3. While examining the available literature of these proteins, we have also extracted information related to our criteria for SLC-likeness, especially with a focus on structure, function and transport mechanism. In total, 53 proteins could be assigned a small-molecule substrate and therefore fulfill our criterion for substrate size (2), and for a further 6, putative small-molecule substrates have been suggested. Nevertheless, 74 proteins, while showing sequence similarity to transporters, are still “orphans” with no indication of a substrate. We have not found any indication in the literature that any of the proteins found in our search would have nucleoside-driven transport activity, therefore all of these proteins likely also fulfill our criterion 4. Interestingly, 9 proteins have been found that have described roles other than transmembrane solute transport, while showing similarity to SLC-like transporter proteins. These proteins potentially do not fulfill our functional criteria (5, 6, 7), they are marked with an asterisk in Table 3 and the details are further evaluated in S4 Table. In total, 3669 potentially SLC-like proteins have been found, and out of the 520 human proteins, 134 proteins that are not classic SLC transporters showed similarity to SLC-like proteins. Ruling out proteins with suggested other functions and a channel-like mechanism leaves 119 proteins, out of which 79 show similarity to transporters with a proposed transport mechanism, while 47 could be assigned endogenous small-molecule substrates.

## Discussion

Our curation and search efforts have revealed a surprising 134 human proteins that are SLC-like but have not been officially part of the SLC nomenclature before the start of our study. Interestingly, around 30 of them were addressed in an earlier study and referred to as “atypical SLC transporters” [10,11]. All of these atypical transporters have also been identified in our study, together with many others, including several that are not part of the major facilitator superfamily (MFS) or the amino acid-polyamine-cation (APC) transporter superfamily.

Our HMM fingerprint-based classification method yielded 102 protein families. Interestingly, in order to reproduce some of the classic SLC families, we had to introduce clustering constraints to artificially split or merge predicted families. In turn, increasing or decreasing the clustering threshold by 0.1 changes the number of families to 98 and 107, respectively. Researchers have reported similar problems with the classification of the highly diverse protein set of the Pfam database [29]. In particular, Finn et al. note that closely related Pfam families may have artificially high thresholds to prevent them from overlapping, while divergent families cannot always be covered by a single model [29]. Thus, we believe that however protein families are shaped, the discovery of evolutionary relationships between them will always be necessary to define higher-order superfamilies, similarly to Pfam clans [29]. Similarly, the analysis of the evolutionary relationship of proteins within each family is equally validated, and several classic SLC families indeed contain established subfamilies based on sequence and/or functional comparisons [37–40]. In this regard, we believe that our work provides correlation between transporter proteins in the studied organisms with the hierarchy of families and subfamilies in the TCDB database on the one hand, and a scale-free framework for protein grouping and classification through HMM fingerprints on the other hand. Still, more work is needed to examine the evolutionary relatedness of different SLC-like families using more robust phylogenetic approaches.

The phylogenetic trees of SLC-like protein families presented in S1 File can be instrumental for functional annotation based on orthology analysis in lower organisms, as well for studying species-specific differences in transport pathways. Of particular note, different orthologs of hepatic drug transporters are present in humans and experimental mammals used in pre-clinical studies, contributing to imperfect prediction of drug half-life and toxicity in animal models [41]. On the other hand, genetic ablation of transporters and phenotype studies in lower organisms could shed light on the function of their human orthologs. However, as can be seen in Fig 2 and the phylogenetic trees in S1 File, a significant number of SLC-like protein sequences were found in *D. melanogaster* and *C. elegans* that do not appear to have orthologs in the higher organisms studied here. Most of these protein sequences are poorly annotated and their expression and function have yet to be confirmed.

The interesting cases of apparent outlier sequences in several families (SLC5A7, SLC10A7, SLC25A46, SLC30A9, SLC39A9, MPDU1/SLC66A5 as well as the SLC9B and SLC35 families) have previously been partially documented. These members show striking sequence and/or functional divergence from other members of their SLC families. SLC5A7 diverges from other SLC5 proteins in phylogenetic trees [42,43] and shares only 20-25% sequence identity with them [44]. The sequence of SLC10A7 is more similar to its bacterial relatives than to other SLC10 members, and its genetic structure, particularly its number of exons, is also different from other proteins in the SLC10 family [45,46]. SLC10A7 also appears to have diverged at the functional level, as it has been suggested to be a regulator of Ca^2+^ influx [47], while having no documented transport activity. SLC25A46 turns out to be an outer mitochondrial membrane protein [48], in contrast to most other members of the SLC25 family, which are generally located on the inner mitochondrial membrane. Consistent with this, its yeast ortholog Ugo1, referred to as “a degenerate member of the mitochondrial metabolite carrier family”, was reported to be part of the mitochondrial outer membrane fusion machinery [49]. Similarly, MTCH1 and MTCH2, which also seem to be distantly related to other SLC25 members according to our dendrogram (Fig 1), are also outer mitochondrial membrane proteins [50,51], and together with SLC25A46, are collectively referred to as “peculiar” members of the family [52]. SLC30A9 is found in the cytosol and nuclear fractions and functions as a coactivator of nuclear receptors after hormonal stimulation [53], in contrast to other members of the family, which are Zn^2+^ transporters. SLC39A9, the only member of “subfamily I” of Zrt/Irt-like proteins (ZIPs) [37] might function as an androgen hormone receptor [54], while most other family members are primarily transporters of divalent metal ions. SLC9B subfamily proteins NHA1 and NHA2 of the SLC9 Na^+^/H^+^ exchanger family are more similar to their bacterial homologs than to SLC9A members [55].

In the following sections, we discuss the non-classic set of human SLC-like proteins resulting from our search, with a particular focus on the identities of the transported substrates and, if known, the structural and mechanistic aspects of transport. In certain cases, the proteins have already been officially included in the SLC nomenclature as per our initiative and as a result of the precursor of this work, following approval by the Human Gene Nomenclature Committee (HGNC). Where existing information about substrate and function is not available, we have speculated on these aspects using available information in the literature and considerations of sequence similarity. For several proteins, however, their classification requires further consideration. We would also like to articulate open questions about what further work is needed in order to identify additional transporters. We believe our approach has been useful to pinpoint proteins that have a high probability of being novel transporters. Thus, specific biological, biochemical and structural efforts could focus on these specific targets highlighted in our work, all of which would contribute to a complete assessment of the SLC-ome in human cells.

### Transporters with existing evidence for transport function

Importantly, our search highlighted several proteins for which our literature search uncovered previously reported evidence of transporter activity. Several of the proteins in these families have been assigned SLC family numbers and, in collaboration with the HGNC, included in the SLC nomenclature. Among these hits are several mitochondrial transporters (MPC/SLC54, LETM/SLC55, Sideroflexins/SLC56), which have been reviewed before [56]. In terms of transported substrate, many of the proteins with documented transport activity appear to be ion transporters or exchangers (Table 3). In the next paragraph, we will highlight certain proteins and families that have particularly caught our attention.

Interestingly, the **SLC60** family contains two MFS-like proteins (MFSD4A, MFSD4B), of which MFSD4B has been shown to transport D-glucose and urea [57,58].

The **SLC61A1** protein (MFSD5) is the only protein in human that shows similarity to the #2.A.1.40 family of molybdate transporters and contains the “MFS_5” Pfam model. It has been claimed to be the homolog of similar transporters from algae and plants, and complementation assays suggested its ability to transport molybdate [59]. While molybdenum is a biologically active trace element, not much is known about its transport and homeostasis in human [60].

Interestingly, the **TMEM163** protein clustered together with SLC30 zinc transporter (ZnT) proteins, since the “Cation_efflux” Pfam model, representative of the SLC30 family, was present in its sequence, although at a low score and non-significant e-value (2.9e-3). In the TCDB, TMEM163 is also classified under subfamily #2.A.4.8, sharing a common family with SLC30 transporters (#2.A.4.2). Multiple sequence alignment as well as pairwise alignments with existing SLC30 members reveal very low sequence identity with SLC30 proteins (4.2-14.4%), albeit these numbers are similar to those of SLC30A9 (6.5-13.4%). Given the marginal similarity to the “Cation_efflux” domain, it is tempting to assume that SLC30 proteins and TMEM163 are distantly related. Indeed, TMEM163 has been shown to bind [61] and transport Zn^2+^ [62–64], and substitution of its proposed substrate-binding residues with alanine abolished Zn^2+^ efflux activity [64]. Transport has been demonstrated to be H^+^-coupled, and the protein functioning as a dimer [62], while extruding Zn^2+^ from the cell [64]. Intracellularly, TMEM163 was originally shown to be expressed in synaptic vesicles [65]. In overexpression systems, it is localized to both the plasma membrane and intracellular membrane compartments [64]. TMEM163 has been linked to Parkinson’s disease (PD) [66], even though the opposite conclusion has also been drawn [67]. TMEM163 has also been reported to be upregulated by olanzapine, a psychotropic drug prescribed for PD patients [68]. In addition, TMEM163 was also shown to be highly expressed in insulin secretory vesicles in human pancreas [69], and has been identified as a risk factor in type 2 diabetes [70,71]. Disruption of TMEM163 expression might impair insulin secretion at high glucose stimuli [69].

The **TMEM165** protein clustered into its own family and is the only protein in human containing the “UPF0016” Pfam domain and showing similarity to TCDB family #2.A.106.2. TMEM165 is a member of a highly conserved family of transmembrane proteins that is present in many species of eukaryotes and bacteria [72]. Initially, TMEM165 and its yeast homolog, Gdt1p, have been hypothesized to be Ca^2+^/H^+^ exchangers [73,74]. However, recently, evidence has been mounting about its involvement in manganese (Mn^2+^) homeostasis [74], and both Ca^2+^ and Mn^2+^ transport activity has directly been shown [75]. TMEM165 is localized to the trans-Golgi in human cells [72], and is proposed to play a crucial role in regulating Mn^2+^ uptake into the Golgi apparatus [72,74]. In line with this, its homologs in other organisms, also containing the UPF0016 domain, are also annotated as Mn^2+^ transporters [74]. Manganese plays an important role as a co-factor for enzymes involved in glycosylation, and impairment of TMEM165 function results in glycosylation defects. Indeed, mutations of TMEM165 found in patients with congenital disorder of glycosylation (CDG) type II hamper the transport function or localization of TMEM165 [75]. Due to the importance of TMEM165 in lactate biosynthesis [76], it has also been suggested that TMEM165 could be a transporter importing both Ca^2+^ [77] and Mn^2+^ into the Golgi in exchange for protons [74]. TMEM165 proteins contain two copies of the UPF0016 domain, and each domain contains a signature motif, E-<π-G-D-(K/R)-(T/S), where <π denotes a hydrophobic amino acid. The glutamic acid of the second motif, E248, has been shown to be crucial for affecting the glycosylation function of the Golgi but not the expression of the protein [78], and so can be speculated to form part of a binding site for transporter function. However, in the absence of an experimentally determined structure, further investigation will be required to understand the transport mechanism of TMEM165.

### Proteins with sequence similarity to existing transporters

Our search uncovered a large number of proteins that show sequence similarity and thus possible relationships to existing transporters in the SLC nomenclature. Since transport activity has not been demonstrated, these proteins are either orphan transporters or they could have transceptor functions. What follows is a comprehensive discussion of these proteins, as their similarity to transporters makes them ideal targets for further studies to elucidate their putative transporter activity.

#### Atypical transporters

A previous effort by Perland and coworkers has uncovered novel transporter-like proteins mostly from the MFS and APC superfamilies [11], which have also been recognized by our search. In general, the function of these atypical transporters is not well known, but some have been reported to be expressed in the brain, and their expression levels seem to be affected by nutrient availability [79–82]. For MFSD1, MFSD6 and UNC93A, the study of the *D. melanogaster* and *C. elegans* orthologs have provided some information on the loss-of-function phenotype [83–86]. MFSD8 and MFSD10 have been linked to the Wolf-Hirschhorn syndrome and to LINCL (late-infantile-onset neuronal ceroid lipofuscinoses), respectively [87,88], and MFSD8 seems to be localized in the lysosomes [89,90]. More studies about the biological function and transport activity of these proteins is required to fully understand their physiological roles.

Some of the “atypical” SLC-like transporters (e.g. MFSD8, MFSD9, MFSD10 and MFSD14 proteins) clustered together with members of the classical SLC18 family, which prompted us to examine the relationship of these and neighboring proteins in more detail. We constructed multiple alignments and a phylogenetic tree of the proteins one level above these proteins in our clustering dendrogram (i.e., members of the SLC17, SLC18, SLC37 families as well as MFSD8, MFSD9, MFSD10, MFSD14A-C and SLC22A18 proteins, Fig 3). As expected, the phylogenetic tree gives a better separation of these very similar proteins than the HMM fingerprint-based dendrogram, and the branch support values suggest a clear separation of the SLC17, SLC18 and SLC37 families. In addition, the phylogenetic tree highlights that the atypical SLC proteins MFSD8, MFSD9, MFSD10, MFSD14A-C as well as SLC22A18, while being more divergent, are likely to have evolved from a single common ancestor. The relationship between the MFSD9, MFSD10 and MFSD14A-B proteins also agrees with earlier studies [10,11]. Similarly, the evolutionary dendrogram created using all 7 organisms in our study for the SLC18 family (S1 File) suggests that MFSD9, MFSD10, MFSD14A-B and SLC22A18 likely share a common evolutionary origin and are thus more closely related to each other than to SLC18 proteins, while MFSD8 is more distantly related. Further studies may be required to elucidate the particular evolutionary relationship between these proteins.

**Figure 3.**
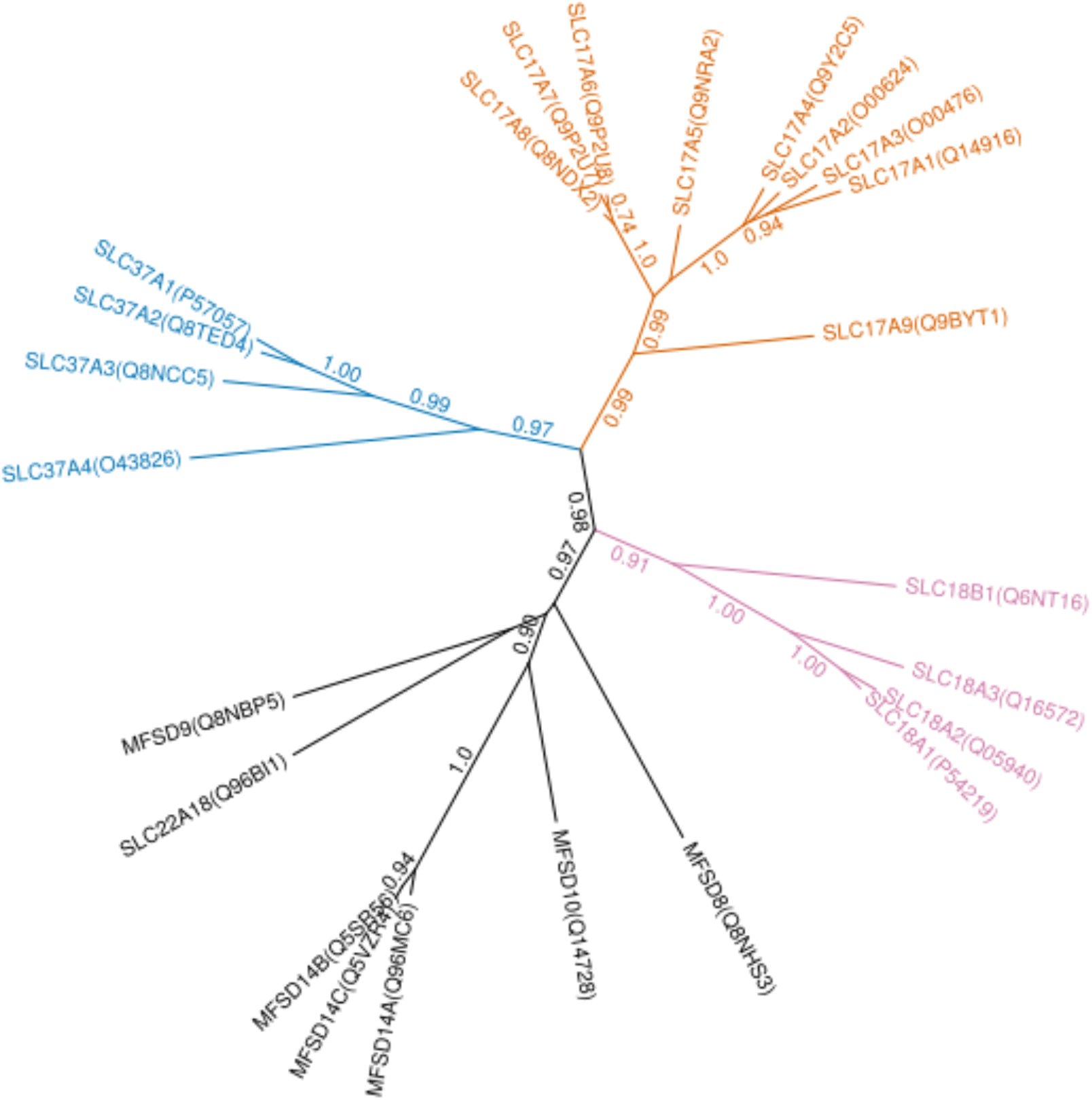
Phylogenetic tree of the SLC17, SLC18 and SLC37 families and proteins clustering in their neighborhood. Branch support values ≥ 0.7 are shown.

Interestingly, **MFSD3** has clustered together with the SLC33A1 protein in our HMM fingerprint-based clustering analysis. Indeed, the “Acatn” (Acetyl-coenzyme A transporter 1) Pfam domain is present in MFSD3, albeit with a relatively low score, but with significant e-value (5.6e-12). Sequence alignment between SLC33A1 and MFSD3 gives 18.2% sequence identity. Even though the sequence identity between MFSD3 and SLC33A1 is relatively low, the “Acatn” domain was found only in these proteins. The relatedness of MFSD3 and SLC33A1 is also corroborated by previous results of other groups [81]. The biological function of MFSD3 is still unclear [81,91,92].

The **TMEM104** protein in our analysis clustered together with amino acid transporter families SLC32, SLC36 and SLC38. Based on multiple and pairwise sequence alignments and sequence identity, TMEM104 was most similar to SLC38A7 (13.3-15.1%), SLC38A8 (13.1-15.1%), and SLC36A1 (10.9-16.1%). Interestingly, TMEM104 also bears moderate similarity to the “Aa_trans” Pfam domain, which describes the transmembrane region of SLC38 proteins. In our SLC classification dendrogram (Fig 1), TMEM104 clustered with SLC38 proteins, even though it seems to be an outlier from the family, similarly to SLC38A9. In addition, TMEM104, SLC38A7 and SLC38A8 all show low similarity to the “Trp_Tyr_perm” Pfam domain, which describes bacterial tyrosine and tryptophan permeases. Despite the low sequence similarity to SLC38 members, these data suggest that TMEM104 might be an amino acid transporter distantly related to the SLC38 family. To get a more detailed picture of the evolutionary relationship of TMEM104 and the SLC38, SLC36 and SLC32 families, we constructed a multiple alignment and a phylogenetic tree of these proteins (Fig 4). While the tree undoubtedly separates the SLC32 and SLC36 clades due to high branch support values, TMEM104 could not be clearly separated from the SLC38 family, and it likely has a similar relationship to the rest of the family as SLC38A9, which is playing a transceptor role in cells [93]. However, there is currently no experimental evidence for this and the biological function of TMEM104 remains elusive.

**Figure 4.**
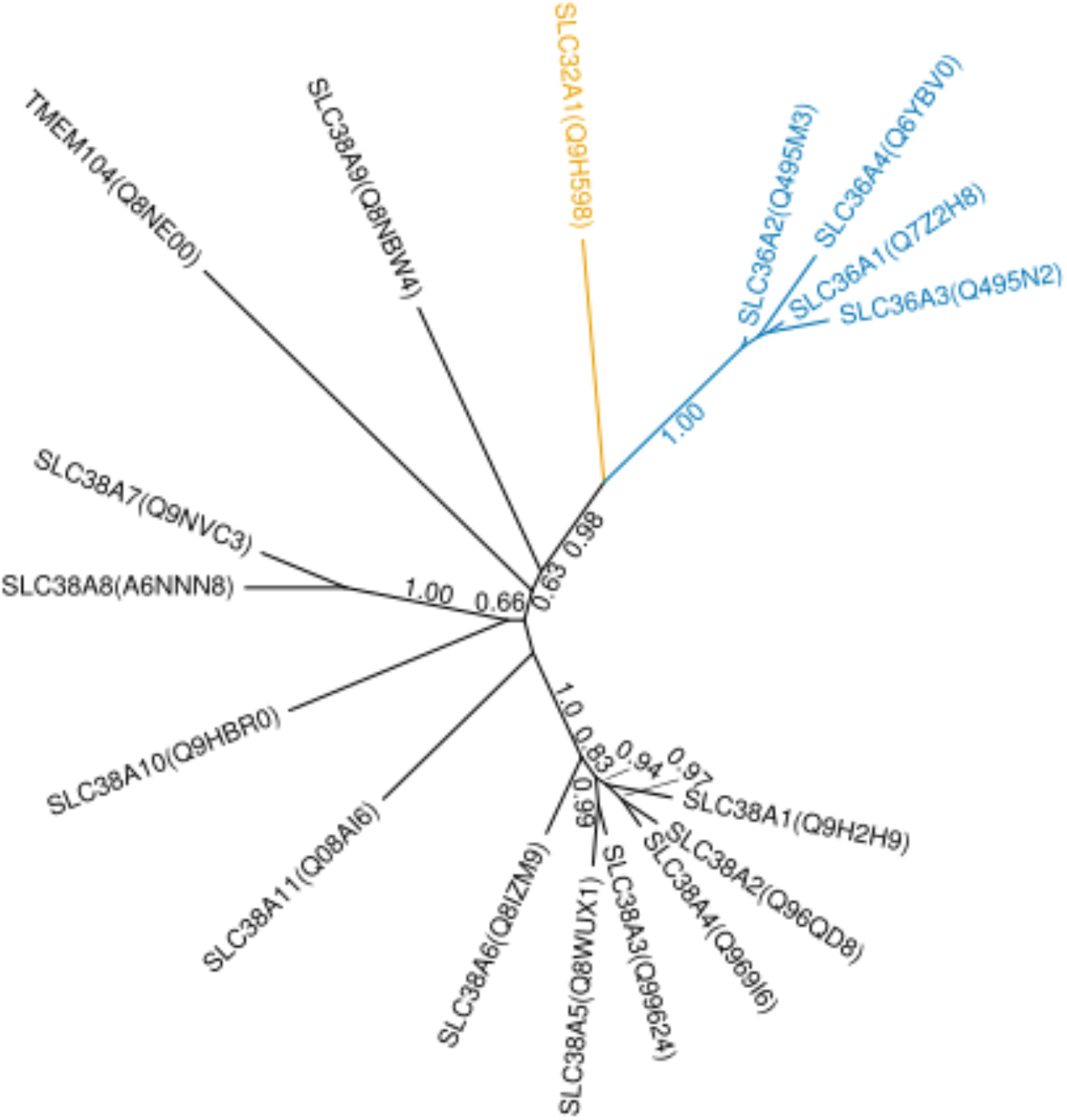
Phylogenetic tree of proteins clustering in the SLC38-36-32 families and TMEM104. Branch support values ≥ 0.5 are shown. SLC32 and SLC36 family members are colored gold and blue, respectively.

#### Proteins similar to SLC35 transporters

Interestingly, our search revealed several proteins that show sequence similarity to transporters of the SLC35 family. SLC35 proteins are currently classified into subfamilies A-G, which have relatively low sequence identity among them (4.0-22.0%). SLC35 transporters belong to the family of “DMT” (drug-metabolite transporters), which is classified in TCDB family #2.A.7, and corresponds to a clan of Pfam families, called DMT. Currently known substrates of human SLC35 members include nucleotide-sugar conjugates [40]. However, the substrate range of this superfamily is substantially larger [94].

The **TMEM144** proteins harbors its dedicated Pfam model called “TMEM144”, which itself is a member of the DMT clan of transporters. Its relatedness to the DMT family is further corroborated by high-scoring similarity of the TCDB subfamily #2.A.7.8 to the sequence of TMEM144. Otherwise, functionally, the protein is uncharacterized, although it might be related to sterol metabolism/transport, because its function has been linked to bovine milk cholesterol levels [95], the hypothalamic-gonadal axis and testosterone response [96]. It is also highly expressed in the hypothalamus [96].

**TMEM234** is classified in the TCDB #2.A.7.32 family and also contains a corresponding “TMEM234” Pfam domain, which is a member of the “DMT” clan of Pfam domains as well. The physiological role of TMEM234 is not known. However, in zebrafish, its homolog might play a role in the formation of the kidney filtration barrier, as its knockdown causes proteinurea [97].

In our clustering analysis, **TMEM241** clustered with the SLC35 family very closely. Its HMM fingerprint shows similarity to the TC# 2.A.7.13 subfamily, and weak similarity to the “TPT” Pfam model (which also belongs to the “DMT” Pfam clan) over the whole length of the protein. The proteins in the #2.A.7.13 family are Golgi GDP-mannose:GMP antiporters from plants, yeast and other organisms [98,99], but not from vertebrates. Nevertheless, the protein seems to be present in many higher organisms according to the Swiss-Prot database. However, these protein are not listed in the TCDB. The biological function of TMEM241 is still unknown, but it has been suggested to affect serum triglyceride levels [100].

Due to the sequence diversity of the SLC35 family, we were interested in the relationships between individual proteins. To this end, we have built a phylogenetic tree of human proteins that showed similarity to existing SLC35 transporters (Fig 5). In the tree, most SLC35 subfamilies could be resolved as a single clade, while TMEM241 and TMEM234 form clades with SLC35D and SLC35F3-5 proteins with a support value of 0.91 and 0.71, respectively. The relationship of TMEM241 with the SLC35D subfamily is also supported by our HMM fingerprint-based clustering results. TMEM241 shows 12.0-21.4% sequence identity with SLC35D proteins. In contrast, our phylogenetic tree with SLC35 proteins from all 7 organisms (S1 File) indicated that TMEM241 is most closely related to SLC35E4. On the other hand, TMEM234 only weakly associated with SLC35F proteins, with sequence identities 3.6-9.2%. TMEM144 appears to be only distantly related to SLC35 proteins. The elucidation of evolutionary relationships between proteins in the SLC35 thus likely requires further investigation.

**Figure 5.**
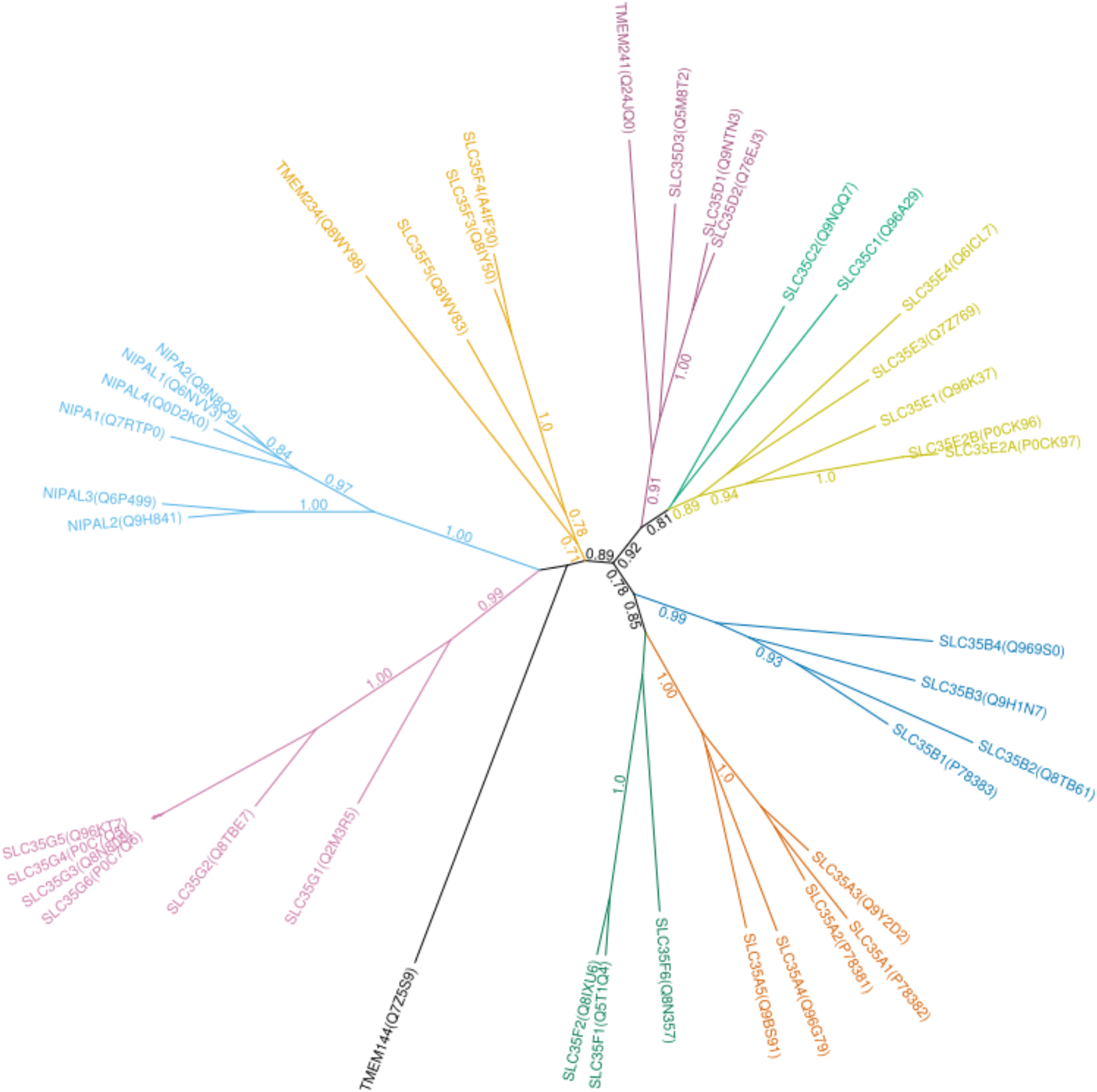
Phylogenetic tree of proteins with HMM fingerprints overlapping with the SLC35 family. Branch support values ≥ 0.7 are shown. The different subfamilies are colored in various colors.

#### Others

**GPR155** is an enigmatic protein that seems to be a concatenation of a membrane transporter domain (Pfam: “Mem_trans”) and a G-protein coupled receptor (GPCR) domain, which might be the reason why it is annotated as a GPCR. The membrane transporter part seems to be most similar to TCDB #2.A.69.3 subfamily proteins, which are annotated as malate/malonate transporters in the Auxin Efflux Carrier (AEC) family (#2.A.69). Gene structure analysis suggests that the concatenation is real [101], and both the human, mouse and fruitfly proteins seem to contain 17 TMHs according to UniProt annotations.

The “Mem_trans” domain is only present in GPR155 from all human proteins analyzed, and matches the first 10 TMHs of the protein in a 5+5 arrangement. In our structural search, the second half of this transporter domain (TMHs 6-10) of GPR155 exhibits similarity to the N-terminal half of sodium/bile transporters of the AsbT fold (SLC10 family). On the other hand, the last 7 TMHs of GPR155 (TMHs 11-17) indeed show similarity to GPCR-fold (7-TM) proteins with known structure, with highest similarity to structures of the human Smoothened receptor homolog (PDB ID: 6OT0). Interestingly, in our search, the N-terminal half of the transporter domain of GPR155 did not show any structural homologues.

The precise function of GPR155 still remains elusive. However, because highest expression levels were found in the brain, especially in GABAergic neurons, it might play a role in GABAergic neurotransmission [101]. It also has been suggested that GPR155 might play a role in neurons involved in motor brain function as well as sensory information processing [101]. In *D. melanogaster*, knockdown of the homologous gene, “anchor”, resulted in increased wing size and thickened veins [102]. This phenotype was similar to what appeared in bone morphogenetic protein (BMP) signaling gain-of-function experiments [102]. GPR155 has also been linked to a number of different cancers [103,104].

The **RFT1** protein was originally thought to be a scramblase of lipid-linked origosaccharides [105]. However, these molecules have at least 12-14 sugar moieties, so given their size, it is unlikely that a single transporter could catalyze their flipping. Later studies refuted the scramblase concept and suggested instead that RFT1 could serve as an accessory protein to a flippase, but would not act as a flippase itself [106–108].

Nevertheless, the corresponding “Rft1” Pfam model shows similarity to multidrug and toxic compound extrusion (MATE) transporters (SLC47 family) and belongs to a clan of Pfam domains (“MviN_MATE”) that contains transporters as well. In line with this, human RFT1 showed significant similarity to MATE transporters in our search for structural homologs, indicating likely structural similarity. Some members of the corresponding TC# 2.A.66.3 subfamily also contain weak hits of the “MatE” Pfam domain. Thus, while RFT1 shows similarity to existing transporters, its biological function is still unclear.

The C-terminal half of the **TMEM245** protein shows weak similarity to TC# 2.A.86 proteins (Autoinducer-2 Exporter/AI-2E family), which contain both small-molecule exporters [109,110] as well as Na^+^/H^+^ antiporter proteins [111–113]. Accordingly, TMEM245 also has weak similarity to the corresponding Pfam model (“AI-2E_transport”).

The HMMs of TC# 2.A.86 and #2.A.86.1 match residues 444-866 of human TMEM245, which are the last 6 TMHs according to UniProt predictions. The last 5 TMHs are separated from the previous ones by a slightly larger loop. This architecture is similar to the 3+5 arrangement of the previously described bacterial Na^+^(Li^+^)/H^+^ antiporter TC# 2.A.86.1.14 according to UniProt predictions (accession code: NLHAP_HALAA). This bacterial protein also matched the full-length “AI-2E_transport” domain from Pfam, while only the last 5 TMHs of TMEM245 match with C-terminal region of “AI-2E_transport”. The human TMEM245 protein contains 14 TMHs in total according to UniProt predictions. Thus, TMEM245 might have a transporter-like domain at the C-terminus. In terms of the structure, we have not found any similarities to proteins with known structure. Therefore both TMEM245 and bacterial exporters and antiporters in the TC# 2.A.86 family are likely to have a yet uncharacterized tertiary structure. Functionally, the TMEM245 protein also remains elusive.

The **TMEM41A**, **TMEM41B**, and **TMEM64** proteins clustered to the same family in our results. These are the only proteins in human that show any similarity to the “SNARE_assoc” Pfam domain, as well as to the TCDB family #9.B.27. While no protein with this domain or from this family has direct evidence for transport activity, Pfam reports SCOOP-based similarity [31] of the “SNARE_assoc” domain with “Sm_multidrug_ex”, which is a domain encoding transporter proteins. Some members of the family in the TCDB have been proposed to be “cation:proton importers” (#9.B.27.2.2) or “selenite transport proteins” (#9.B.27.2.3).

The most characterized member of the human protein family is TMEM41B. Interestingly, a recent study reported a putative structure generated *ab initio* using evolutionary covariance-derived information [114].

Strikingly, this structural model shows features reminiscent of secondary transporters, such as a tandem internal repeat with two-fold rotational symmetry, and the authors suggest a H^+^ antiporter activity as a mechanism of transport [114].

While the exact function of TMEM41B is still unclear, it forms a complex with vacuole membrane protein 1 (VMP1), also harboring the “SNARE_assoc” domain, and both are required for autophagosome formation [115,116]. Tmem41b localized to mitochondria-associated ER membranes [117–119]. Interestingly, TMEM41B seems to be an absolutely required factor for SARS-CoV-2 [120], and probably also flaviviral [121] infection, possibly by facilitating a membrane curvature that is beneficial for viral replication [121].

The proteins **TMEM184A**, **TMEM184B**, **TMEM184C** clustered together with **SLC51A** (family of transporters of steroid-derived molecules) in our analysis. While human TMEM184B and SLC51A are included in the TCDB as members of family #2.A.82, TMEM184A and C are not. Independently, the “Solute_trans_a” Pfam model was found in all four proteins with high scores and significance, but not in other human proteins. Therefore, it is likely that the four proteins, TMEM184A-C and SLC51A, are homologous. In spite of this, sequence identity between TMEM184 proteins and SLC51A is low (12.3%-13.6%), but moderate among TMEM184 proteins (26.5%-62.0%). All four proteins are predicted to harbor 7 TM helices according to UniProt, yet our search has found no similar proteins with known structures.

TMEM184A was identified as a heparin receptor in vascular cells [122], but no transport activity has been reported. Interestingly, while SLC51A is known to function as a bile acid transporter [123–125], TMEM184B has been proposed to be responsible for ibuprofen uptake [126]. This is interesting in view of the partial chemical similarity between steroid acids and ibuprofen, both harboring a hydrophobic hydrocarbon part and a carboxyl moiety. TMEM184C resides in a genetic locus that has been suggested to be responsible for the pathogenicity of X-linked congenital hypertrichosis syndrome [127], but no transport activity has been suggested.

#### Putative transporters

Our search also identified proteins whose transport activity is either controversial or not characterized, and which do not show sequence similarity to transporters of known function. Thus, the proteins in these families require further investigation to uncover their function.

The **CNNM1-4** proteins (also called ACDP1-4) are distant homologs of the cyclins, but have no documented enzymatic activity. Instead, CNNM proteins belong to a highly conserved family of Mg^2+^ transport-related proteins [128], and CNNM2 and CNNM4 have been proposed to be the long sought after basolateral Na^+^/Mg^2+^ exchangers in the kidney and intestine, respectively [129,130]. The function of these proteins is, however, controversial [131–134], and there are hypotheses that CNNM proteins *per se* are not Mg^2+^ transporters [135]. Most recently, however, the structure of a bacterial homolog, CorC, has been resolved, revealing its membrane topology, as well as a conserved Mg^2+^-binding site [136]. Strikingly, the Mg^2+^ ions in the structure are fully dehydrated, in contrast to those in other known Mg^2+^ channel structures [136], which makes it unlikely that the proteins function via a channel-like transport mechanism. In line with this, the authors suggest an alternating-access exchange mechanism [136], however, further studies are required to understand how and whether CNNM proteins might be able to mediate the translocation of Mg^2+^ ions across the membrane.

A family of 4 lysosomal-associated transmembrane proteins (**LAPTM4A**, **LAPTM4B**, **LAPTM5** and sequence **B4E0C1**) turned up in our search, corresponding to the TCDB subfamily #2.A.74.1 and Pfam model “Mtp” (mouse transporter protein). The family also includes an uncharacterized transcript with the UniProt accession “B4E0C1”. Originally, the mouse transporter protein (Mtp, ortholog of LAPTM4A) was characterized as a transporter mediating the transport of nucleosides and nucleobases between the cytoplasm and intracellular compartments [137], and was later also associated with multidrug-resistance (MDR) in yeast, where its expression changed the subcellular compartmentalization of a heterogenous group of compounds [138,139]. LAPTM4A was shown to be involved in glycosylation and glycolipid regulation [140,141]. All three proteins seem to be lysosomal [137,142,143]. Nevertheless, these proteins appear to interact with other characterized transporters, such as SLC22A2 (hOCT2), SLC7A5/SLC3A2 (LAT1/4F2hc) and MDR-related ABC transporters [144–146]. But it has been claimed that they are not *per se* transporters, but rather regulatory factors, either assisting the localization and targeting or the function of other transporters [146]. Interestingly, the transcript “B4E0C1” appears to have 4 TMHs at its N-terminus, which is identical to human LAPTM5 apart from a ∼40-amino acid insertion between TMH3 and TMH4. This region also shows significant similarity to both the “Mtp” Pfam model and the TC# 2.A.74.1 subfamily. However, the C-terminal region of the transcript is identical to the C-terminal segment of “actin filament-associated protein 1-like 1” protein (UniProt accession Q8TED9). We did not find any similar fusion sequences in the other organisms we analyzed. The “Mtp” Pfam domain, which is the hallmark of the family, belongs to the “Tetraspannin” Pfam clan, which has no other domains with annotated transporter function and no similarity to existing transporters. Structural information on “Tetraspannin” proteins is also not available. In summary, the transport function of these proteins requires further investigation.

Our search identified four proteins in human (**LMBR1**, **LMBR1L**, **LMBD1/LMBRD1**, **LMBRD2**) bearing the “LMBR1” Pfam domain, which clustered into two families in our analysis. These proteins correspond to TCDB family #9.A.54. The LMBD1 protein, encoded by the LMBRD1 gene, was suggested to function as a vitamin B12 (cobalamin) transporter, exporting vitamin B12 from the lysosomes into the cytoplasm [147]. However, it was later shown that LMBD1 actually interacts with ABCD4 and assists in its lysosomal trafficking [148], and that ABCD4 transports vitamin B12 even in the absence of LMBD1 [149]. Therefore, it is likely that LMBD1 itself is not a vitamin B12 transporter. LMBD1 was originally coined as having “significant homology” to lipocalin membrane receptors [147], and indeed the LIMR (lipocalin-1-interacting membrane receptor) protein, encoded by the LMBR1L gene, is responsible for binding lipocalin 1 (LCN-1) with high affinity [150,151]. LMBRD2 was proposed to be a regulator of β2-adrenoceptor signaling [152], while the first protein identified in the family, LMBR1, was associated with polydactyly and limb malformations [153,154]. However, its physiological role is still elusive. The proteins seem to contain 9 TMHs in a 5+4 arrangement according to UniProt predictions, but the tertiary structure of the proteins is still unknown, and no homologs with a known structure were found in our search.

Two MagT1-like proteins (**MAGT1** and **TUSC3**), as well as **OSTC** and **OSTCL** (oligosaccharyltransferase complex subunit) turned up in our search, showing similarity to TCDB family #1.A.76 members. MAGT1 and TUSC3 also have high-scoring hits for the Pfam domain “OST3_OST6”, which is characteristic of members of the oligosaccharyltransferase (OST) complex. TUSC3 (also called N33) was first identified as a tumor suppressor gene [155], and its presence, together with that of MAGT1 in the OST complex has been attested later on [156–160]. Therefore, it was suggested that these proteins act as oxidoreductases [159]. Meanwhile, MAGT1 and TUSC3 were also proposed to act as Mg^2+^ transporters [161,162]. On the other hand, recent structural findings of human MAGT1 [160] indicated that this protein may not function as a transporter or channel due to the lack of substrate-binding site or pore. OSTC (also called DC2) has similarly been shown to be part of the OST complex and to have a structure similar to MAGT1 [160]. Whether MagT1-like proteins still have a transport function remains to be clarified.

The **TMEM14A**, **TMEM14B** and **TMEM14C** proteins are the only ones in human containing the “Tmemb_14” Pfam domain. While Pfam lists this domain as functionally uncharacterized, a plant protein (FAX1) containing this domain was suggested to be involved in fatty acid export from chloroplasts [163]. However, the physiological roles of TMEM14A and TMEM14B in human remain elusive. The third member of the family, TMEM14C, was identified as a putative mitochondrial protein whose transcript is consistently coexpressed with proteins from the core machinery of heme biosynthesis [164]. It was later shown that TMEM14C mediates the import of protoporphyrinogen IX (PPgenIX) into the mitochondrial matrix [165,166]. While the structure of TMEM14C was solved using nuclear magnetic resonance (NMR) [167], showing a bundle of three TM helices and an amphipathic helix, the transport mechanism remains elusive. Interestingly, despite their proposed function in the mitochondria, TargetP-2.0 [168] did not predict a mitochondrial targeting sequence in the amino acid sequence of any of the human TMEM14 proteins in our hands.

**TMEM205** is a 4-TMH protein according to UniProt annotations, which was linked to cisplatin resistance [169]. The protein is expressed mostly in liver, pancreas and adrenal glands, and is present on the plasma membrane [169]. TMEM205-mediated resistance was shown to be selective towards platinum-based drugs, such as cisplatin and oxaliplatin, but not carboplatin [170]. While structural information about the protein is not available, mutagenesis studies of TMEM205 showed that mutating sulfur-containing residues, especially in TMH2 and TMH4, diminishes the effect of cisplatin resistance [170]. Nevertheless, neither the biological function nor the physiological substrates of TMEM205 are known.

#### Transporters with hydrophobic substrates

In addition to proteins that transport solutes or are similar to transporters that typically translocate water-soluble small molecule compounds, our search has uncovered numerous proteins that have been reported to take part in modulating the intracellular distribution of hydrophobic or amphipathic compounds, such as cholesterol, fat-soluble vitamins, lipids and fatty acids (Table 3). Although these proteins do not, strictly speaking, transport so-called “solutes”, they translocate small hydrophobic molecules that have fundamental biological functions. Thus one could argue that they belong to the SLC superfamily as well. Accordingly, they have been integrated in our search and some of them have already been included in the SLC nomenclature (SLC59, SLC63, SLC65). In view of the biological and pharmaceutical importance of the transport mechanisms of hydrophobic substrates, our results on this topic will be discussed in a separate paper.

### Conclusions

Our study represents the first systematic correlation of the SLC and TCDB nomenclature schemes. Many of the transport proteins discovered in our search are underexplored and there is limited information about them, although they often have important physiological roles and/or potentially represent new therapeutic targets. Even with proteins that have been studied for their physiological involvement, it was often not taken into account that they could have a transport function. Numerous proteins uncovered in our search have similarity to proteins with transport function in other organisms, but their physiological substrates remain unknown. These proteins will be interesting targets for deorphanization studies in order to reveal their natural substrates. In our work, we also highlight proteins for which transport activity has been controversial, and more specific analyses are required to clarify their biological function. In addition, our search reveals new SLC-like proteins that have no structural information. This hinders a deeper understanding of their transport mechanism. Future structure determinations would be of crucial importance to accelerate validation of the identified proteins. The combination of all these efforts would greatly facilitate the completion of the SLC-ome in human cells. Thus, our study points out important directions in which future studies could help resolve the lack of information about SLC transporters, which will help unlock their therapeutic potential.

## Acknowledgements

We acknowledge support by the Swiss National Science Foundation for the grants # CRSII5_180326 (“The role of mitochondrial carriers in metabolic tuning and reprogramming by calcium flow across membrane contact sites”) and # 310030_182272 (“Intestinal absorption of transition metals in human health and disease”) as well as by the Swiss National Research Programme NRP 78, # 4078P0_198281 (“New insights into the COVID-19 pandemic”). Calculations were performed on UBELIX (http://www.id.unibe.ch/hpc), the HPC cluster at the University of Bern.

## Methods

### HMM building for TCDB families and subfamilies

Sequences of selected TCDB families and subfamilies were aligned using PSI-Coffee 11.00 [171] and NCBI BLAST+ 2.6.0 [172] using the “nr” BLAST database of 2018-02-12. PSI-Coffee was run with BLAST mode “LOCAL” and otherwise with default options. Altogether, 4221 different sequences from the TCDB were present in the subfamily and family alignments, and the profiles used for alignments contained 1-1070 sequences. In total, 130 sequences from the TCDB returned zero hits from the “nr” database in the iterative BLAST searches, meaning their profiles just contained the query sequence itself. PSI-Coffee uses the profiles to guide the generation of multiple alignments of the query sequences within each subfamily and family. The alignments were subsequently turned into HMMs using the “hmmbuild” command of HMMER 3.1b2 [30] using default settings.

### Sequence similarity search

All protein belonging to each of the 7 organisms studied were downloaded from UniProt on 2019-07-31 into a FASTA file. Sequence similarity search was performed using the HMMs downloaded for selected Pfam families on 2019-02-14 and those generated for selected TCDB families and subfamilies using “hmmsearch” from HMMER 3.1b2. Hits with bit scores larger than 50 were used for further analysis. These hits presented a maximal hit e-value of 7.9e-12, while hits with bit score larger than 25, used for HMM fingerprint-based clustering (see below), had hit e-values below 6e-4.

### Sequence clustering for fragment elimination

Since the downloaded protein set from UniProt contained fragments as well as predicted open reading frames and sequences from genomic screening methods, we strived to retain one sequence per gene for further analysis. In order to achieve this, hits yielded by the sequence similarity search were clustered using the following method. First, all-against-all similarity searches were performed using NCBI BLAST+ 2.4.0 [172]. Sequences were assigned the same cluster if they either share common gene annotations according to their UniProt records, or a high-scoring segment pair (HSP) with more than 95% sequence identity based on the all-against-all BLAST search. For gene annotation, the fields Gene Symbol (GN), HGNC symbol, GeneID, UniGene, FlyBase, KEGG identifiers were used from UniProt records. Ambiguous or conflicting annotations, as well as annotations conflicting with BLAST-reported sequence similarity were detected and resolved manually. Afterwards, clusters were reduced to representative sequences. For clusters containing a single Swiss-Prot sequence, that sequence was taken as representative. For cluster with no Swiss-Prot sequence, the longest sequence of the cluster was taken as representative. Clusters with more than one Swiss-Prot sequence were manually analyzed and split if necessary.

### HMM fingerprint-based sequence clustering

We introduce the concept of a “HMM fingerprint”, which is a mathematical vector of numbers assigned to a protein sequence, corresponding to the bit scores of similarity to each of the set of HMMs used in our analysis, consisting of the HMMs of TCDB families and subfamilies, as well as Pfam HMMs. We have restricted the number of HMMs to those that gave a hit with bit score > 25, in total 513 HMMs. Two proteins sequences that are related are expected to show similarity to a similar subset of TCDB families, subfamilies, or Pfam families, and therefore a similar pattern in their HMM fingerprints. In turn, if two protein sequences show similarity to the same subset of TCDB families, subfamilies or Pfam families, as indicated by a similar HMM fingerprint, then they can be expected to be related. Once the HMM fingerprint has been assigned to each protein sequence found in our search, the unweighted pair group method with arithmetic mean (UPGMA) method [173] was used using the cosine metric to arrive at a hierarchical clustering of the sequences. The tree representing the clustering was cut at 0.7 cosine distance to arrive at branches that formed the basis of protein families. Join and split constraints have been introduced to keep certain proteins artificially grouped or separated. If a join constraint was acting between a pair of proteins and they were not grouped into the same cluster by the default tree cut threshold, then the clustering tree was cut just above the branch that leads to the smallest cluster containing both proteins. Similarly, split constraints between two proteins caused the clustering tree to be cut just below the branch containing both proteins, unless the two proteins were already in different clusters. Join constraints were introduced between the following protein pairs: SLC5A1-SLC5A7, SLC9A1-SLC9B1, SLC10A1-SLC10A7, SLC25A1-SLC25A46, SLC30A1-SLC30A9, SLC35C1-SLC35G1, SLC39A1-SLC39A9, and SLC66A1-SLC66A5. Split constraints were introduced between the following protein pairs: SLC2A1-SLC22A1, SLC17A1-SLC18A1, SLC17A1-SLC37A1, SLC32A1-SLC36A1, and SLC32A1-SLC38A1. The clustering tree with the resulting families was shown in Fig 1 in a polar coordinate system, with the radial component (*d* ∈ [0; 1]) transformed according to 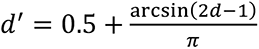 to magnify tree details around *d* = 0 and *d* = 1 for better visual representation.

### Structural homolog search

Sequences of SLC-like proteins were turned into HMMs using “hhblits” from the HH-suite3 package [174], using the UniRef30 database of 2020-06 [35,175], 3 iterations, an E-value threshold of 1e-3 for inclusion, and a probability threshold of 0.35 for MAC re-alignment. The HMMs were searched against the “pdb70” database of 2021-08-04 [35] with no MAC realignment, “predicted vs predicted” secondary structure scoring, and amino acid score of 1. The resulting hits were checked against PDB annotations of transmembrane helices, and were accepted if at least 3 TM segments were contained within the aligned region and the E-value of the hit was less than 1e-4.

### Phylogenetic trees

Selected groups of SLC-like proteins were aligned using Clustal Omega 1.2.1 [176,177] with 5 iterations and default settings. Smart Model Selection 1.8.4 [178] and PhyML 3.3.20190909 [179] were used to generate the phylogenetic trees, with 10 random starting trees and using the approximate likelihood ratio test aLRT method [180]. Trees in main figures were visualized using TreeViewer 1.2.2 (https://treeviewer.org/). Tree rooting, rearrangement (with threshold 0.9) and reconciliation with the species tree was done using NOTUNG 2.9.1.5 [181]. Reconciled phylogenetic trees were visualized using custom Python scripts in the style used by NOTUNG. Orthologs were identified using the reconciled phylogenetic trees and custom Python scripts.

### Data Availability

Scripts and data generated during the preparation of the manuscript are available at https://github.com/bioparadigms/SLCAtlas. In addition, the datasets resulting from this work will also be made available in a browsable format at https://www.bioparadigms.org/.

## Supplementary information

**S1 Table. “SLC-like” Pfam models and their clan memberships.** The dataset is a result of our manual curation based on our “SLC-like” criteria (see text) and Pfam families present in protein sequences listed in the TCDB. Family names and data are from the Pfam database.

**S2 Table. False positive hits or incorrectly annotated human sequences found in our search.** Gene symbol and protein name shown based on UniProt data. Most similar (highest scoring) TCDB families/subfamilies and Pfam families are shown, and reason for being false positive are indicated.

**S3 Table. Table of existing (classic) SLC transporters, indicating the correlation with TCDB families/subfamilies, Pfam families, PDB structures and structural folds.** Gene symbol and protein name based on UniProt information are shown. Most similar (highest-scoring) TCDB families and subfamilies, as well as Pfam families are shown for each protein. The PDB structure from the pdb70 dataset that is most similar (highest-scoring) to each protein is shown, along with its fold family assignment by us, partly based on TCDB family names. AbgT: p-Aminobenzoyl-glutamate Transporter family; Amt: Ammonium Transporter family; APC: Amino acid-Polyamine-Cation family; CDF: Cation Diffusion Facilitator family; CNT: Concentrative Nucleoside Transporter family; Ctr: Copper Transporter family; DAACS: Dicarboxylate/Amino Acid:Cation (Na^+^ or H^+^) Symporter family; MATE: Multidrug And Toxic compound Extrusion family; MCF: Mitochondrial Carrier Family; MFS: Major Facilitator Superfamily; MgtE: Mg^2+^ transporter-E family; NAT: Nucleobase/Ascorbate Transporter or Nucleobase:Cation Symporter-2 (NCS2) family; NCX: Sodium/Calcium exchanger family; NhaA: Sodium/proton antiporter family; NST: Nucleoside-Sugar Transporter family; PiT: Type III Sodium/phosphate cotransporter family; SWEET: Sugar Will Eventually be Transported family.; ZIP: Zrt/Irt-like Transporter family.

**S4 Table. Non-classic “SLC-like” proteins of Table 3 found in our search, extended with additional information.** Protein sequence identifiers, most similar (highest-scoring) PDB structure from the pdb70 dataset, chemical identifiers, PubMed links to references, and comments have been included. The table also contains information about whether individual SLC-likeness criteria (numbered 1-8) have been satisfied or violated.

**S1 File. Phylogenetic trees of SLC-like protein families with more than two members.** Trees were generated using multiple alignment by ClustalO, maximum likelihood tree generation by PhyML, followed by tree reconciliation with the species tree using NOTUNG (see Methods). The species tree with internal names of putative ancestor taxa is shown on each page on the upper-right hand corner. The trees are shown as dendrograms and branch lengths are not indicative of evolutionary distance. Each tree leaf corresponds to an SLC-like protein sequence denoting a gene, labels show the gene symbol, UniProt accession and taxon name. Leaves with labels ending with “*LOST” denote putative genes lost in the indicated ancestral species. Red “D” denote gene duplication nodes, normal nodes correspond to speciation nodes. Light green numbers denote branch support values as calculated by NOTUNG.

