## Supplementary material for "Systematic *in silico* discovery of novel solute carrier-like proteins from proteomes": S1 File

### SLC1 family

Species tree:

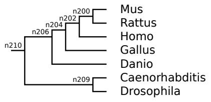


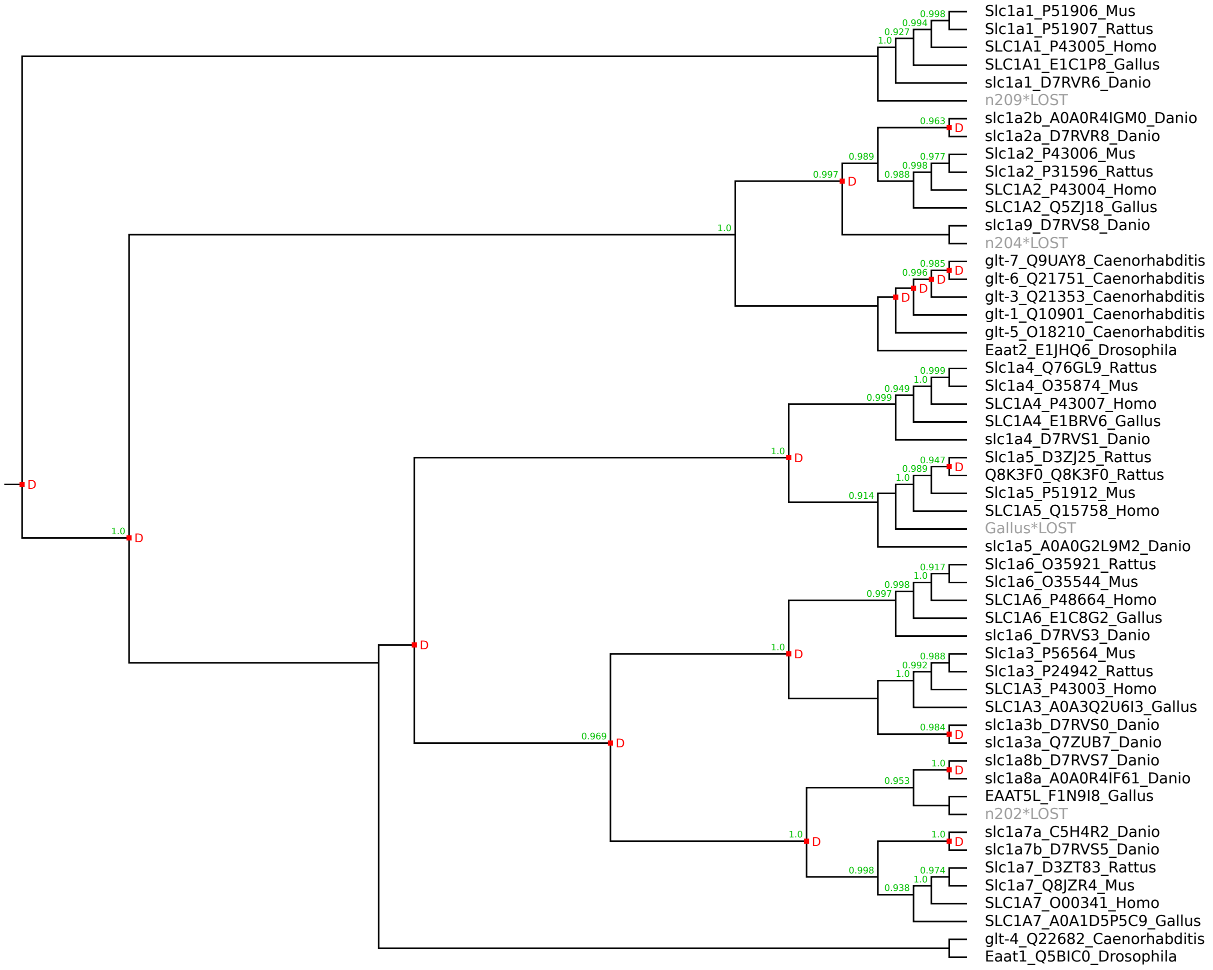

### SLC2 family

Species tree:

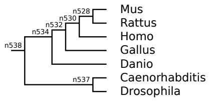


Warning: this is a large figure that had to be reduced to fit on the page.
Please use the zoom function of your PDF viewer to see the details.

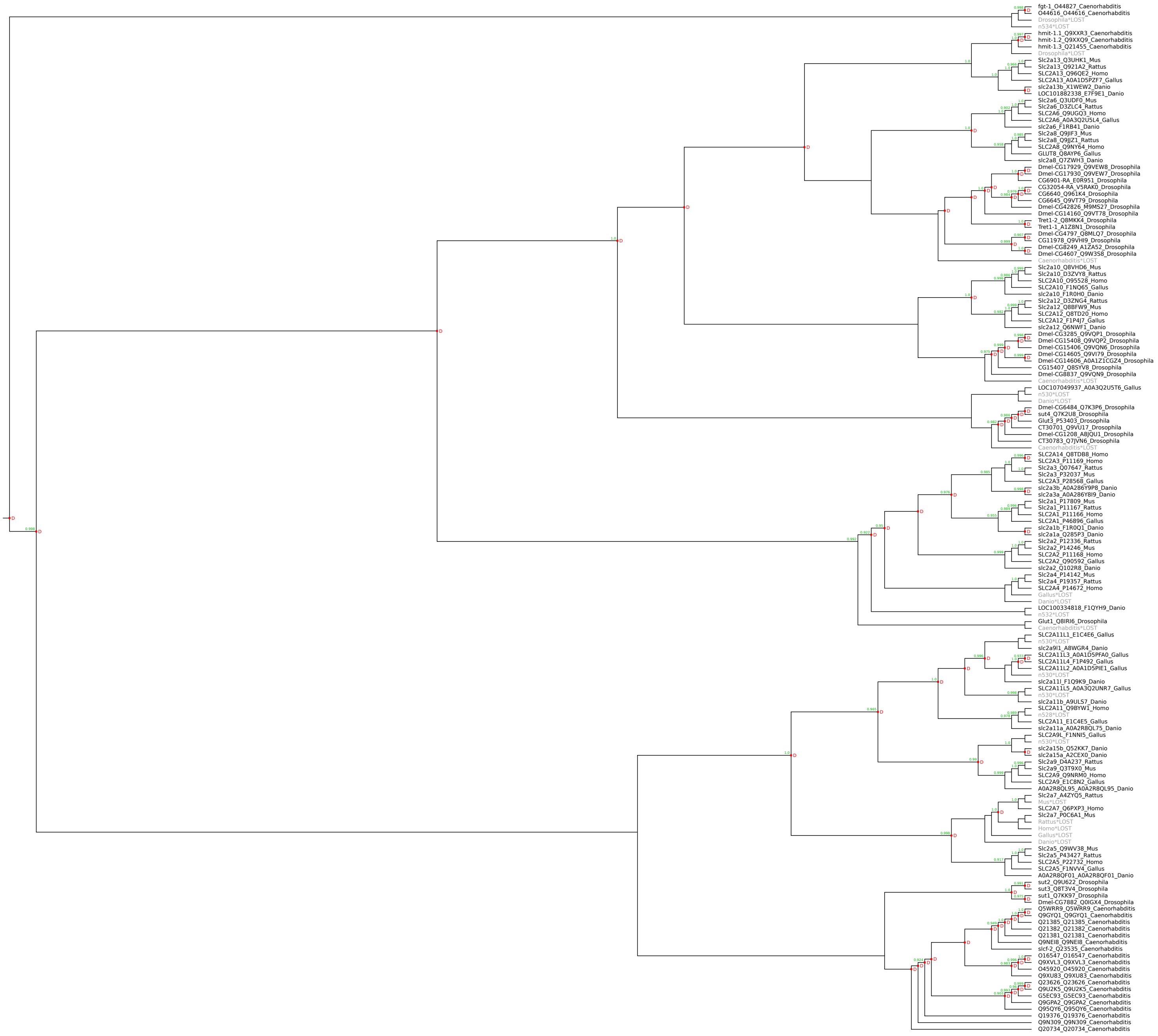

### SLC4 family

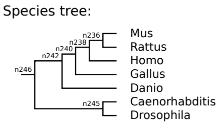

Species tree:
n236
n238
n240
n242
n246
n245
Mus
Rattus
Homo
Gallus
Danio
Caenorhabditis
Drosophila


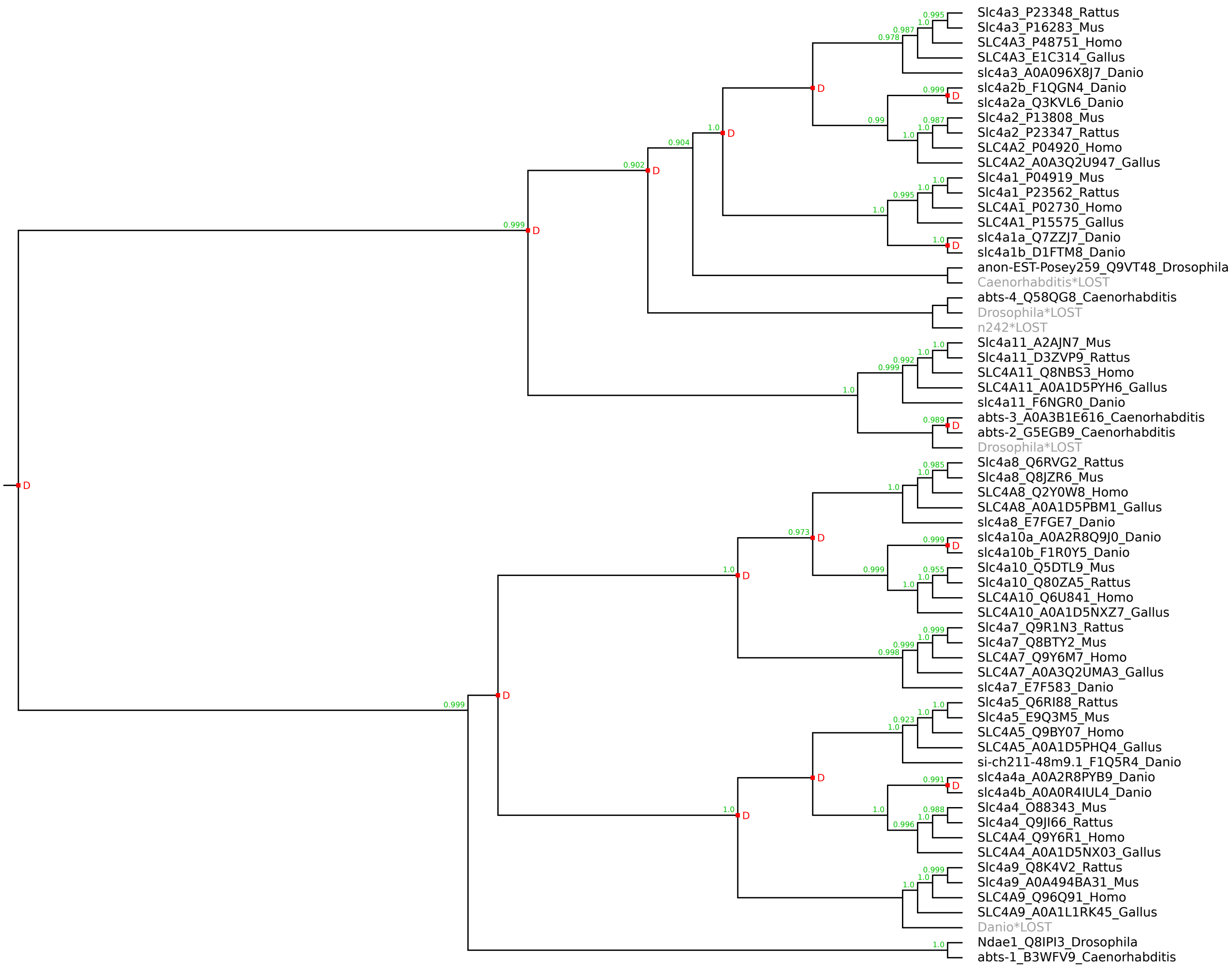

Slc4a3_P23348_Rattus
Slc4a3_P16283_Mus
SLC4A3_P48751_Homo
SLC4A3_E1C314_Gallus
slc4a3_A0A096X8J7_Danio
slc4a2b_F1QGN4_Danio
slc4a2a_Q3KVL6_Danio
Slc4a2_P13808_Mus
Slc4a2_P23347_Rattus
SLC4A2_P04920_Homo
SLC4A2_A0A3Q2U947_Gallus
Slc4a1_P04919_Mus
Slc4a1_P23562_Rattus
SLC4A1_P02730_Homo
SLC4A1_P15575_Gallus
slc4a1a_Q7ZZJ7_Danio
slc4a1b_D1FTM8_Danio
anon-EST-Posey259_Q9VT48_Drosophila
Caenorhabditis*LOST
abts-4_Q58QG8_Caenorhabditis
Drosophila*LOST
n242*LOST
Slc4a11_A2AJN7_Mus
Slc4a11_D3ZVP9_Rattus
SLC4A11_Q8NBS3_Homo
SLC4A11_A0A1D5PYH6_Gallus
slc4a11_F6NGR0_Danio
abts-3_A0A3B1E616_Caenorhabditis
abts-2_G5EGB9_Caenorhabditis
Drosophila*LOST
Slc4a8_Q6RVG2_Rattus
Slc4a8_Q8JZR6_Mus
SLC4A8_Q2Y0W8_Homo
SLC4A8_A0A1D5PBM1_Gallus
slc4a8_E7FGE7_Danio
slc4a10a_A0A2R8Q9J0_Danio
slc4a10b_F1R0Y5_Danio
Slc4a10_Q5DTL9_Mus
Slc4a10_Q80ZA5_Rattus
SLC4A10_Q6U841_Homo
SLC4A10_A0A1D5NXZ7_Gallus
Slc4a7_Q9R1N3_Rattus
Slc4a7_Q8BTY2_Mus
SLC4A7_Q9Y6M7_Homo
SLC4A7_A0A3Q2UMA3_Gallus
slc4a7_E7F583_Danio
Slc4a5_Q6RI88_Rattus
Slc4a5_E9Q3M5_Mus
SLC4A5_Q9BY07_Homo
SLC4A5_A0A1D5PHQ4_Gallus
si-ch211-48m9.1_F1Q5R4_Danio
slc4a4a_A0A2R8PYB9_Danio
slc4a4b_A0A0R4IUL4_Danio
Slc4a4_O88343_Mus
Slc4a4_Q9JI66_Rattus
SLC4A4_Q9Y6R1_Homo
SLC4A4_A0A1D5NX03_Gallus
Slc4a9_Q8K4V2_Rattus
Slc4a9_A0A494BA31_Mus
SLC4A9_Q96Q91_Homo
SLC4A9_A0A1L1RK45_Gallus
Danio*LOST
Ndae1_Q8IPI3_Drosophila
abts-1_B3WFV9_Caenorhabditis

Species tree:

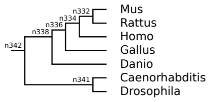


### SLC5 family

Warning: this is a large figure that had to be reduced to fit on the page.
Please use the zoom function of your PDF viewer to see the details.

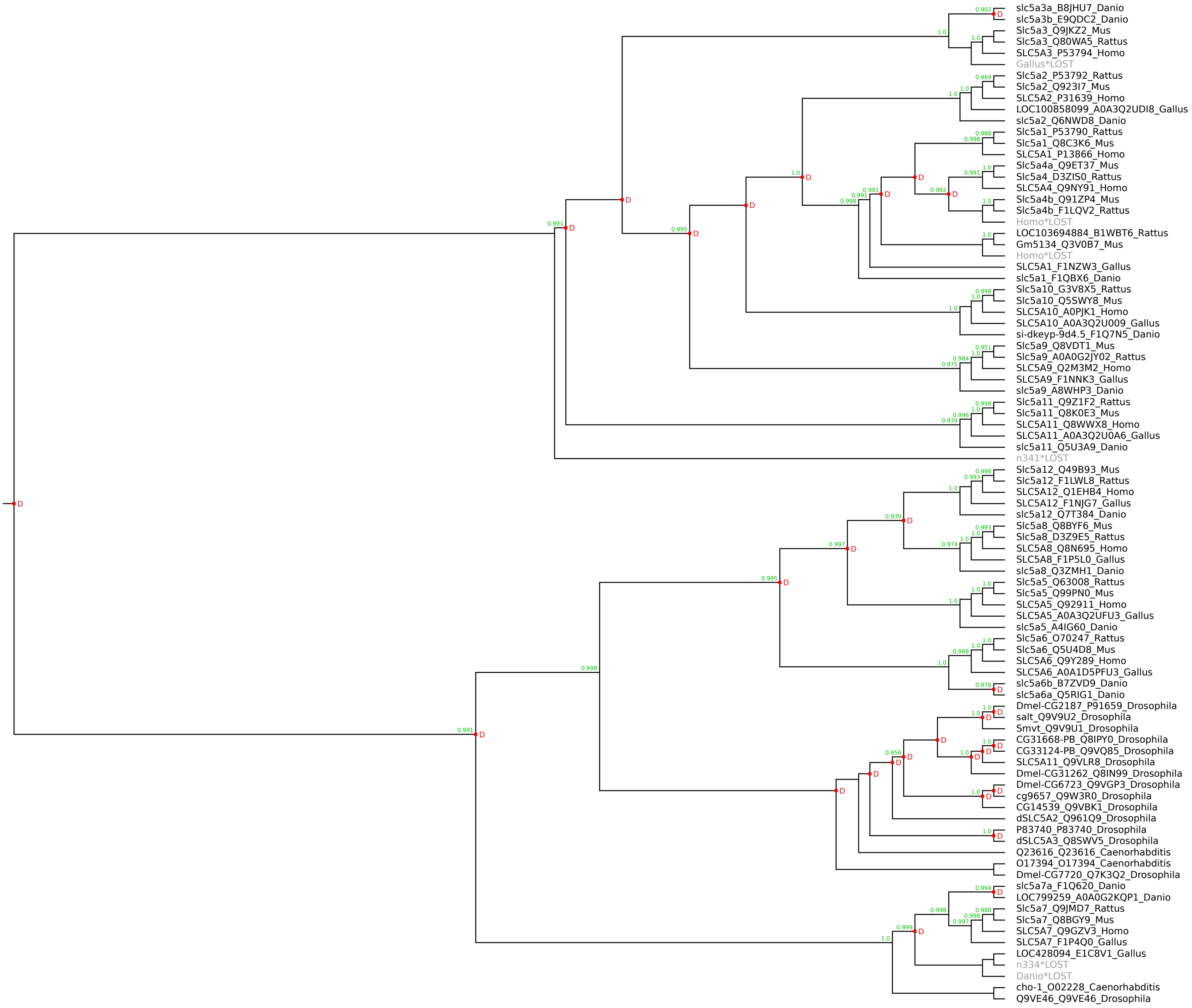

Species tree:

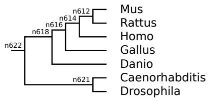


### SLC6 family

Warning: this is a large figure that had to be reduced to fit on the page.
Please use the zoom function of your PDF viewer to see the details.

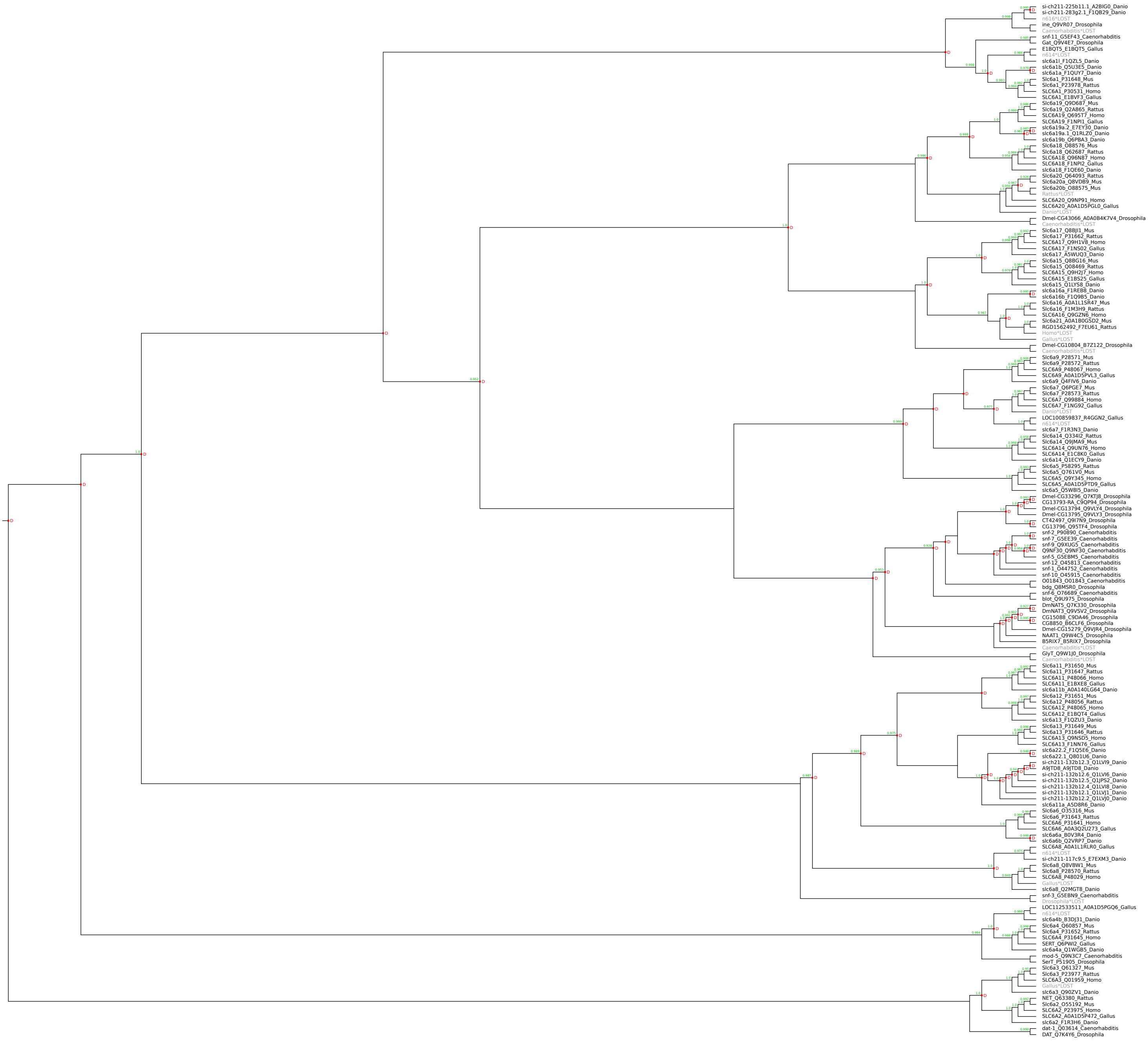

### SLC7 family

Species tree:

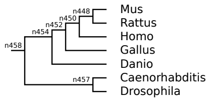


Warning: this is a large figure that had to be reduced to fit on the page.
Please use the zoom function of your PDF viewer to see the details.

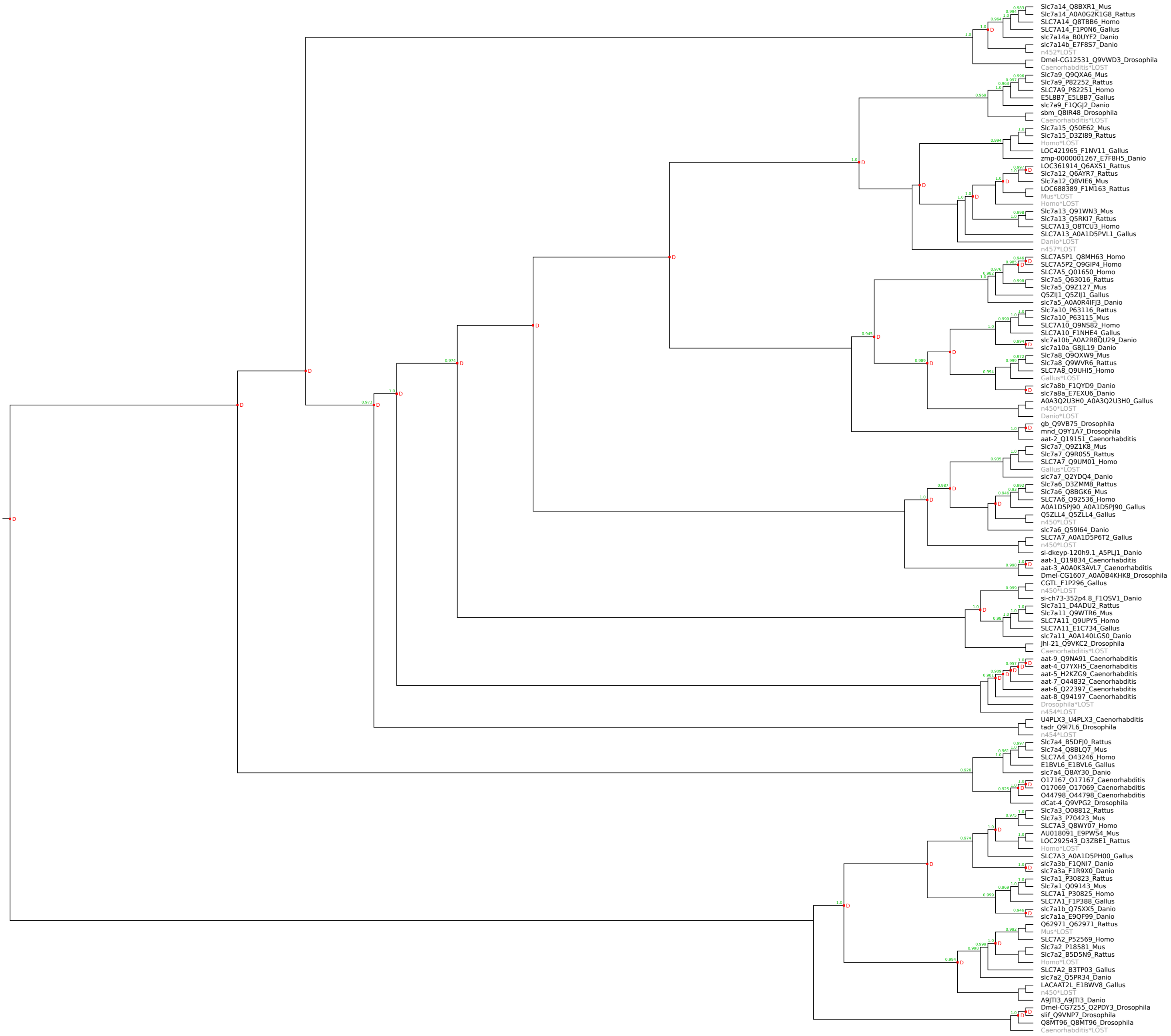

### SLC8 family

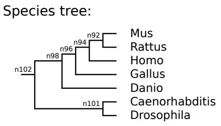

Species tree:
n92
n94
n96
n98
n102
n101
Mus
Rattus
Homo
Gallus
Danio
Caenorhabditis
Drosophila


---

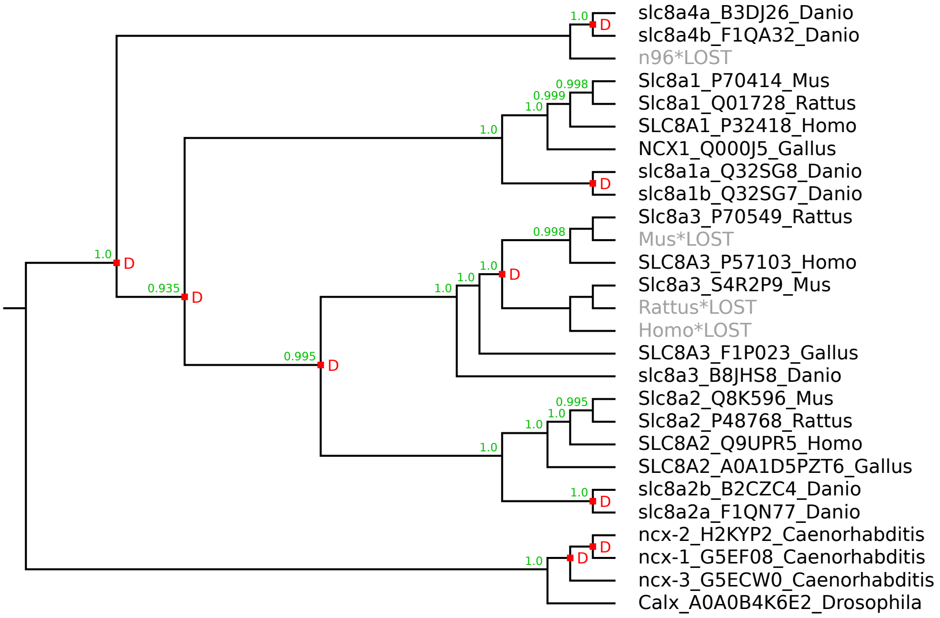

slc8a4a_B3DJ26_Danio
slc8a4b_F1QA32_Danio
n96*LOST
Slc8a1_P70414_Mus
Slc8a1_Q01728_Rattus
SLC8A1_P32418_Homo
NCX1_Q000J5_Gallus
slc8a1a_Q32SG8_Danio
slc8a1b_Q32SG7_Danio
Slc8a3_P70549_Rattus
Mus*LOST
SLC8A3_P57103_Homo
Slc8a3_S4R2P9_Mus
Rattus*LOST
Homo*LOST
SLC8A3_F1P023_Gallus
slc8a3_B8JHS8_Danio
Slc8a2_Q8K596_Mus
Slc8a2_P48768_Rattus
SLC8A2_Q9UPR5_Homo
SLC8A2_A0A1D5PZT6_Gallus
slc8a2b_B2CZC4_Danio
slc8a2a_F1QN77_Danio
ncx-2_H2KYP2_Caenorhabditis
ncx-1_G5EF08_Caenorhabditis
ncx-3_G5ECW0_Caenorhabditis
Calx_A0A0B4K6E2_Drosophila
1.0
0.998
0.999
0.935
0.995
D

### SLC9 family

Warning: this is a large figure that had to be reduced to fit on the page.
Please use the zoom function of your PDF viewer to see the details.

Species tree:

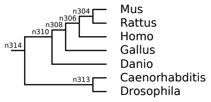


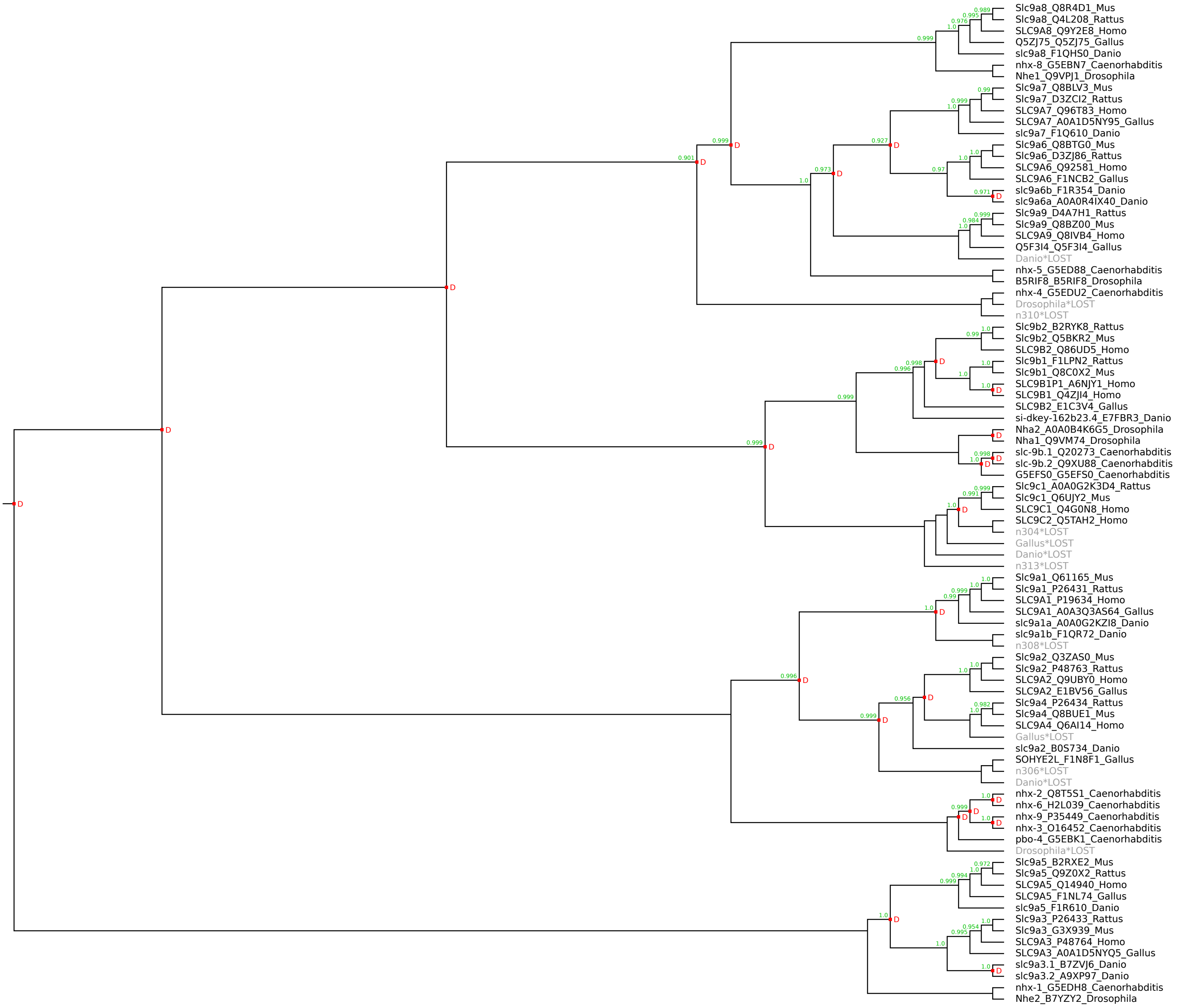

### SLC10 family

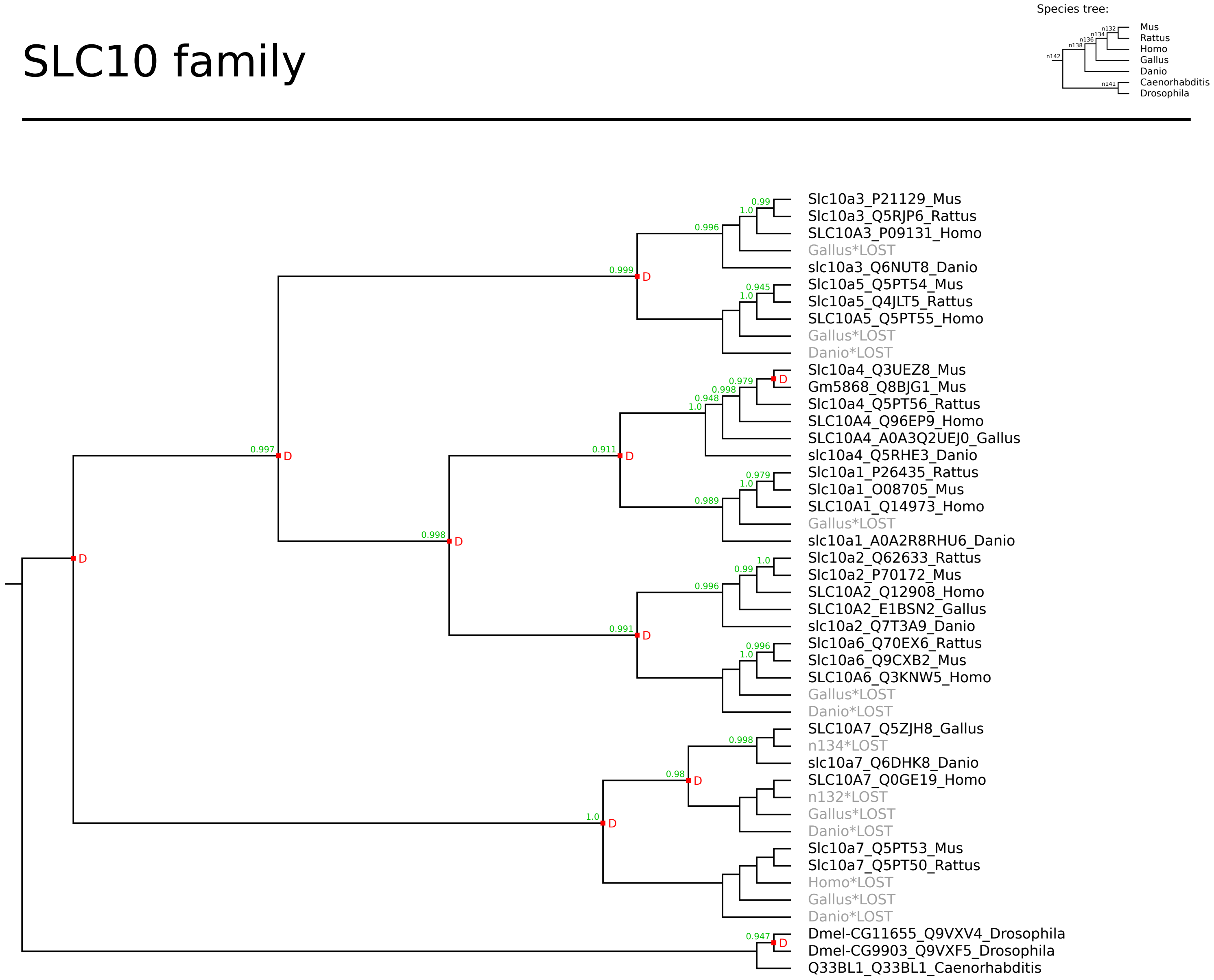

### SLC11 family

Species tree:

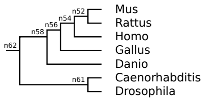


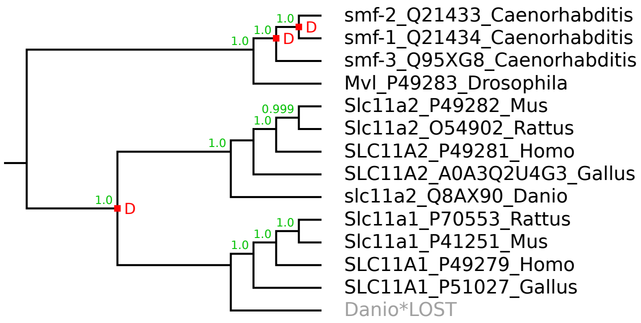

### SLC12 family

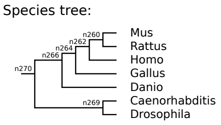


Warning: this is a large figure that had to be reduced to fit on the page.
Please use the zoom function of your PDF viewer to see the details.

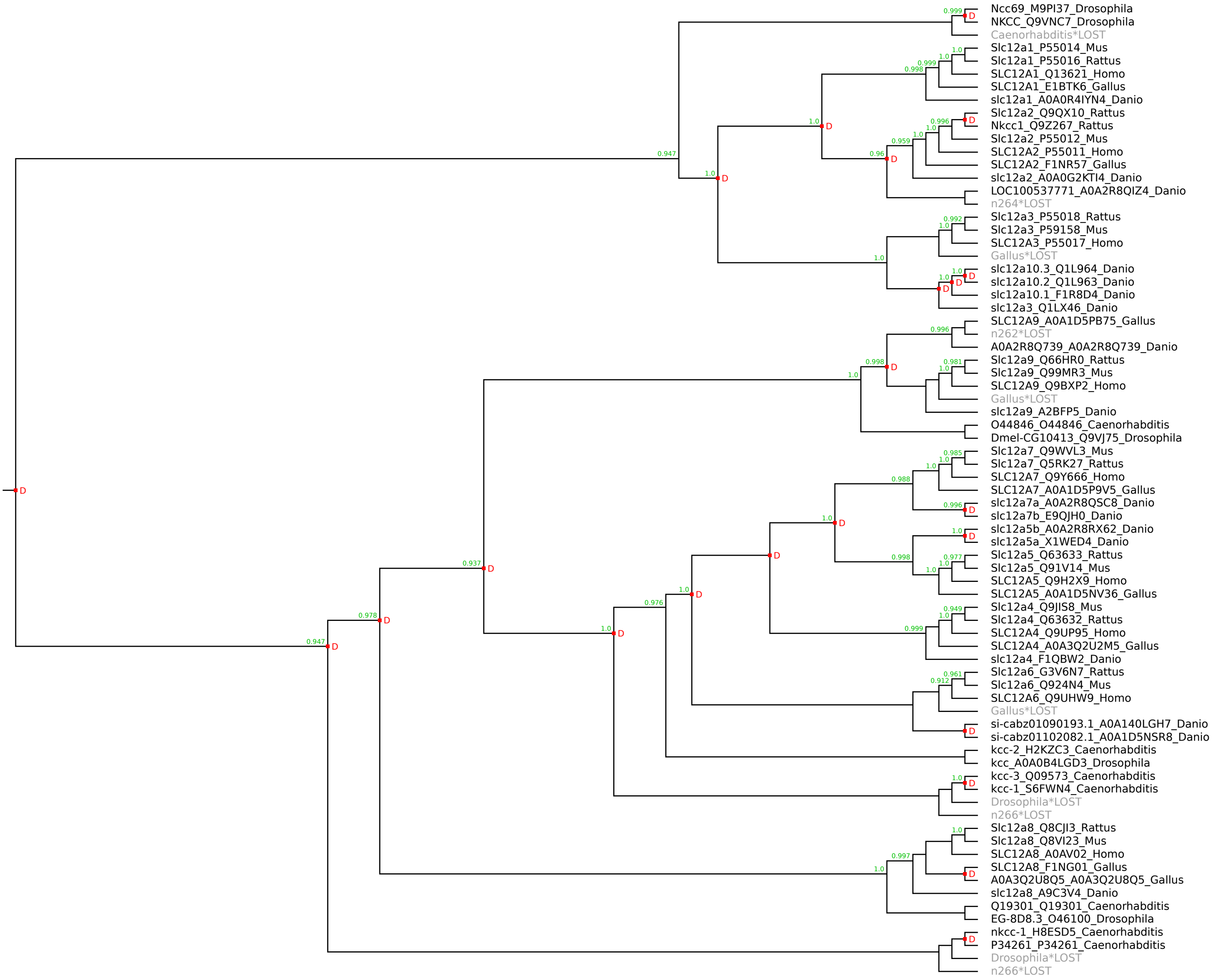

### SLC13 family

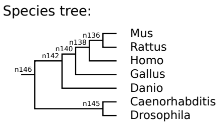


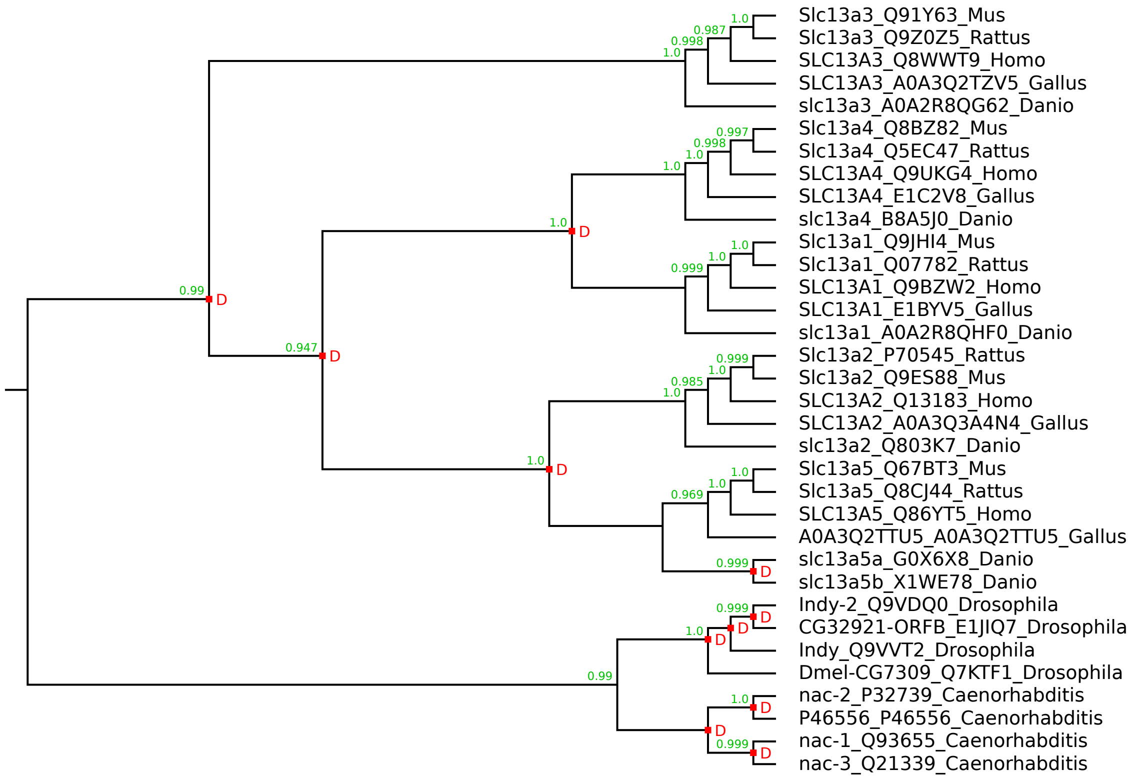

Species tree:

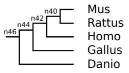


### SLC14 family

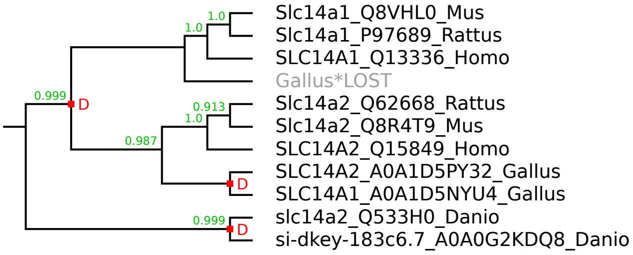

### SLC15 family

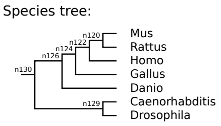


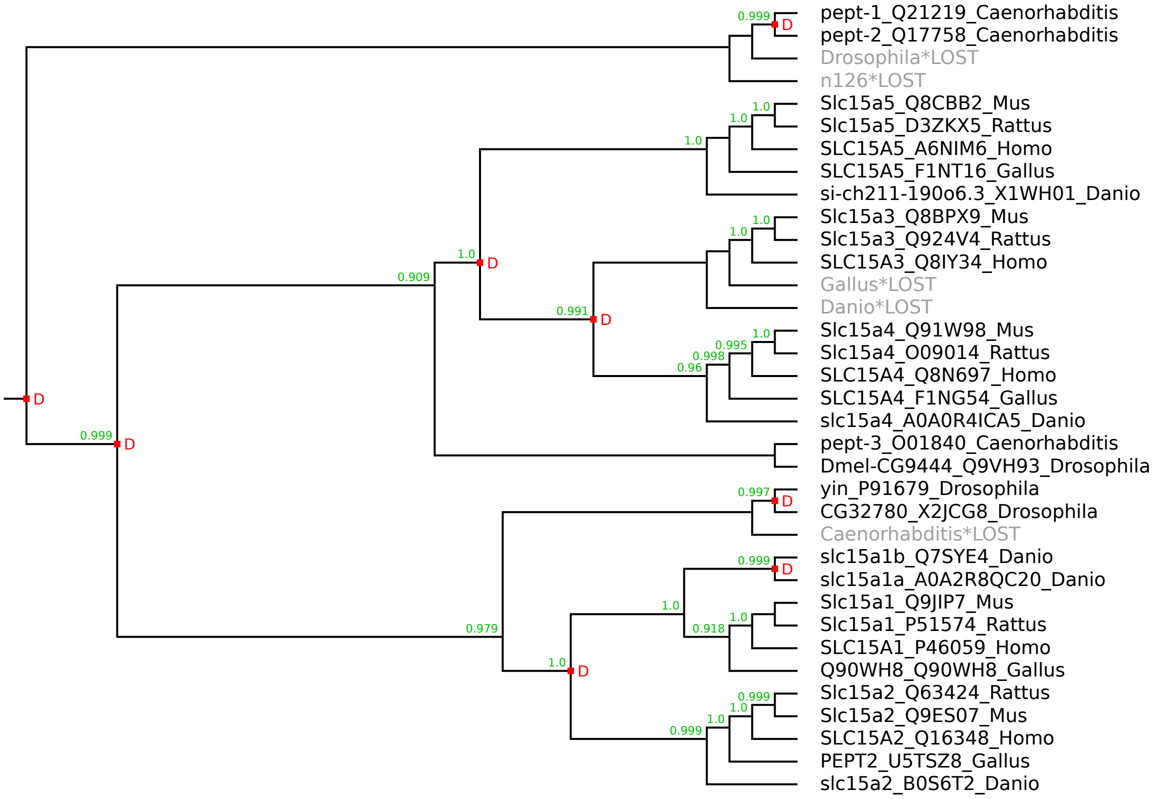

### SLC16 family

Species tree:

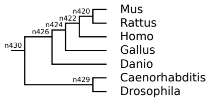


Warning: this is a large figure that had to be reduced to fit on the page.
Please use the zoom function of your PDF viewer to see the details.

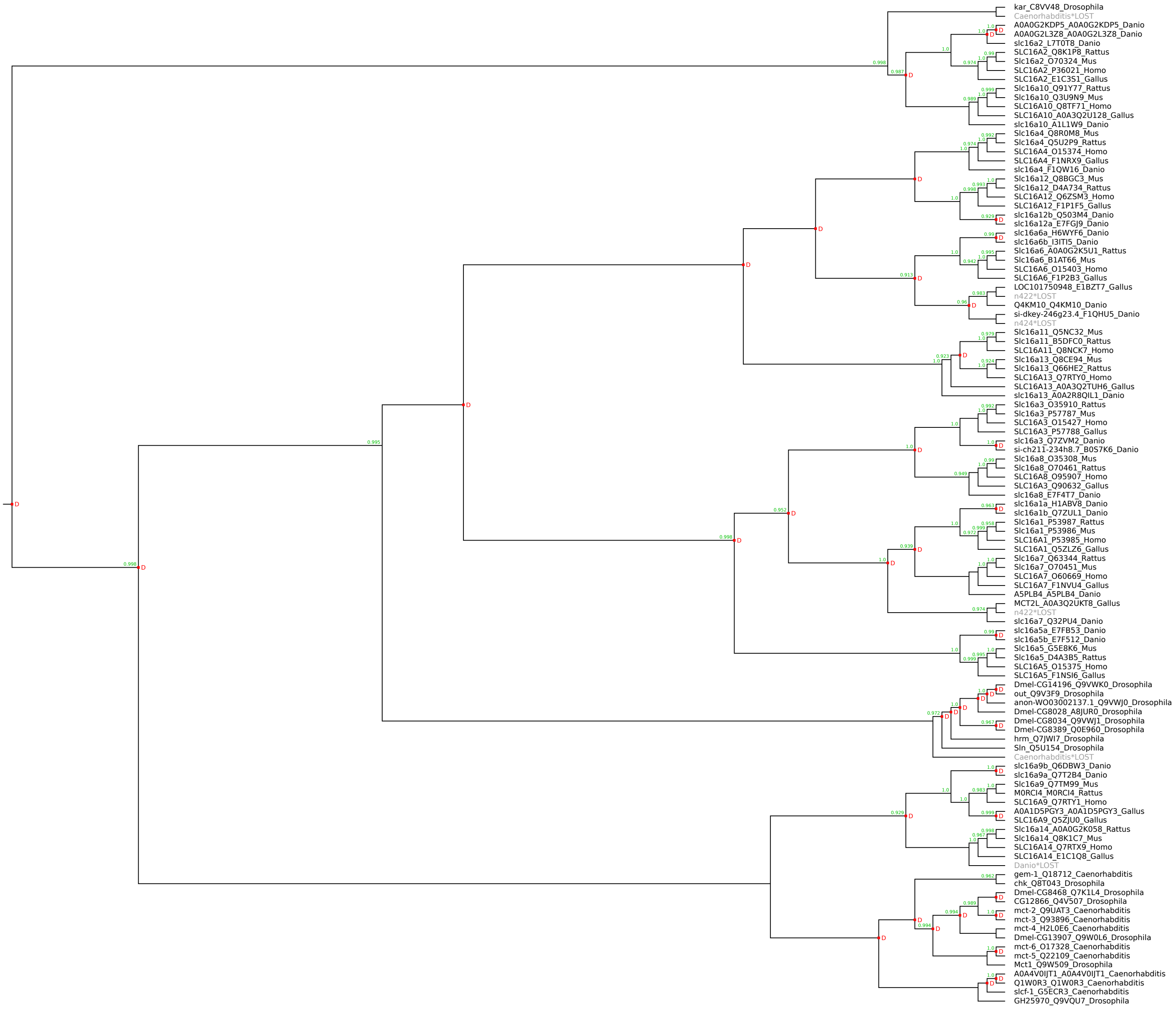

Species tree:

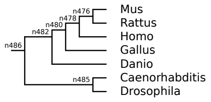


### SLC17 family

Warning: this is a large figure that had to be reduced to fit on the page.
Please use the zoom function of your PDF viewer to see the details.

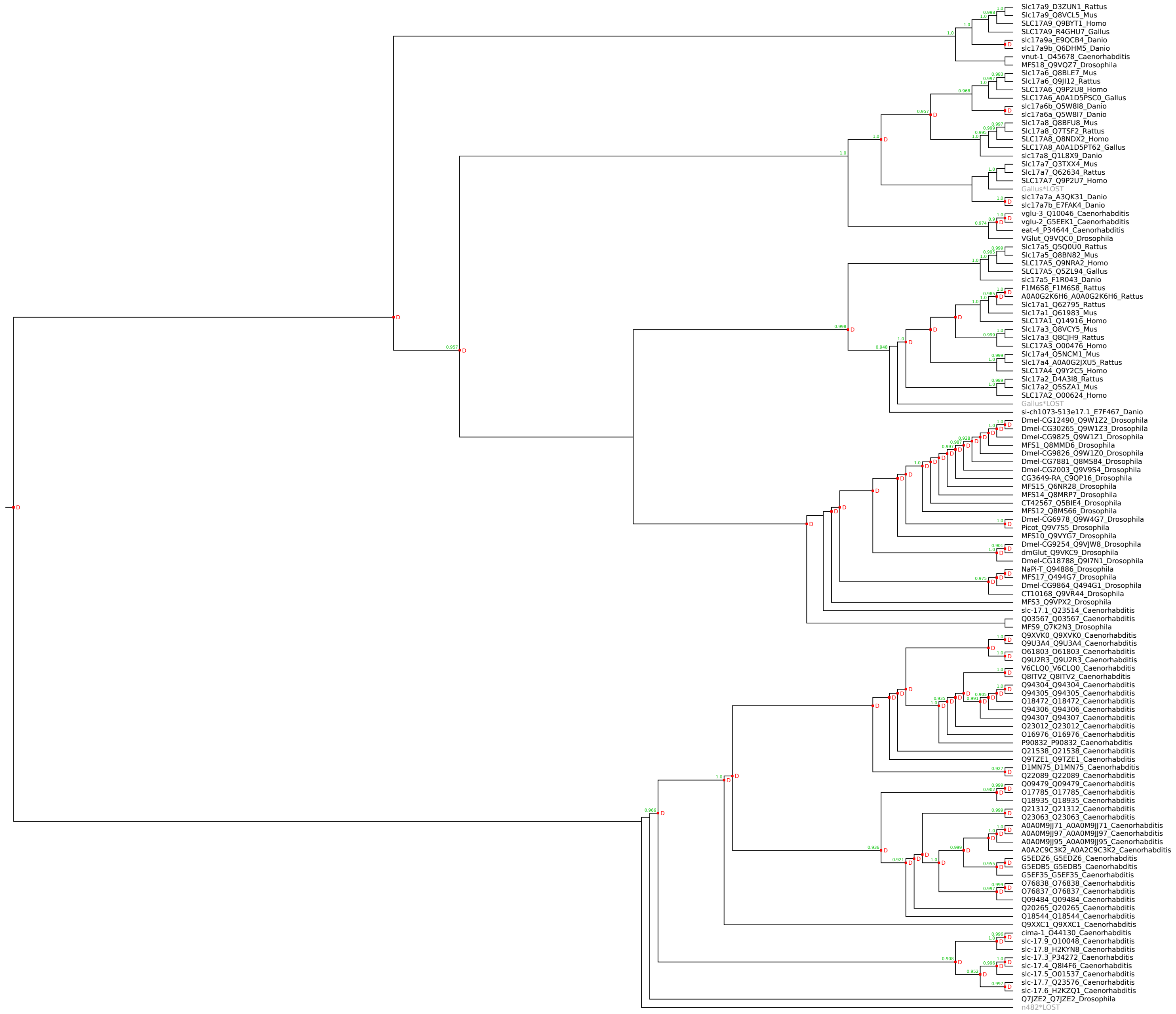

### SLC18 family

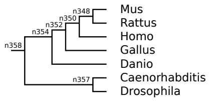


---

Warning: this is a large figure that had to be reduced to fit on the page.
Please use the zoom function of your PDF viewer to see the details.

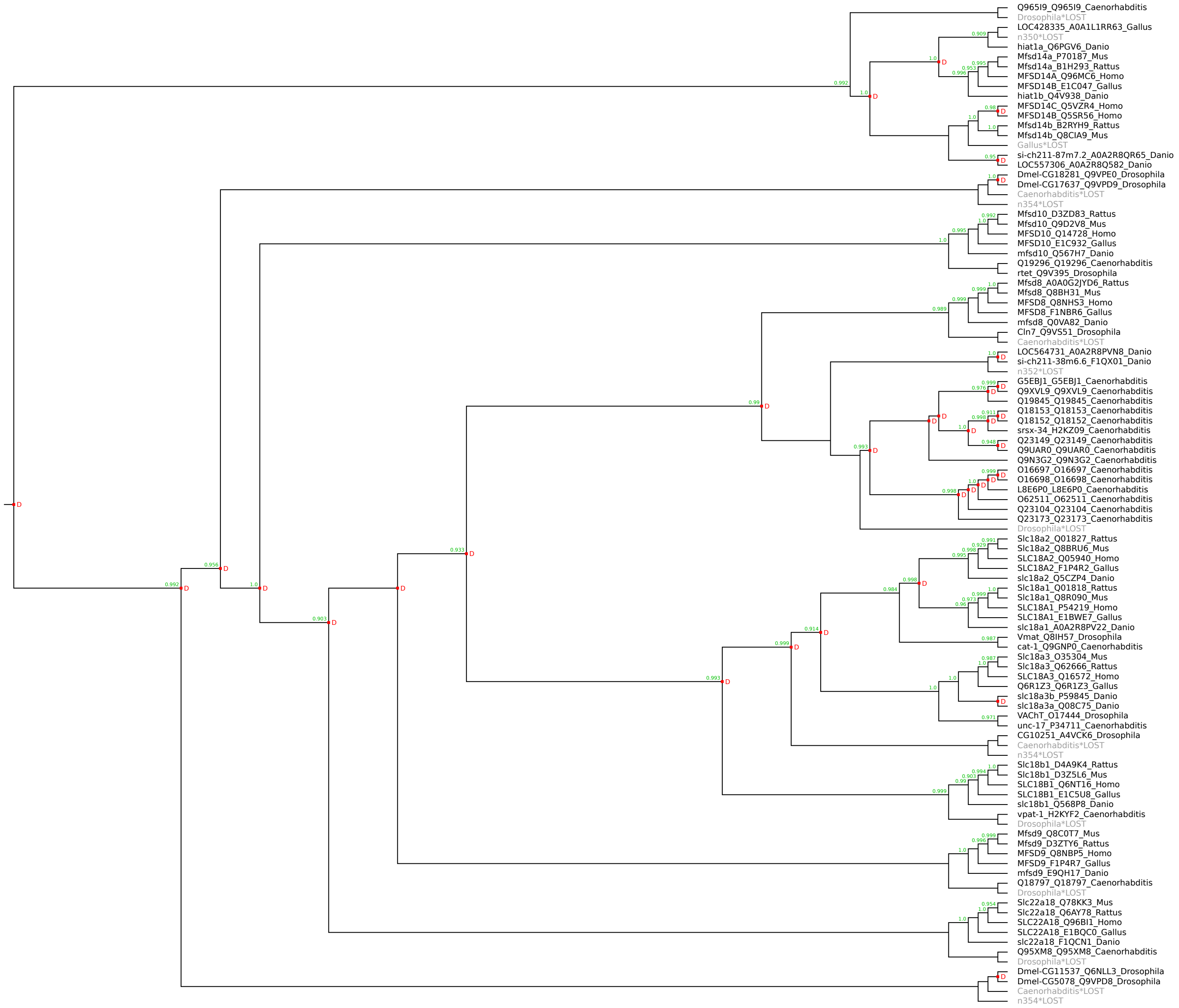

### SLC19 family

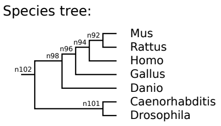


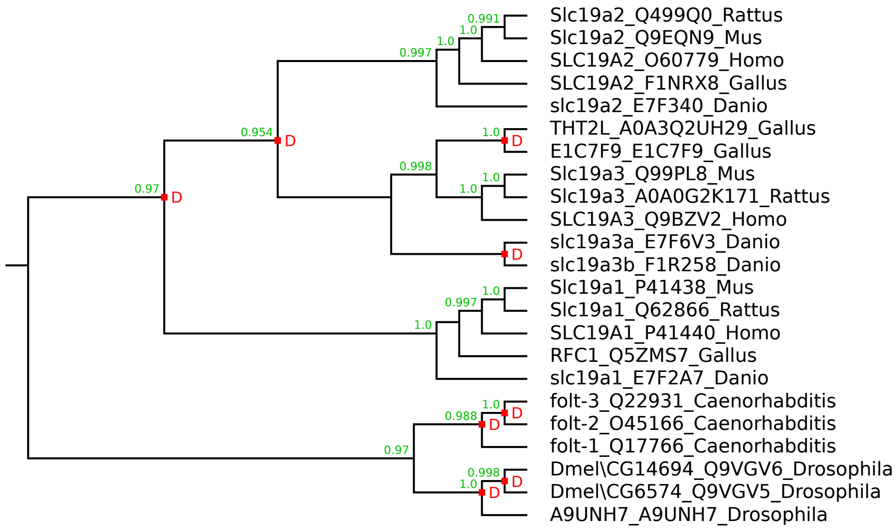

### SLC20 family

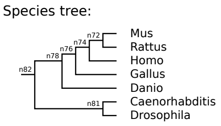


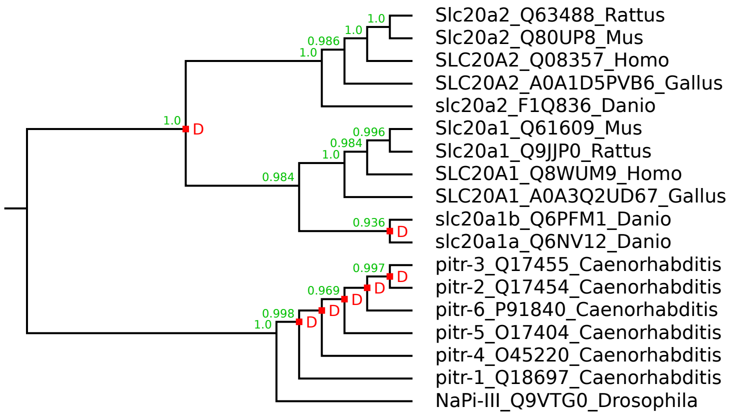

### SLC21/SLCO family

Species tree:

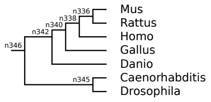


Warning: this is a large figure that had to be reduced to fit on the page.
Please use the zoom function of your PDF viewer to see the details.

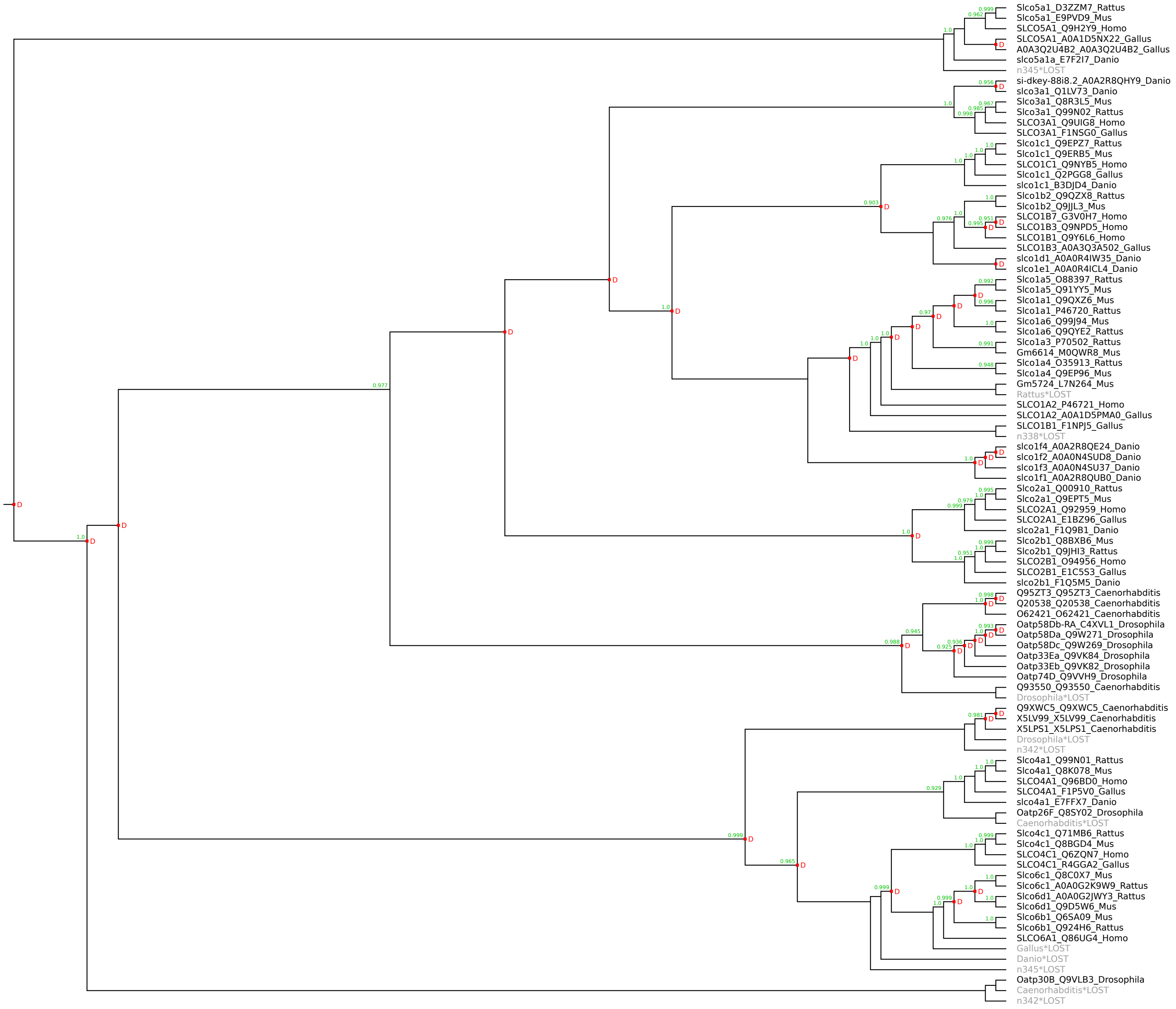

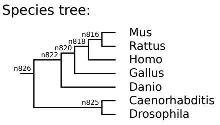


### SLC22 family

Warning: this is a large figure that had to be reduced to fit on the page.
Please use the zoom function of your PDF viewer to see the details.

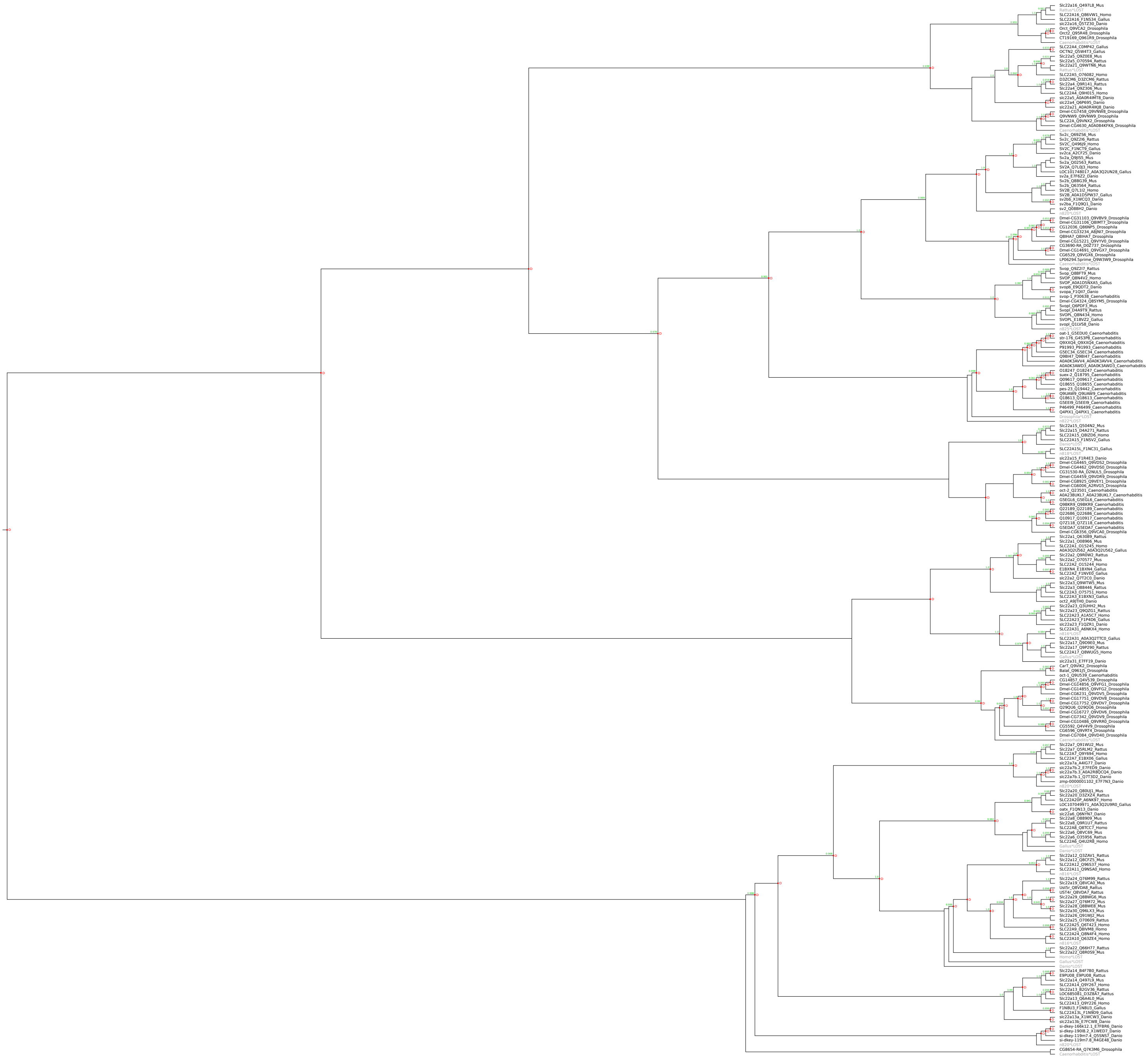

### SLC23 family

Species tree:

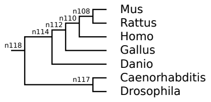


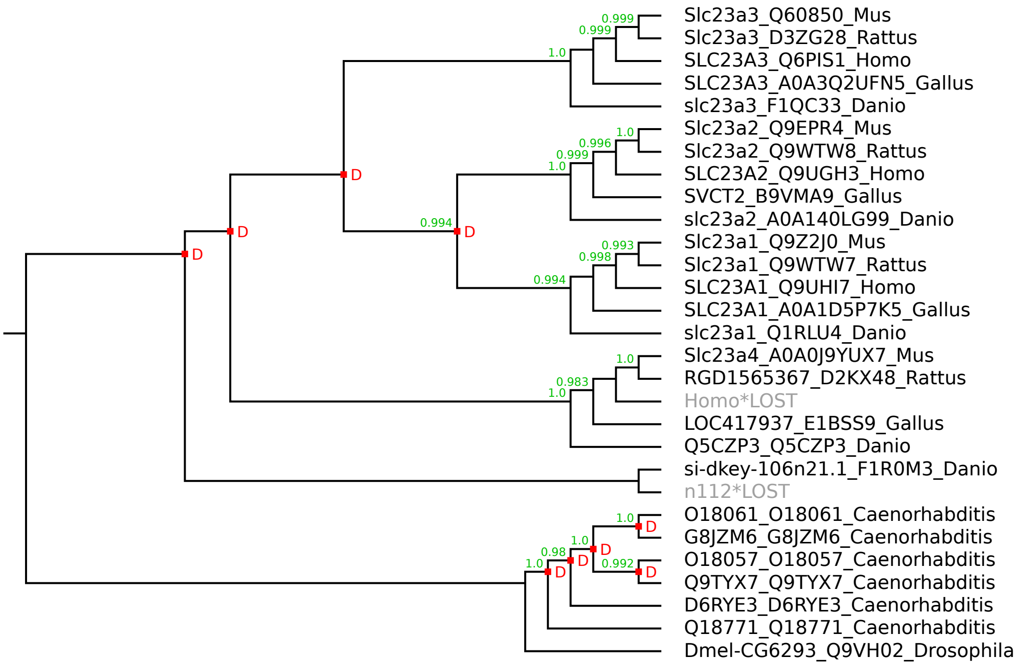

### SLC24 family

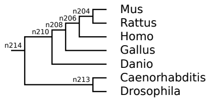


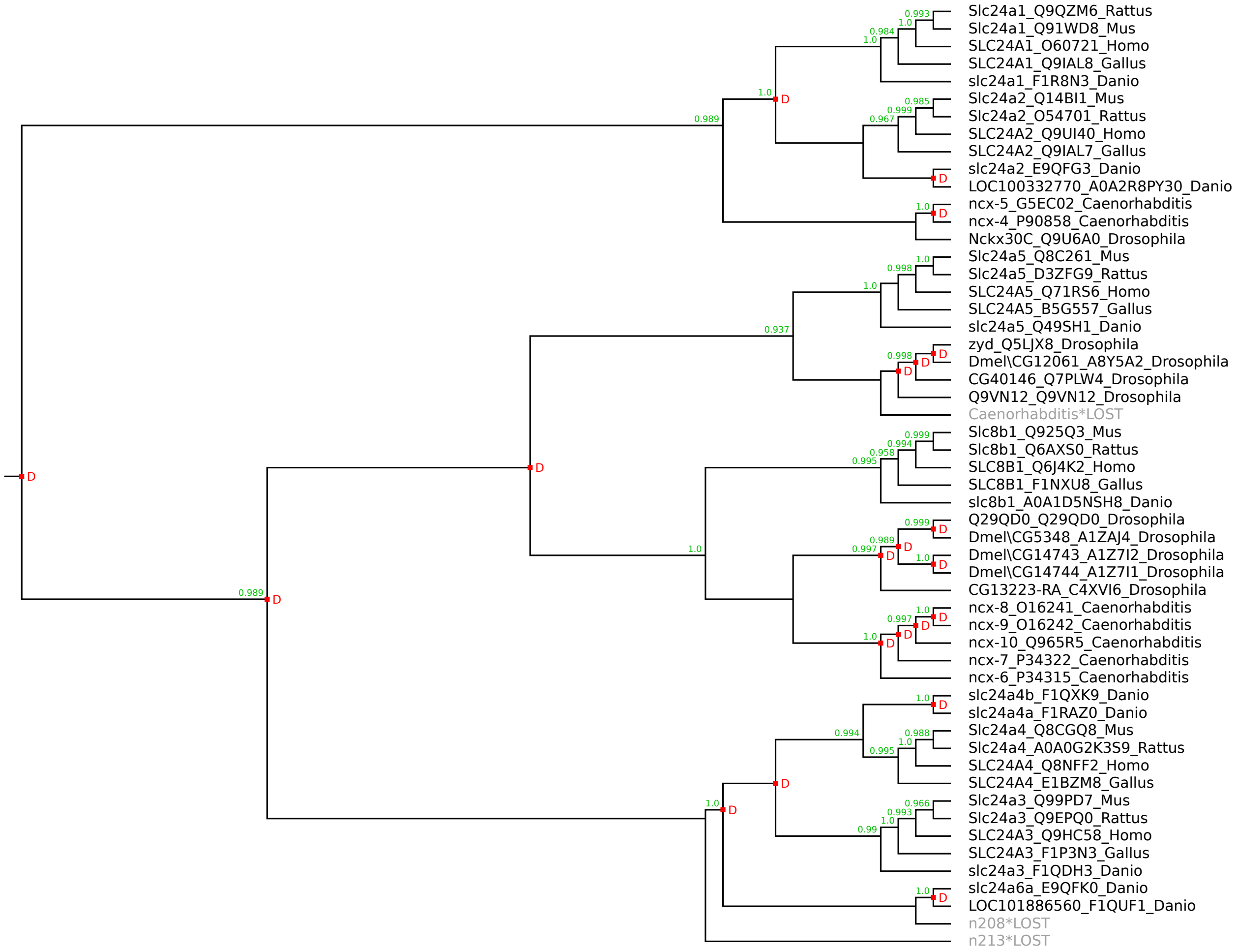

### SLC25 family

Species tree:

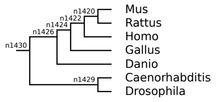


Warning: this is a large figure that had to be reduced to fit on the page.
Please use the zoom function of your PDF viewer to see the details.

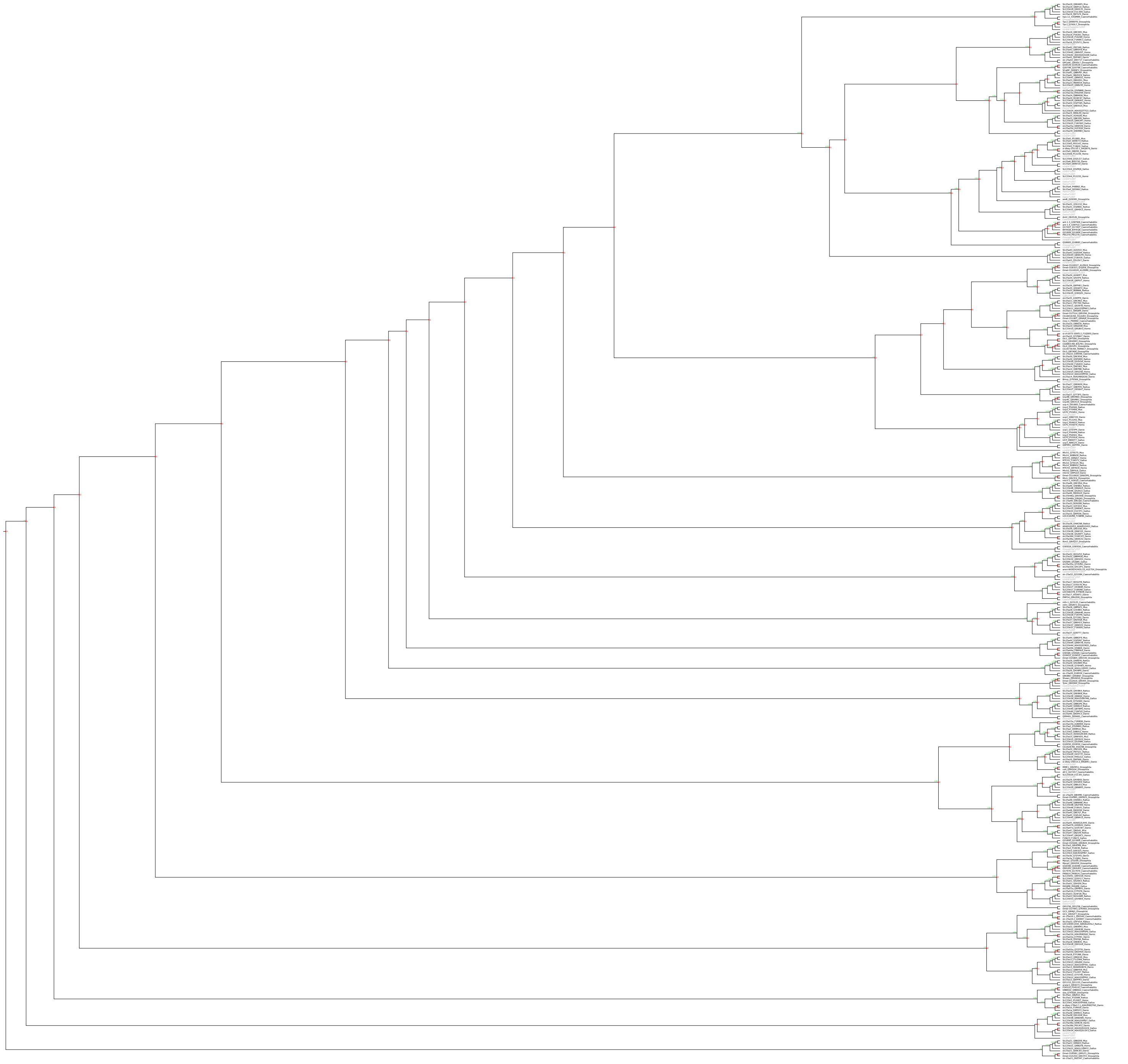

### SLC26 family

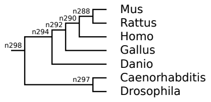


Warning: this is a large figure that had to be reduced to fit on the page.
Please use the zoom function of your PDF viewer to see the details.

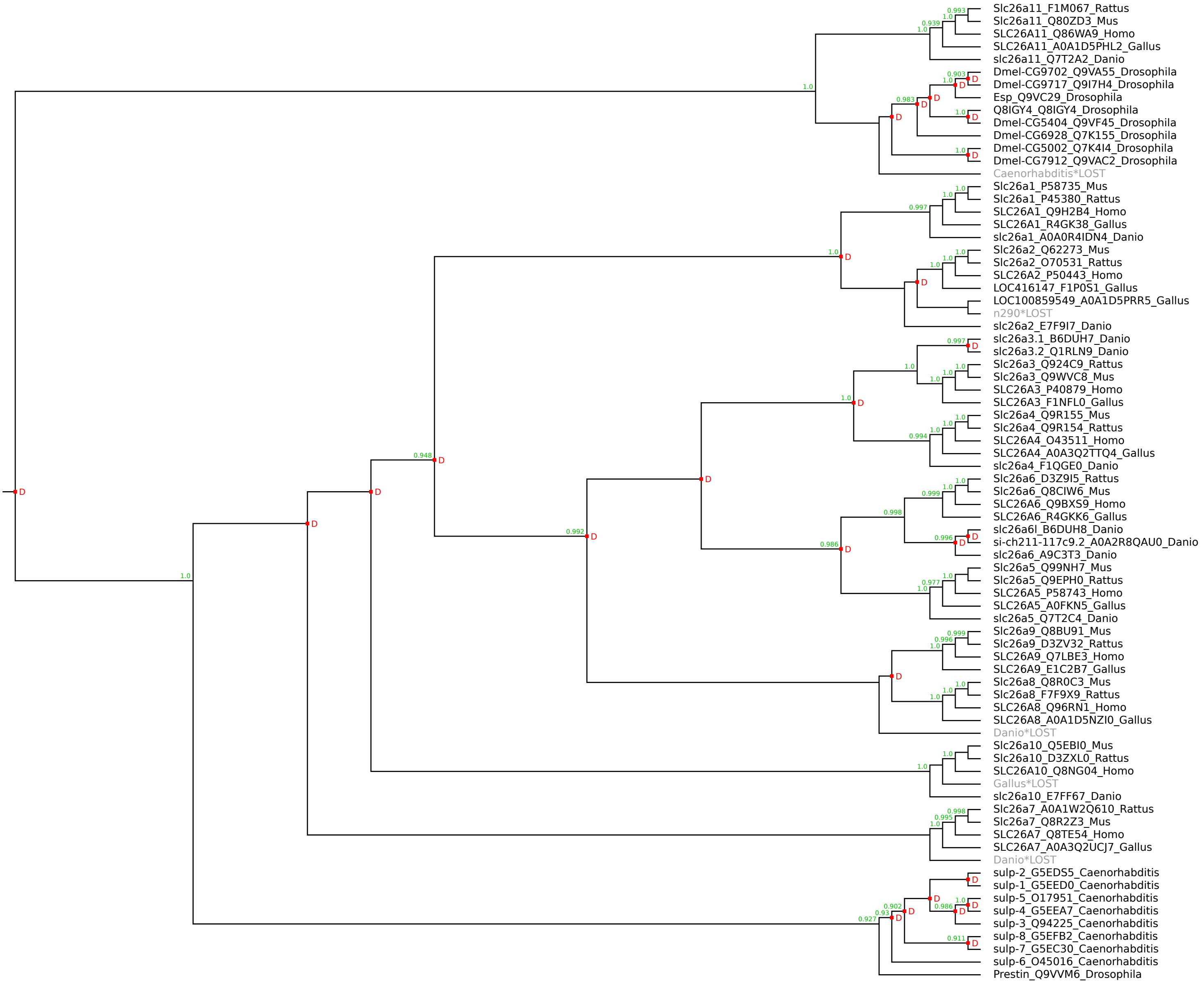

### SLC28 family

Species tree:

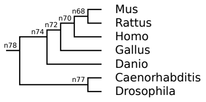


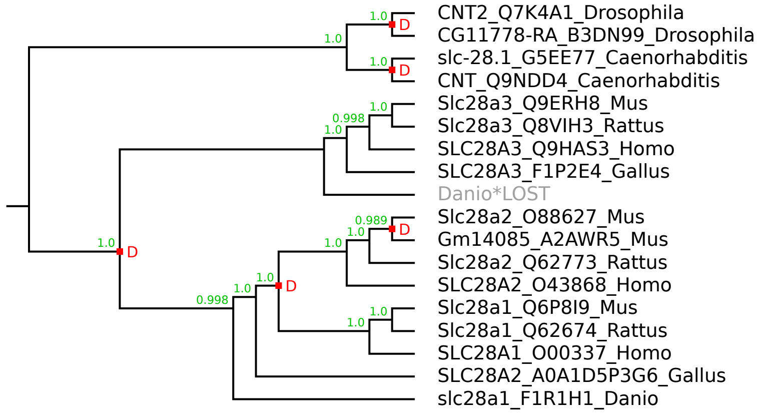

### SLC29 family

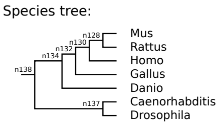


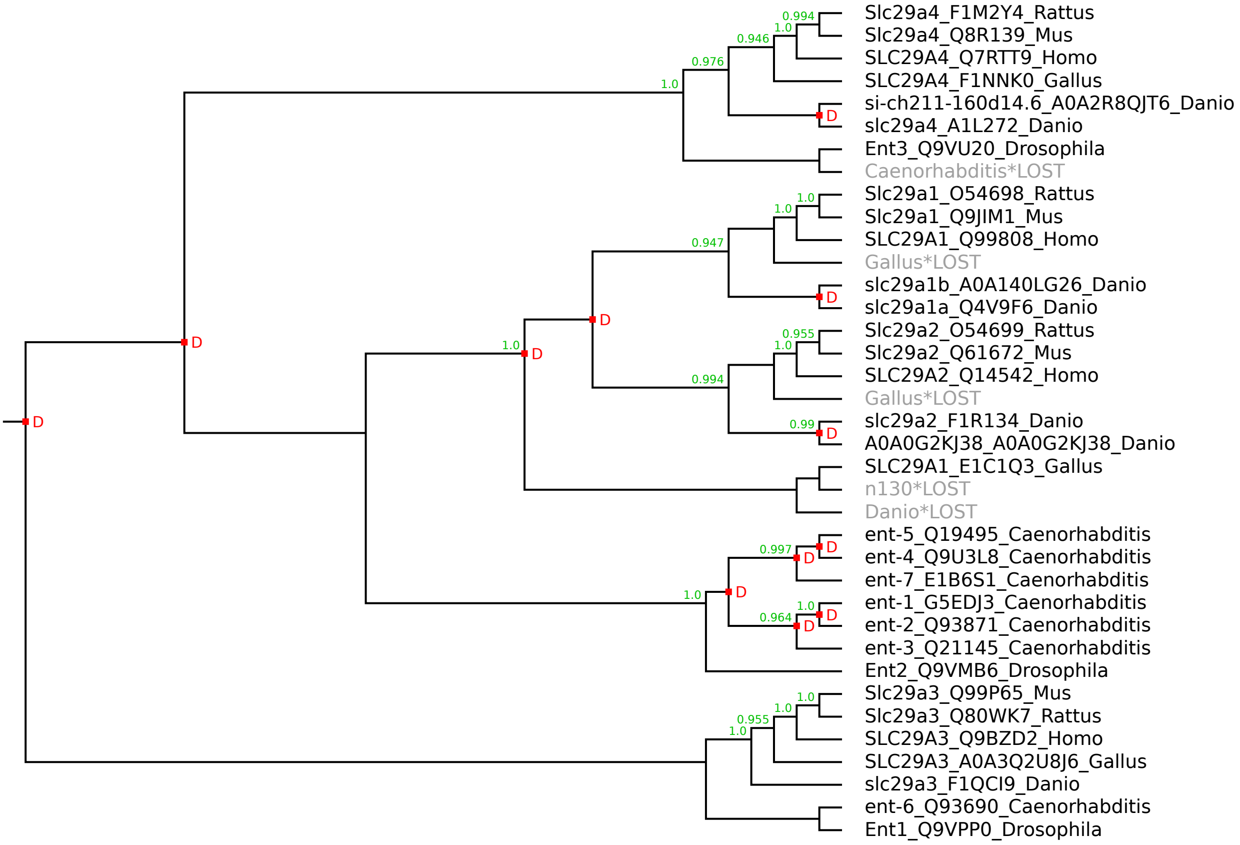

Species tree:

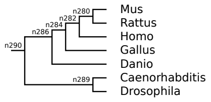


### SLC30 family

Warning: this is a large figure that had to be reduced to fit on the page.
Please use the zoom function of your PDF viewer to see the details.

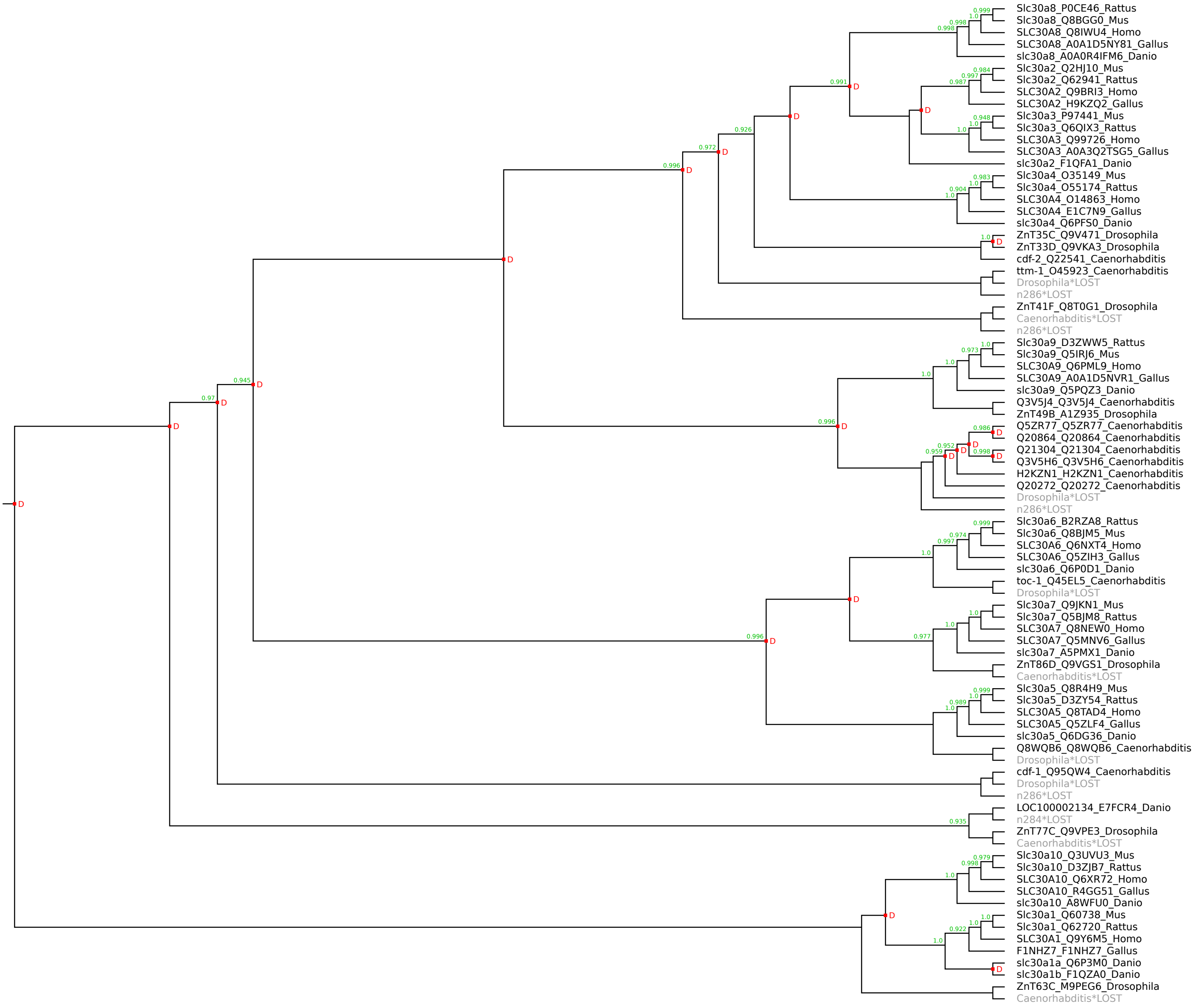

### SLC31 family

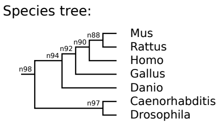


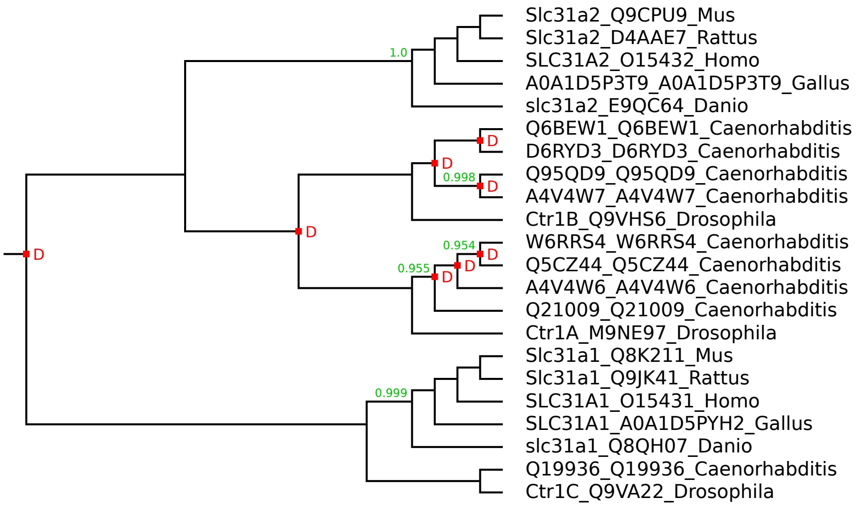

### SLC32 family

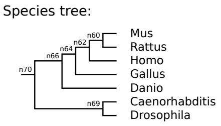


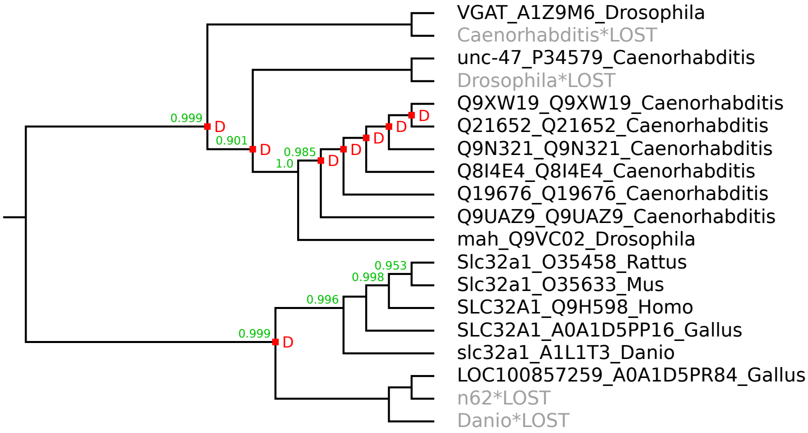

### SLC33 family

Species tree:

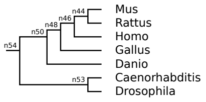


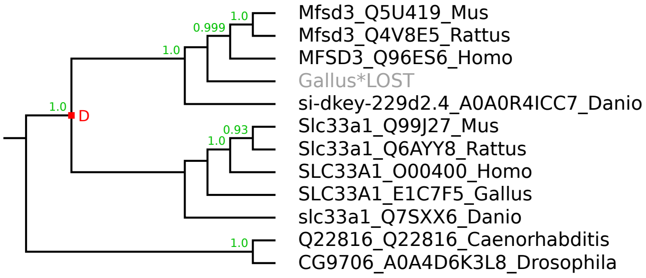

### SLC34 family

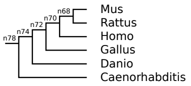


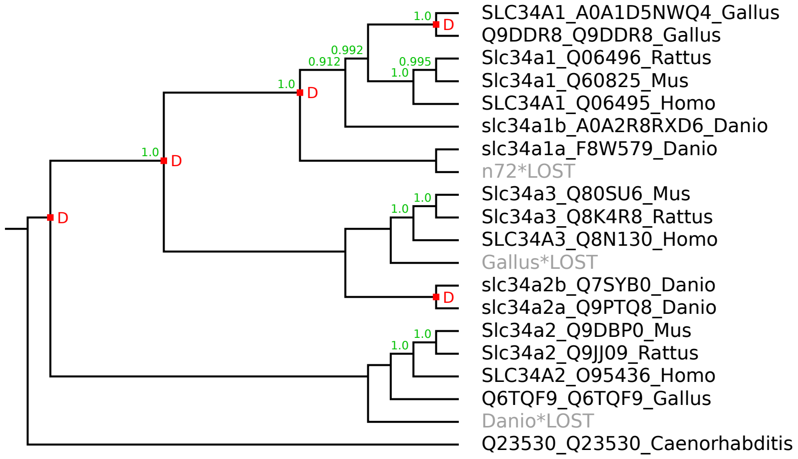

Species tree:

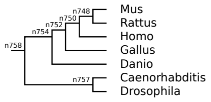


### SLC35 family

Warning: this is a large figure that had to be reduced to fit on the page.
Please use the zoom function of your PDF viewer to see the details.

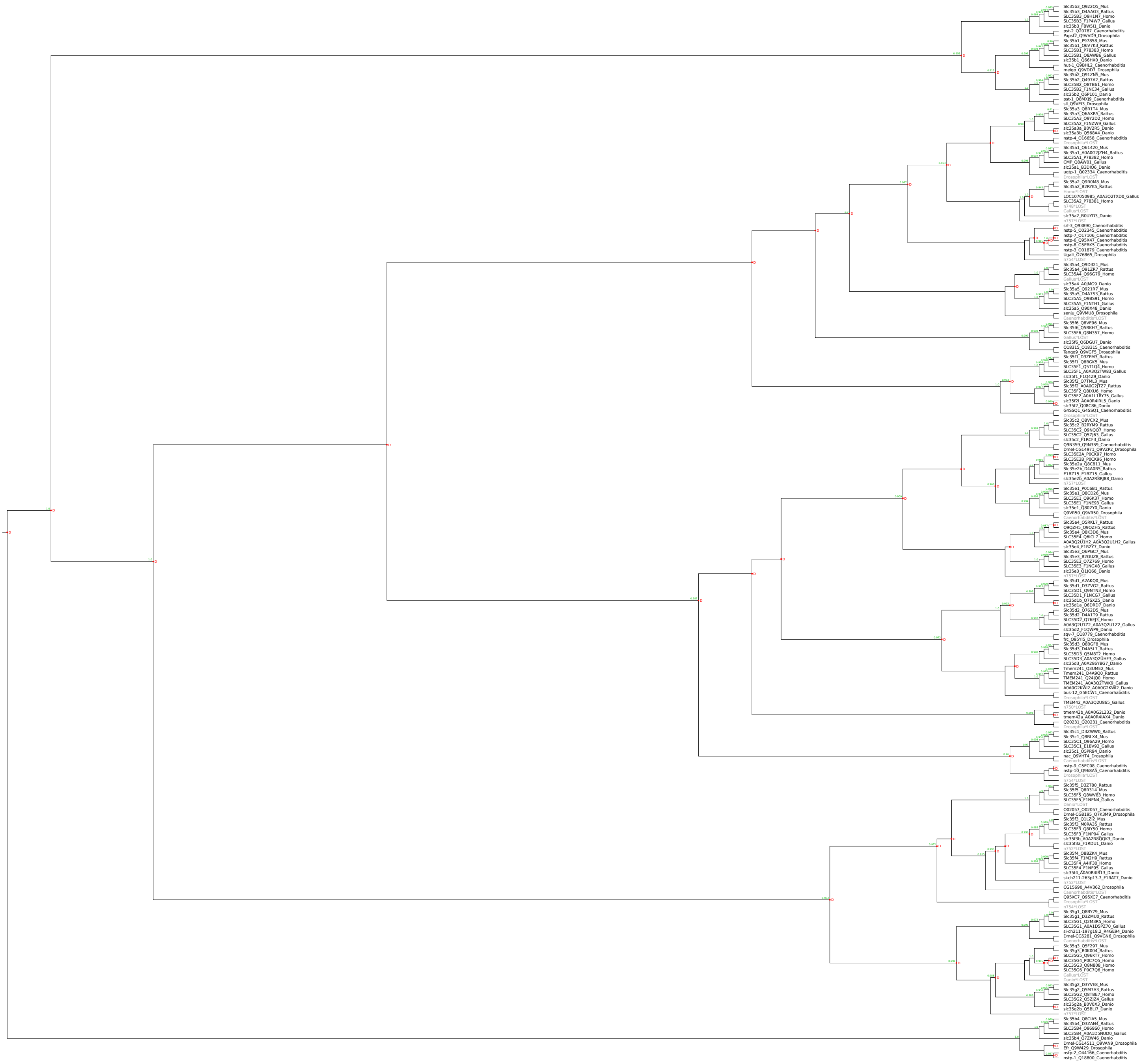

### SLC36 family

Species tree:

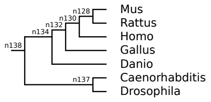


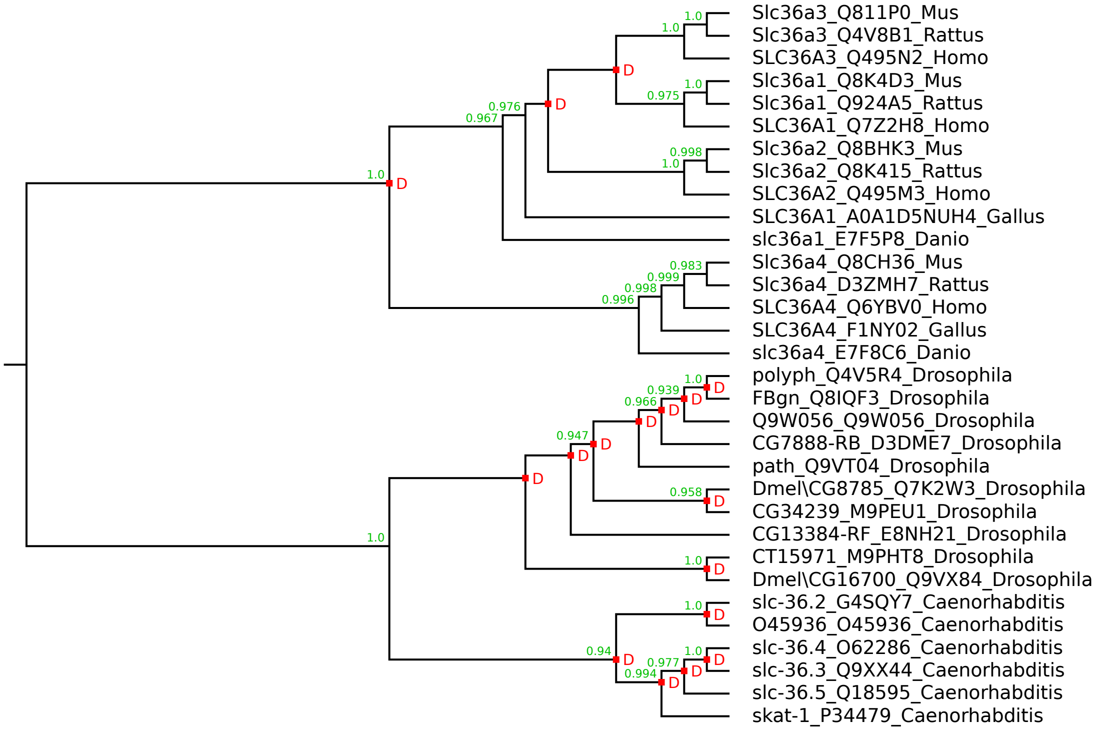

### SLC37 family

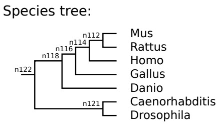


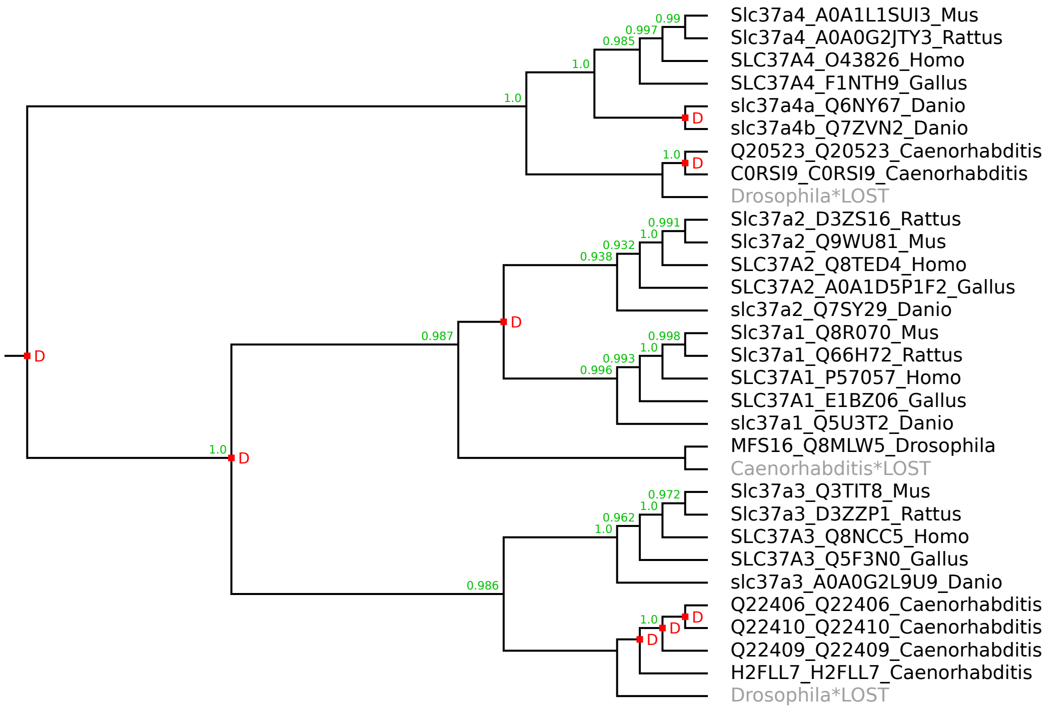

### SLC38 family

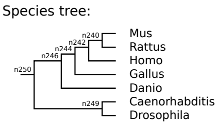


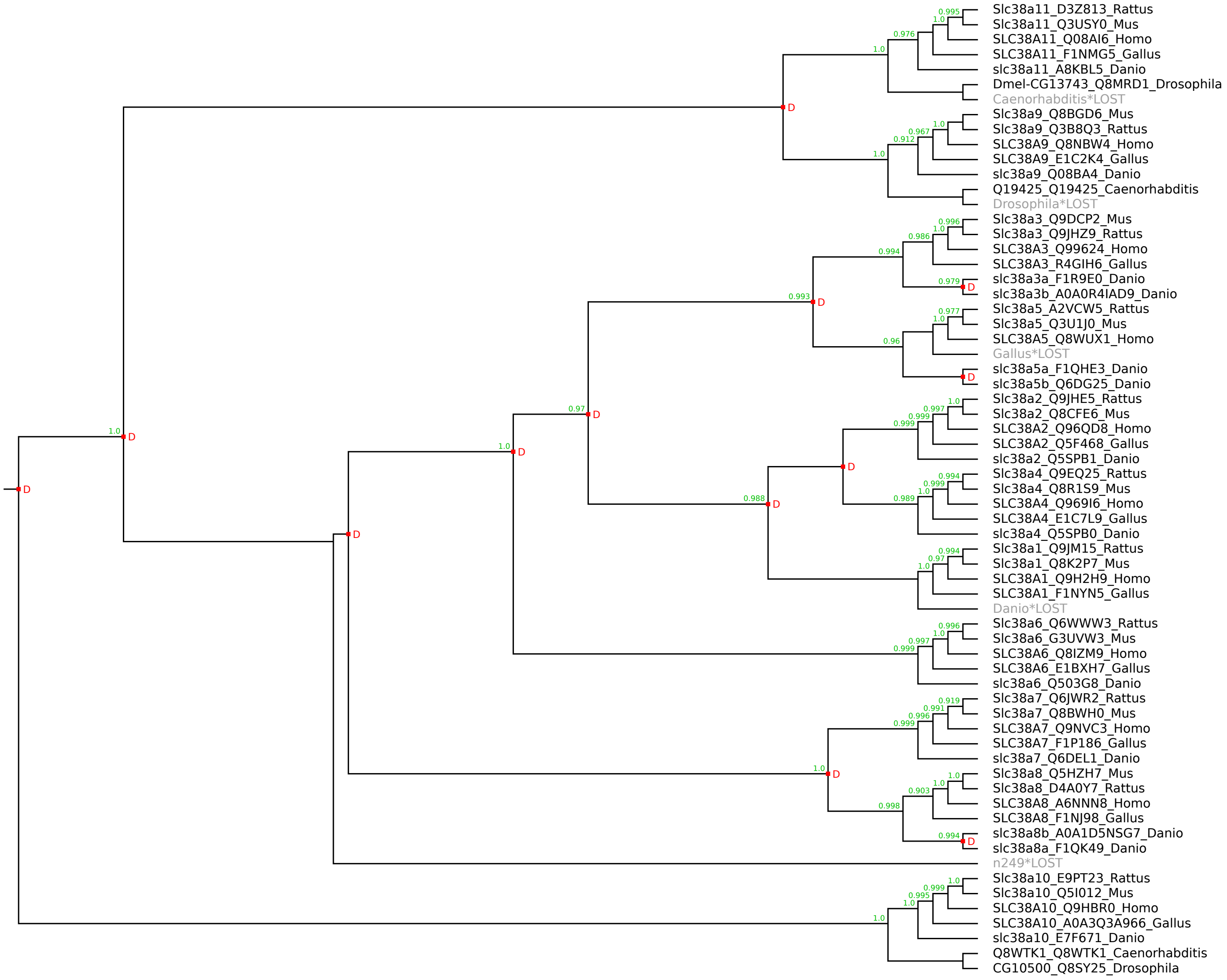

Species tree:

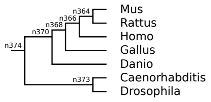


### SLC39 family

Warning: this is a large figure that had to be reduced to fit on the page.
Please use the zoom function of your PDF viewer to see the details.

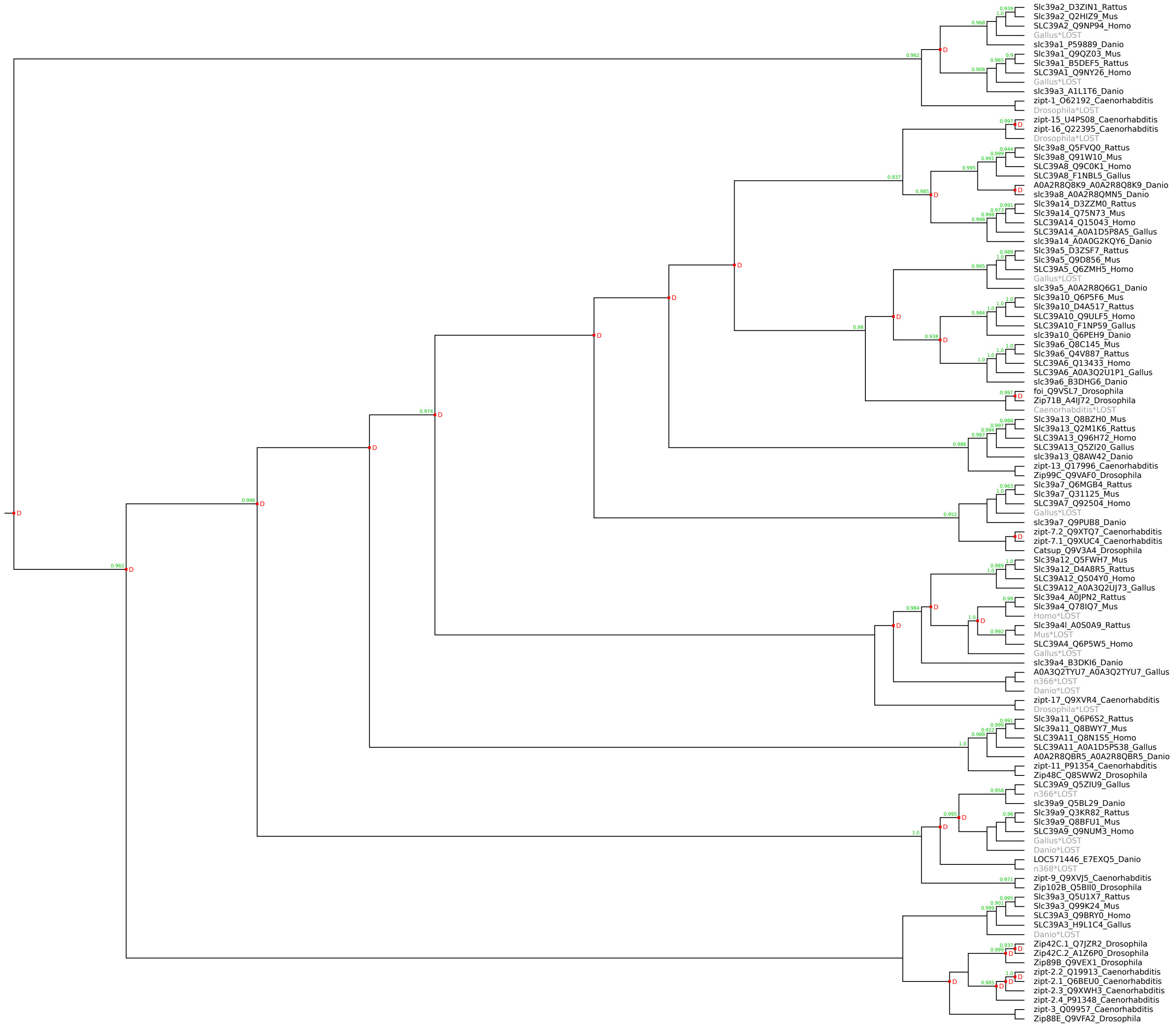

### SLC40 family

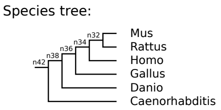


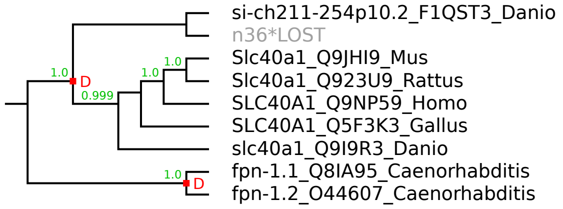

### SLC41 family

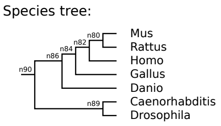


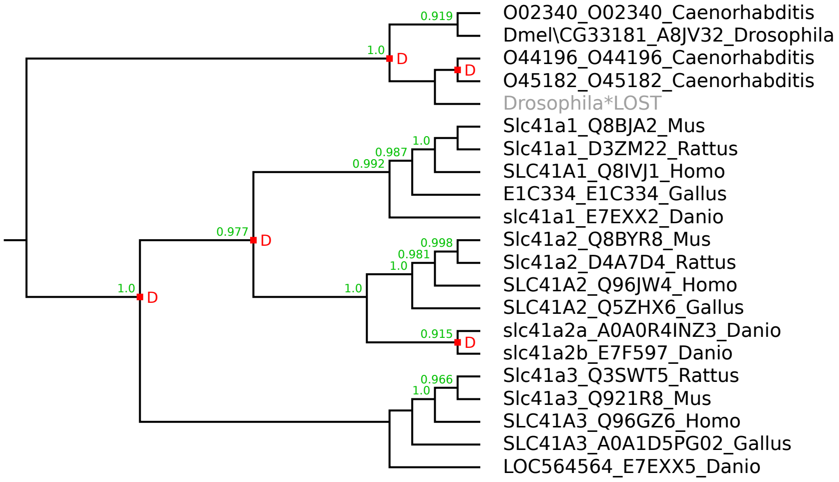

### SLC42 family

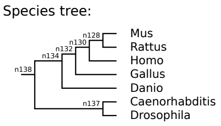


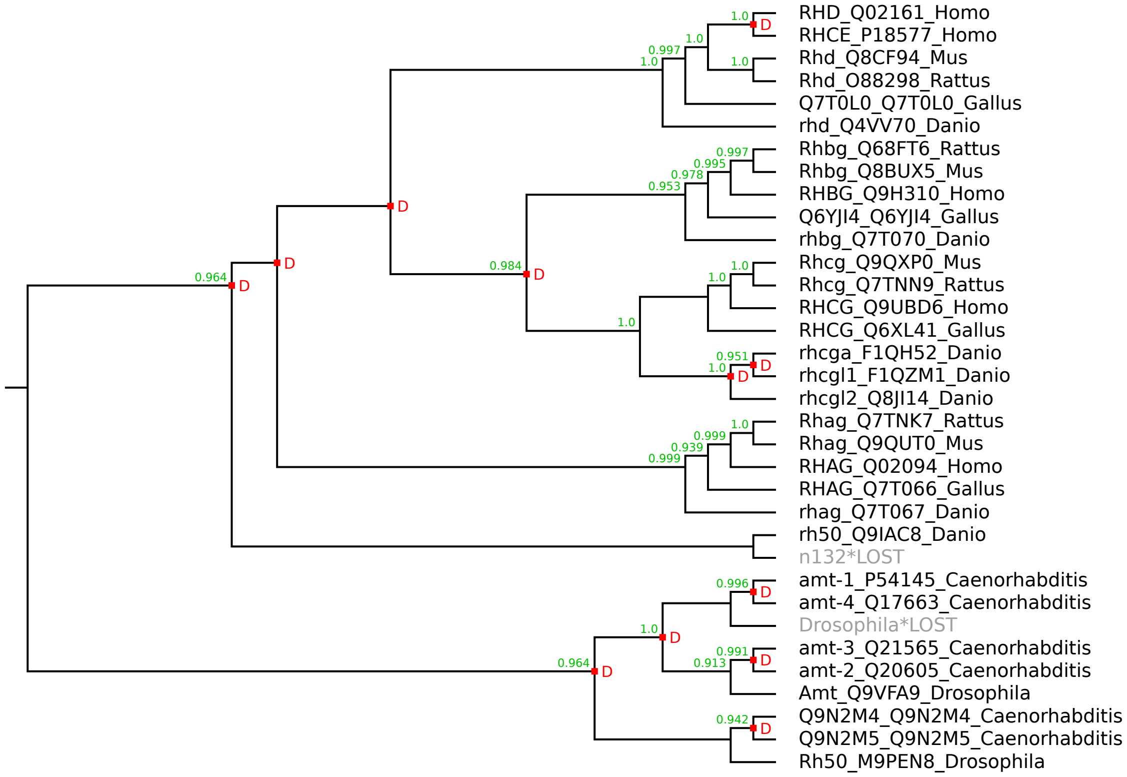

### SLC43 family

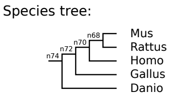


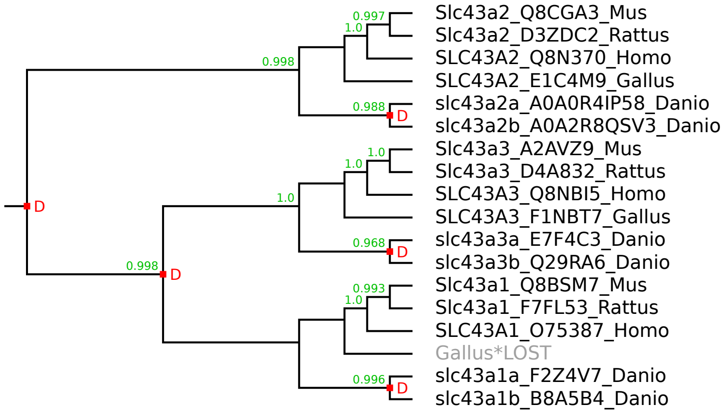

### SLC44 family

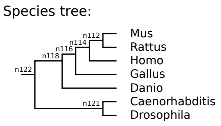


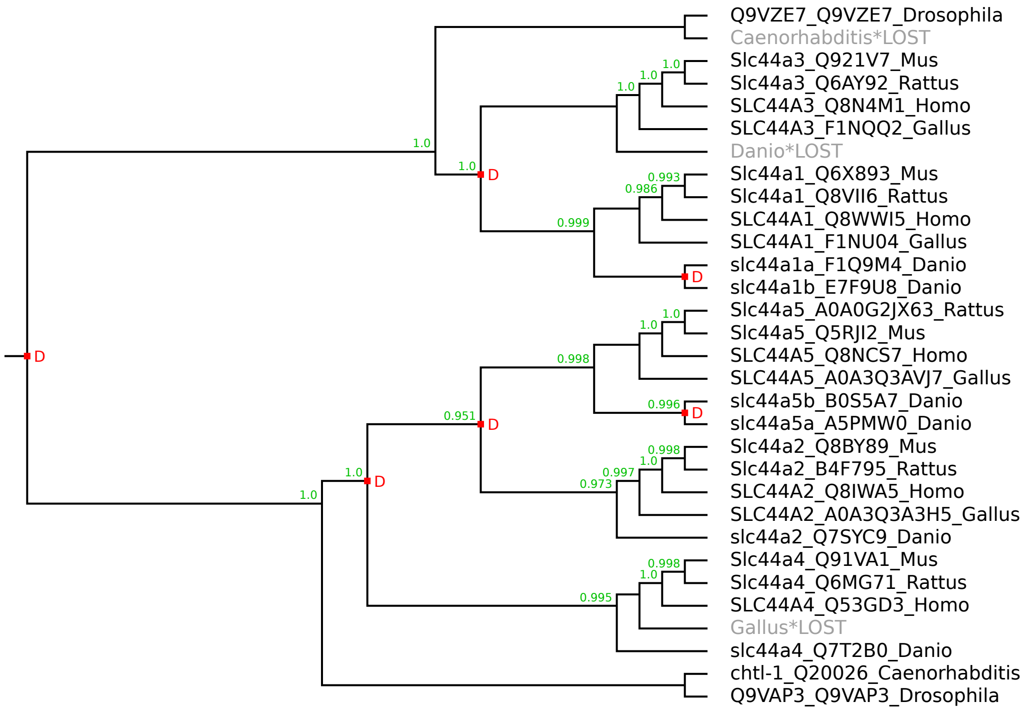

### SLC45 family

Species tree:

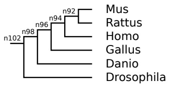


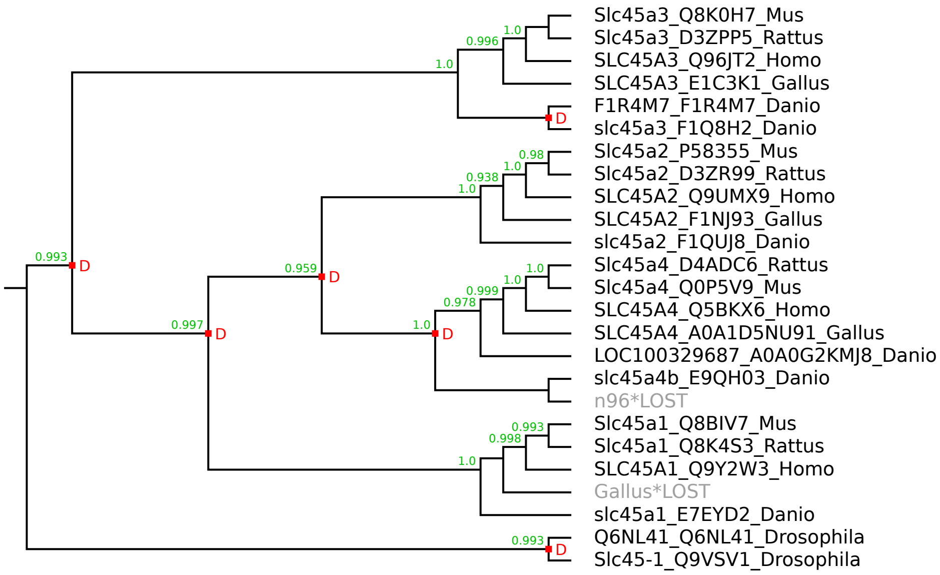

### SLC46 family

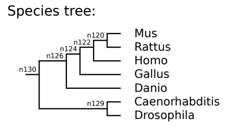


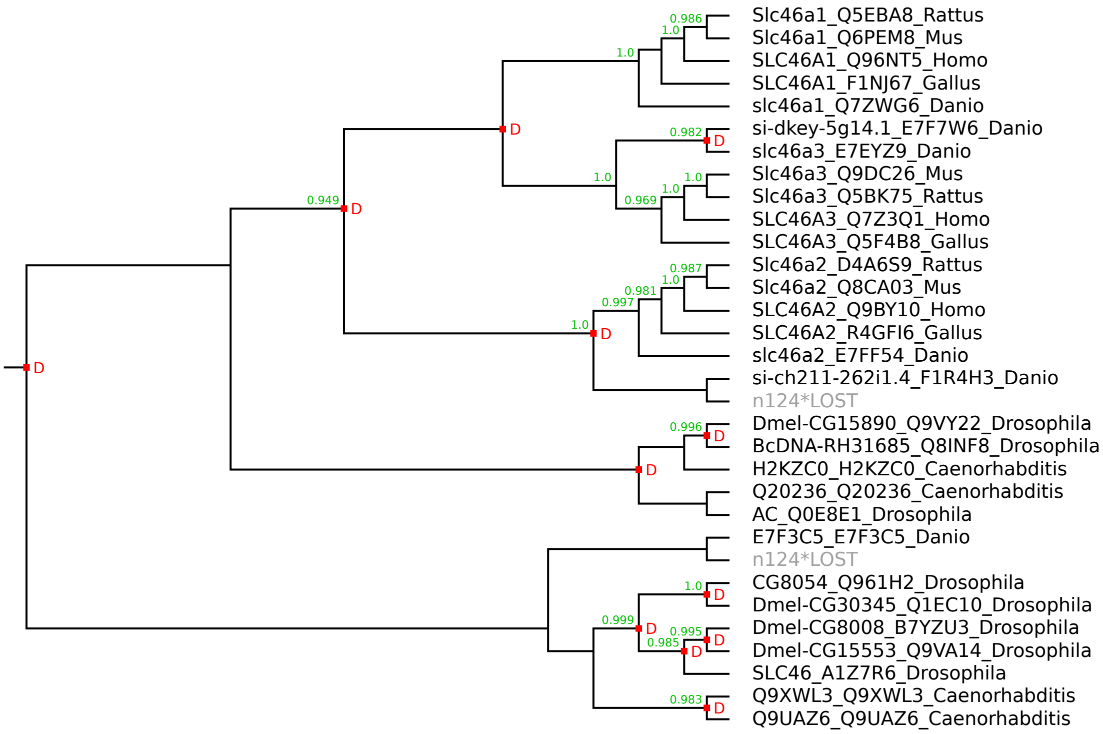

### SLC47 family

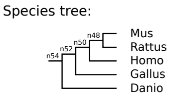


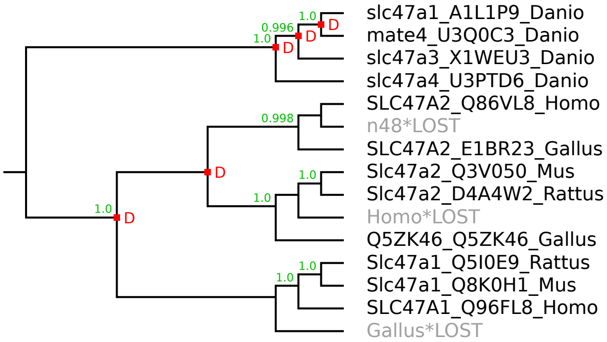

### SLC48 family

Species tree:

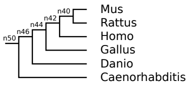


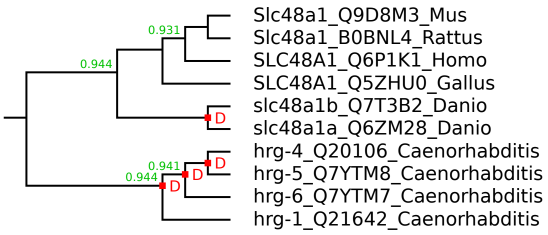

### SLC49 family

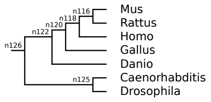


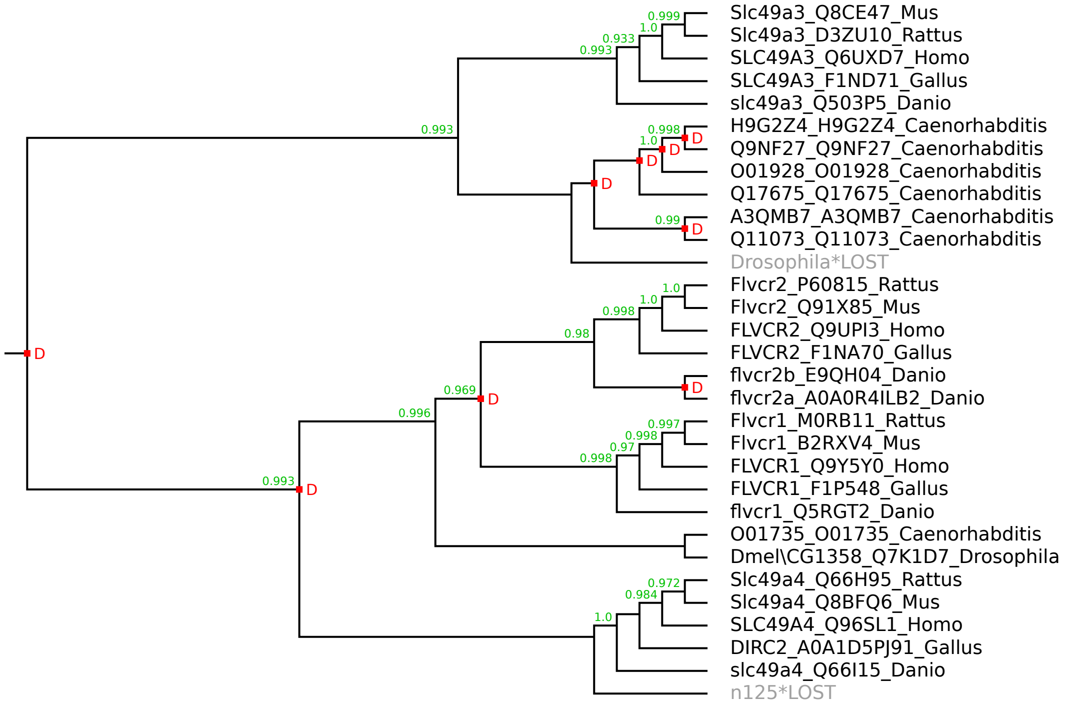

### SLC50 family

Species tree:

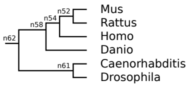


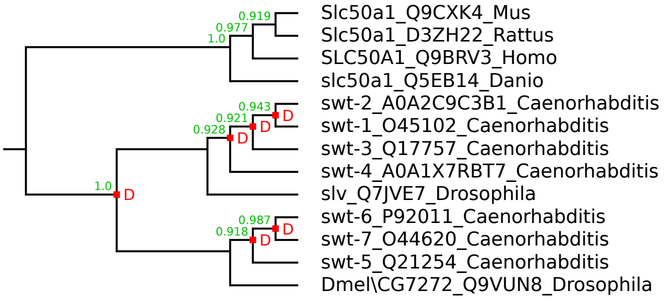

### SLC51A family

Species tree:

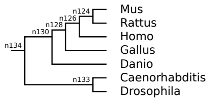


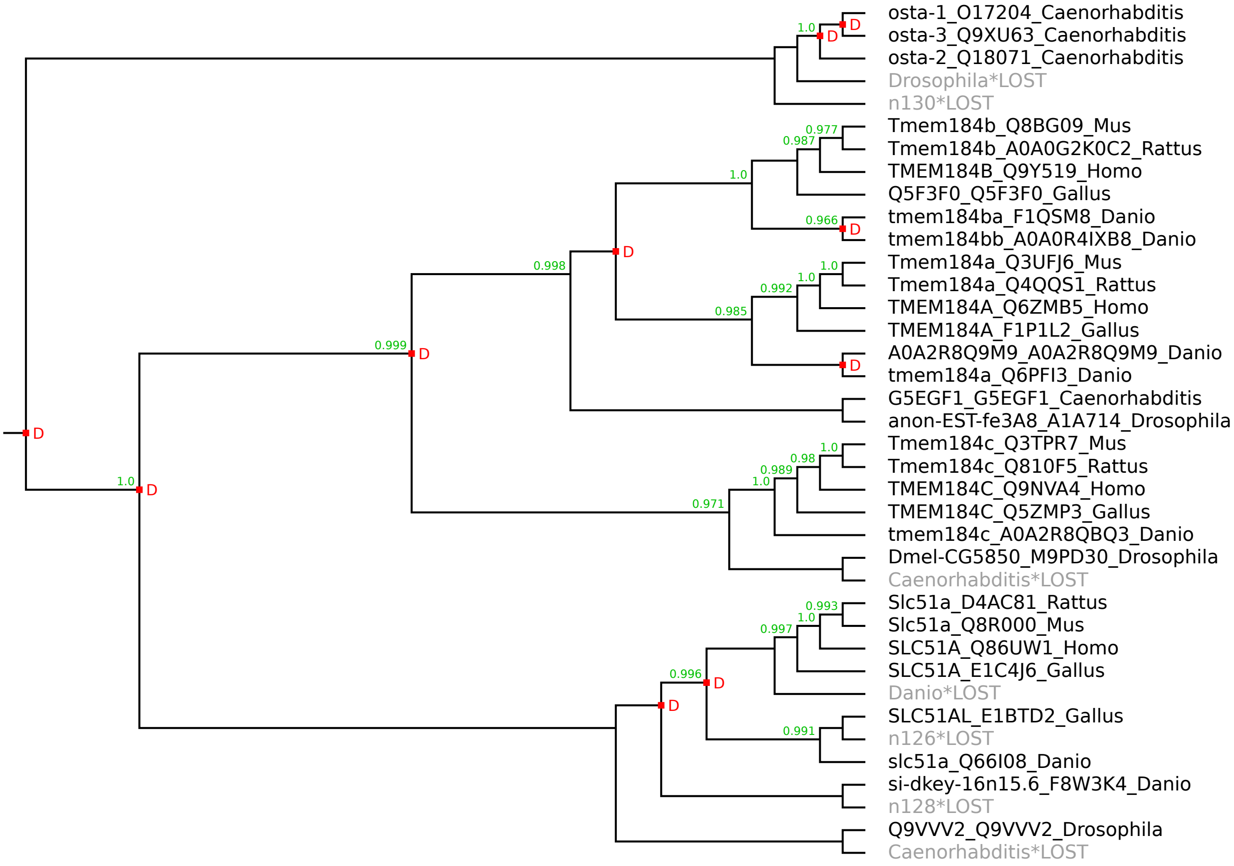

Species tree:

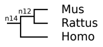


### SLC51B family

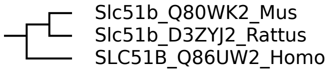

### SLC52 family

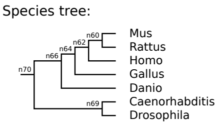


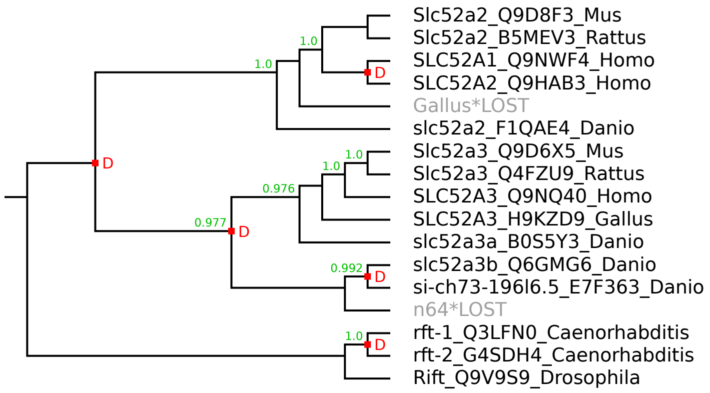

### SLC53 family

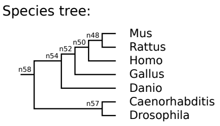


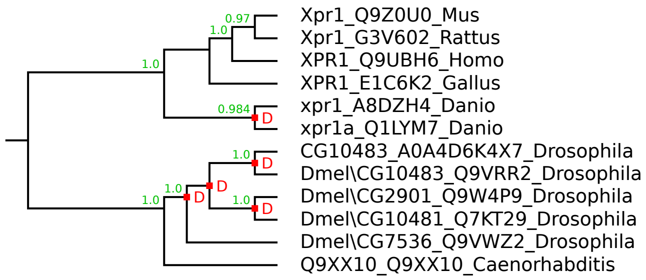

### SLC54 family

Species tree:

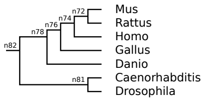


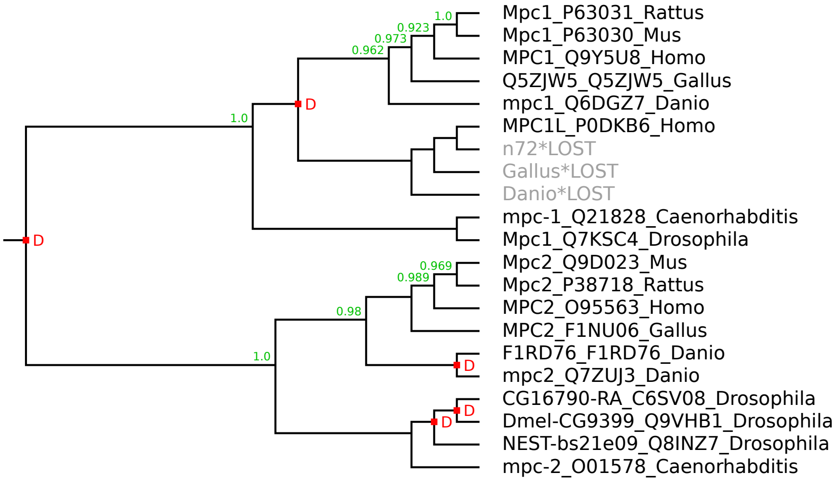

### SLC55 family

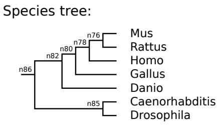


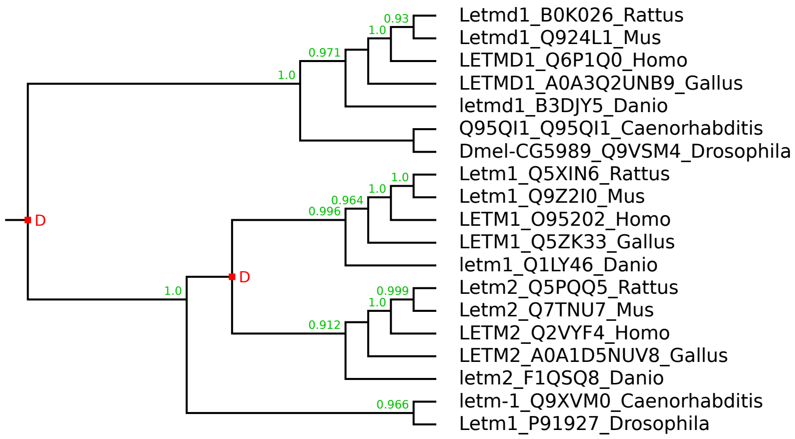

### SLC56 family

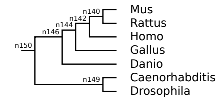


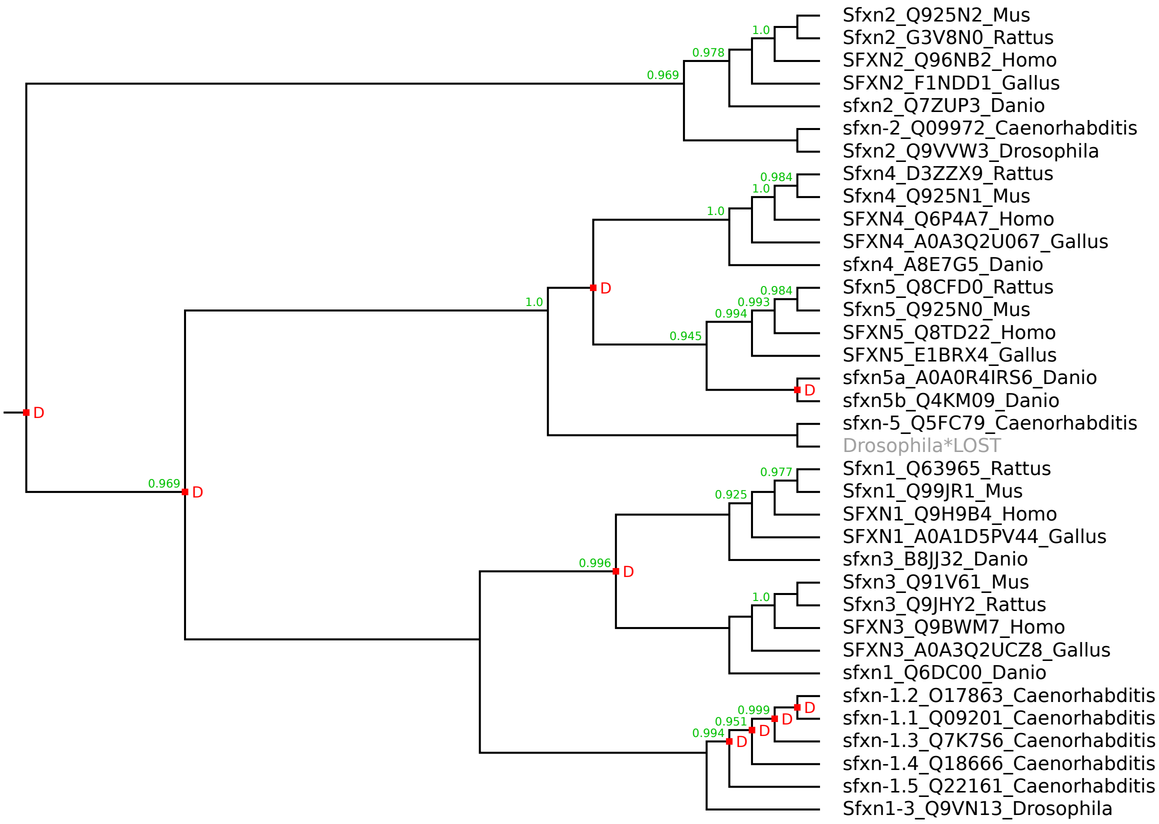

### SLC57 family

Species tree:

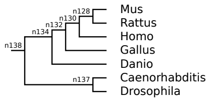


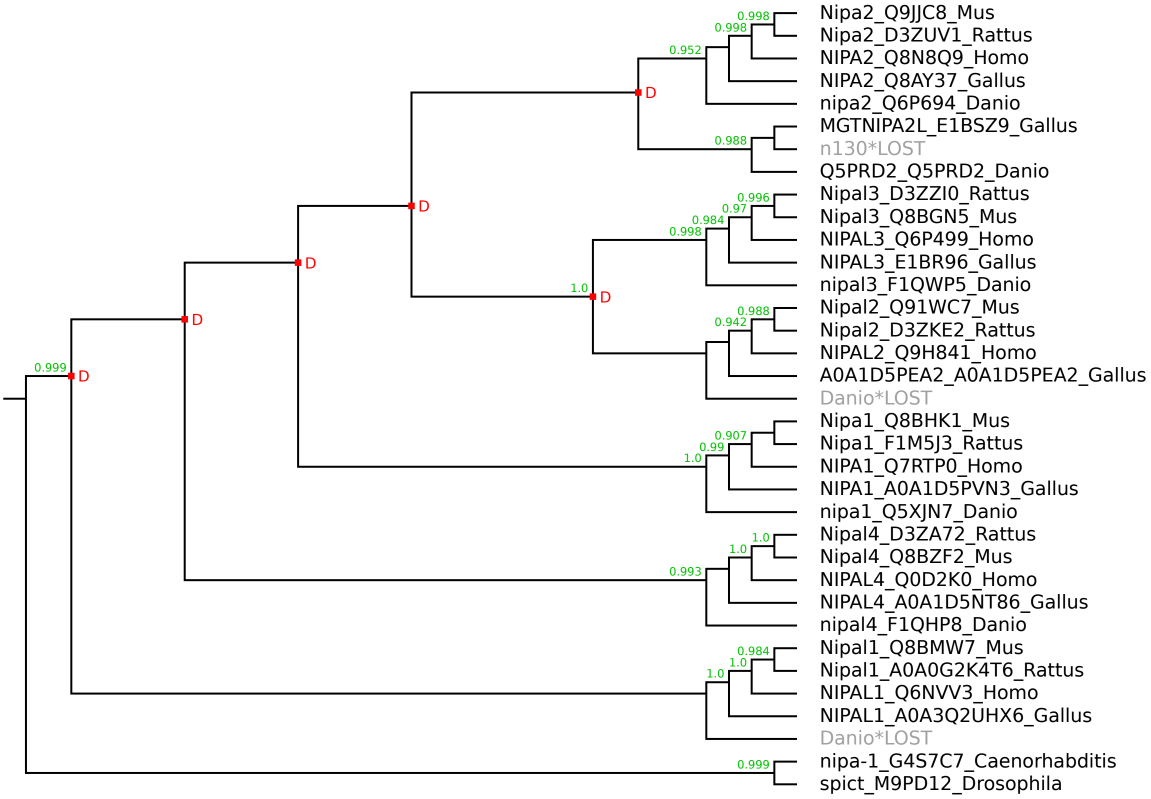

### SLC58 family

Species tree:

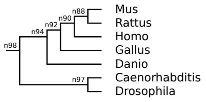


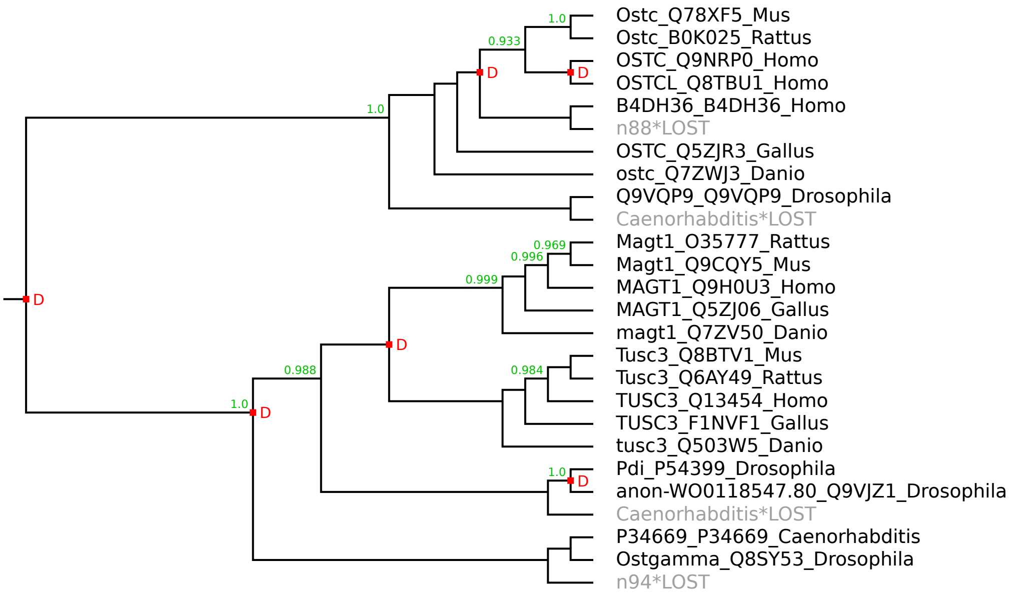

### SLC59 family

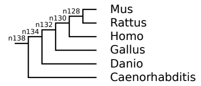


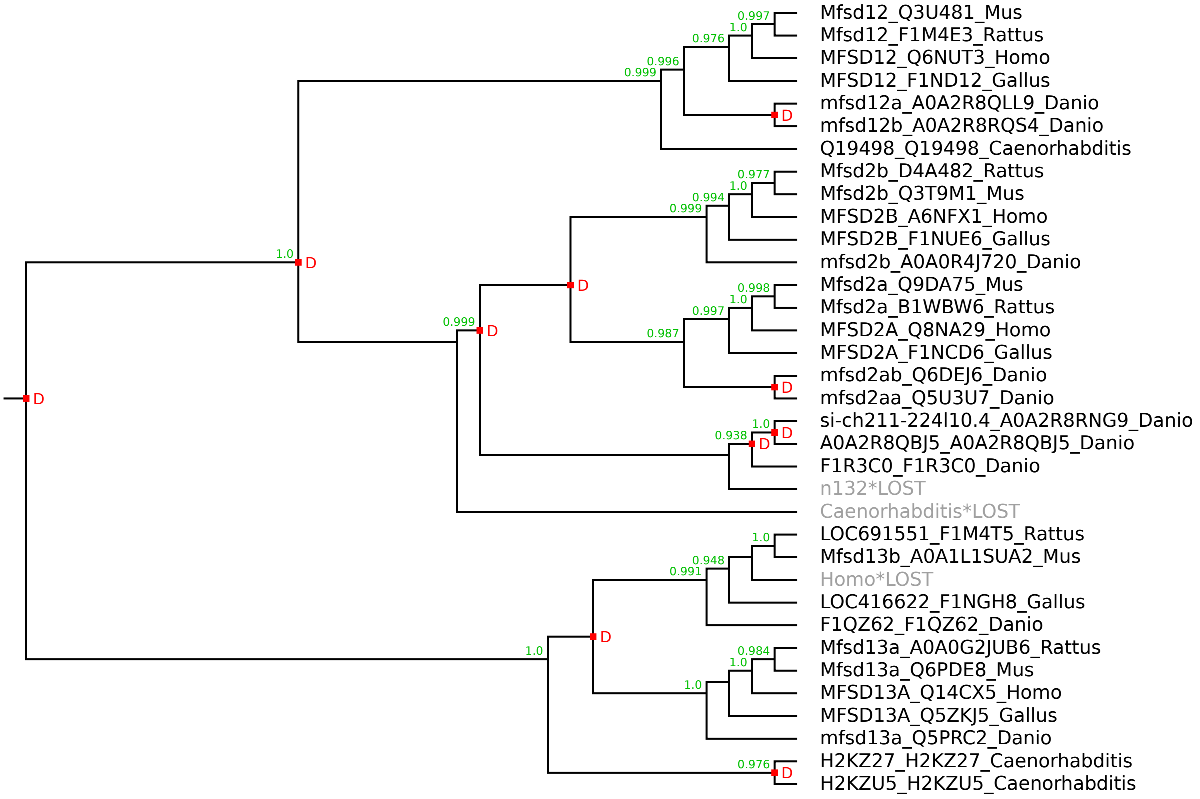

### SLC60 family

Species tree:

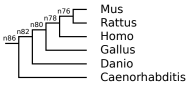


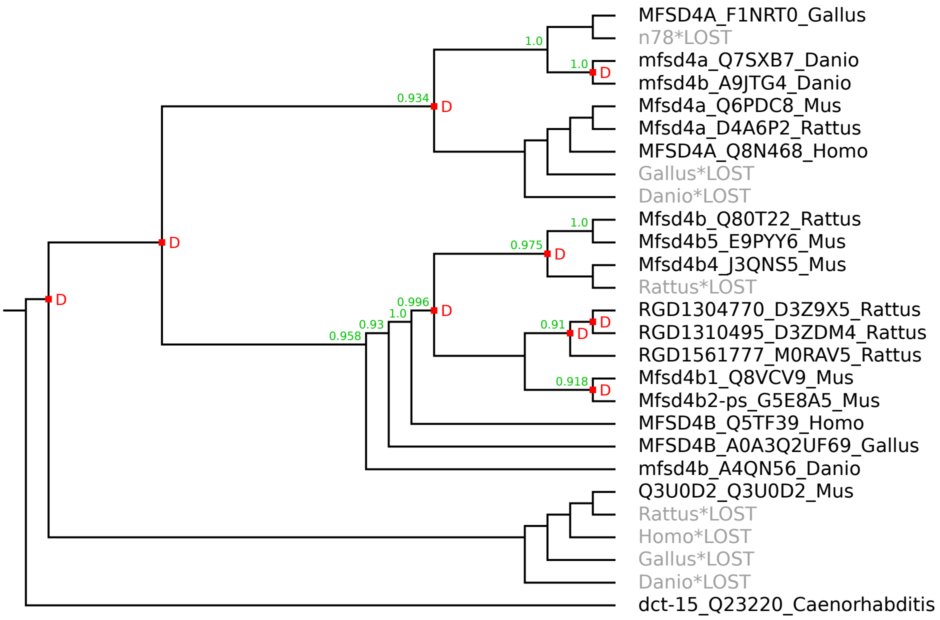

### SLC61 family

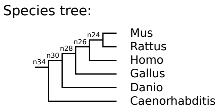


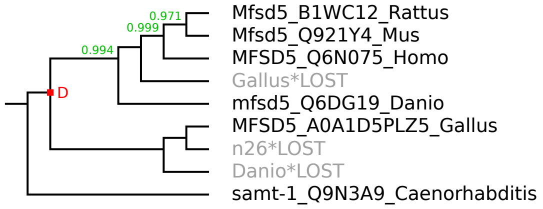

### SLC62 family

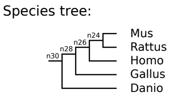


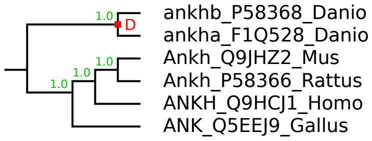

### SLC63 family

Species tree:

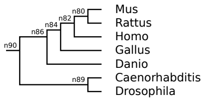


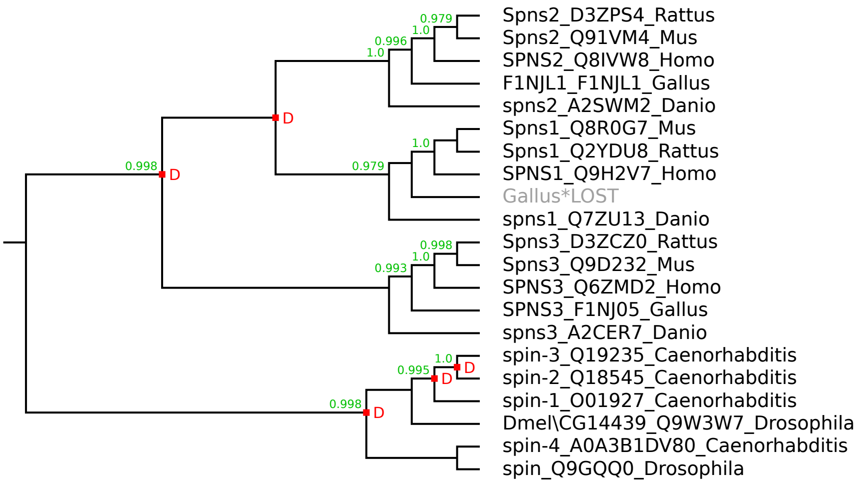

### SLC64 family

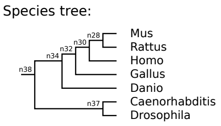


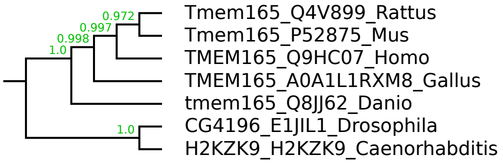

### SLC65 family

Species tree:

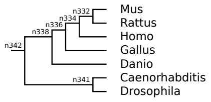


---

Warning: this is a large figure that had to be reduced to fit on the page.
Please use the zoom function of your PDF viewer to see the details.

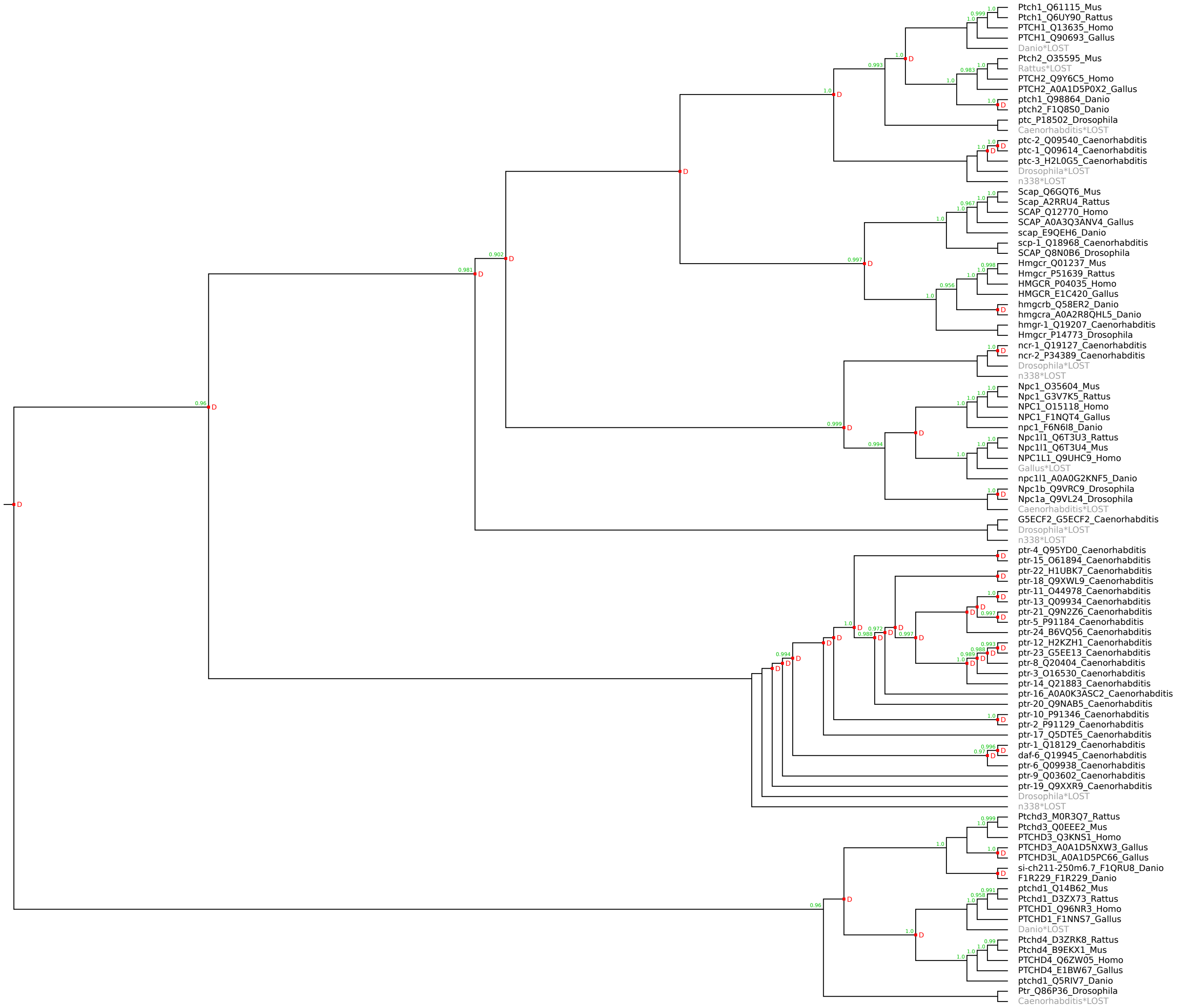

### SLC66 family

Species tree:

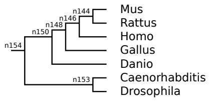


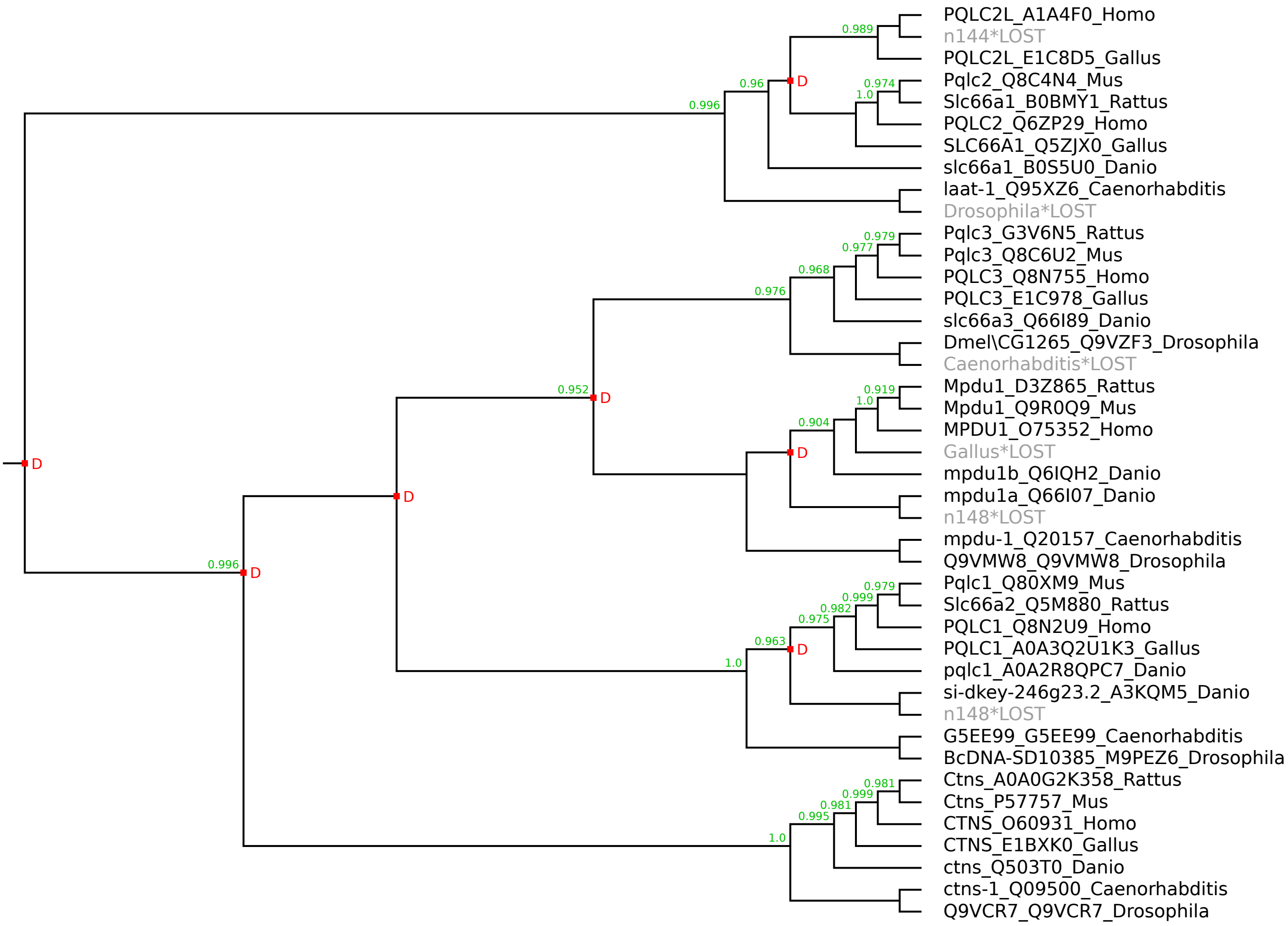

Species tree:

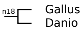


### pSLC.5269 family

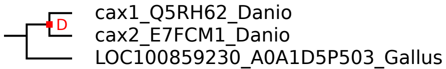

### pSLC.7224 family

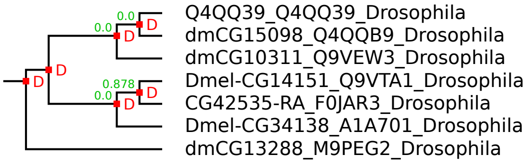

### pSLC.ARV1 family

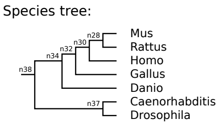


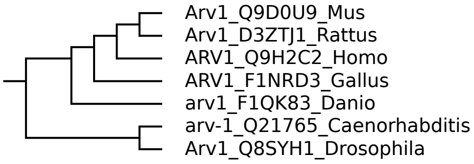

### pSLC.Battenin family

Species tree:

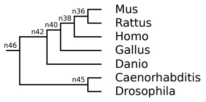


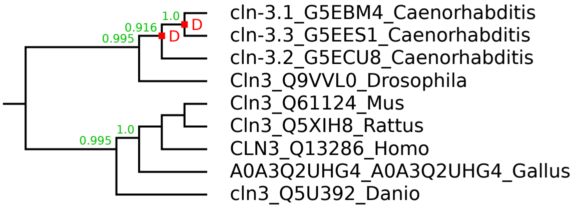

### pSLC.CLCN family

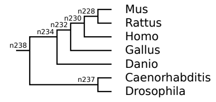


Warning: this is a large figure that had to be reduced to fit on the page.
Please use the zoom function of your PDF viewer to see the details.

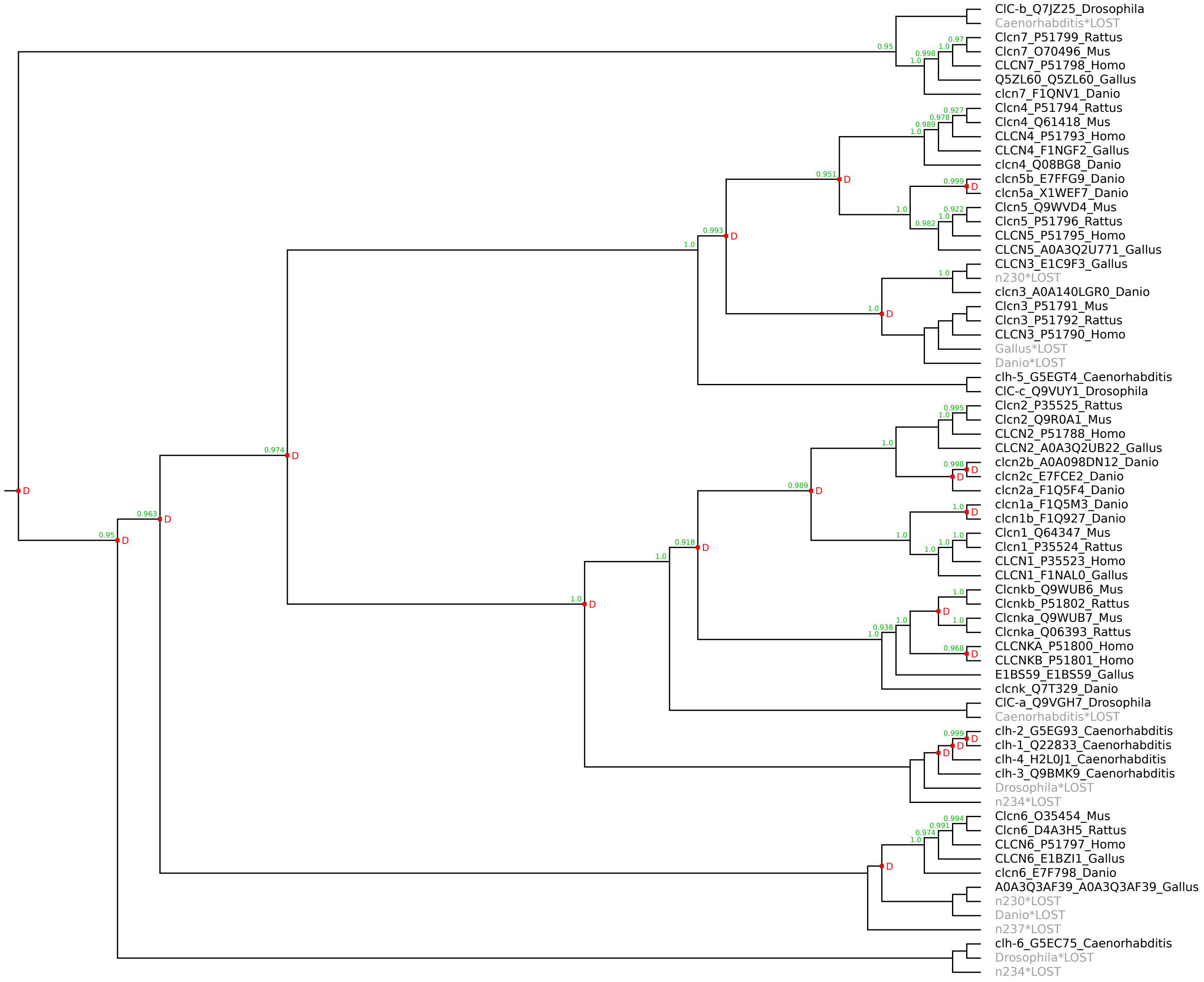

### pSLC.CNNM family

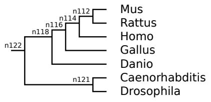


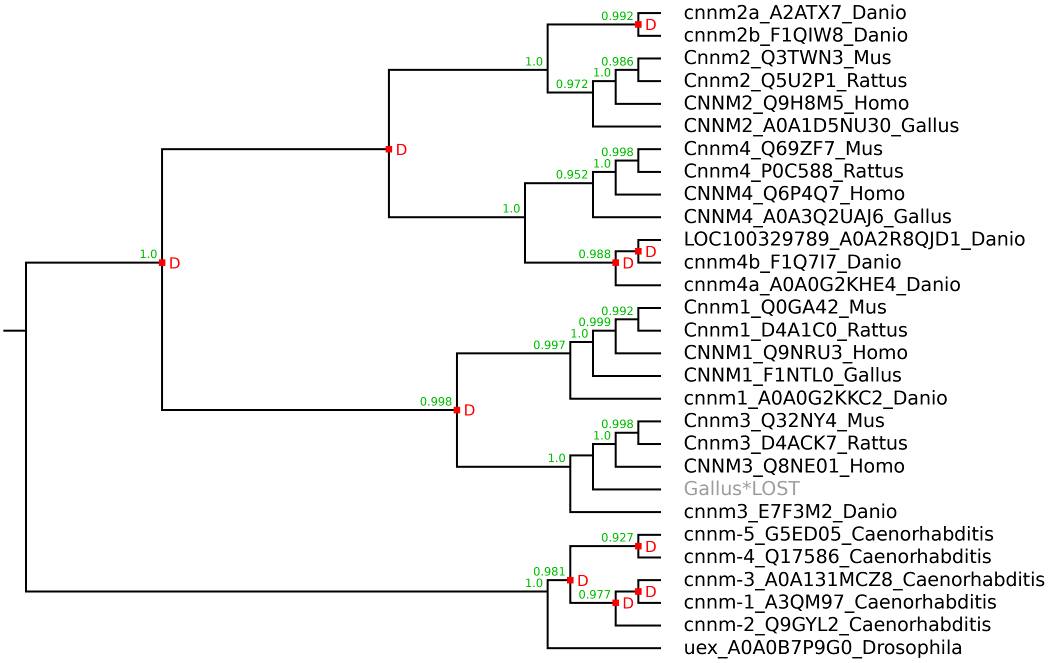

### pSLC.CitMHS family

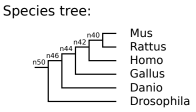


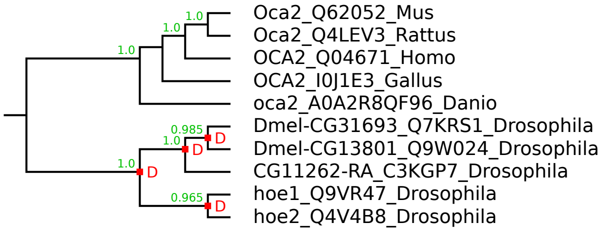

### pSLC.Dispatched family

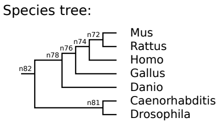


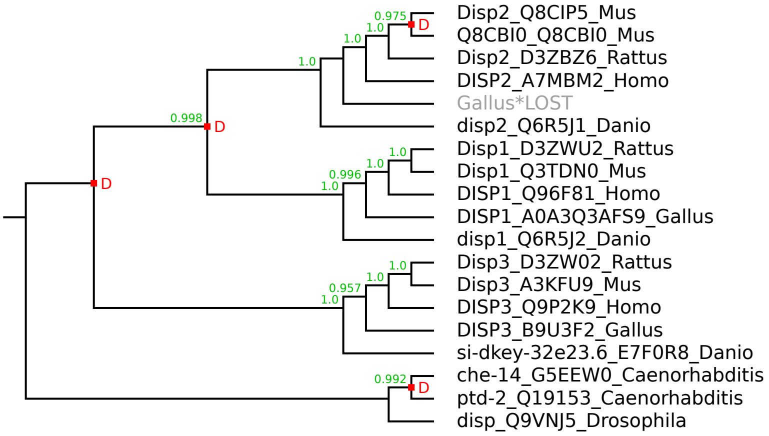

### pSLC.GPR155 family

Species tree:

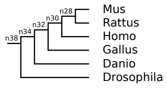


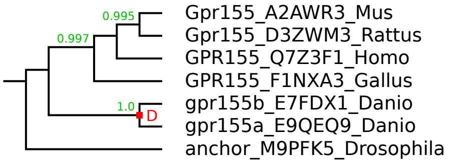

### pSLC.LAPTM family

Species tree:

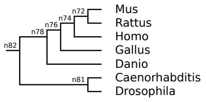


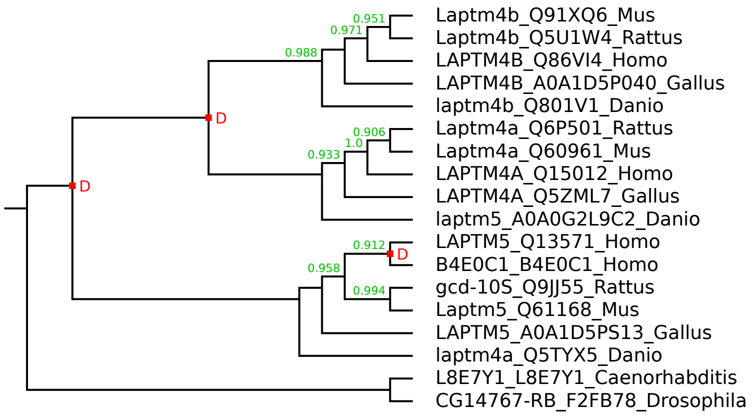

### pSLC.LMBR-A family

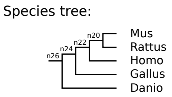


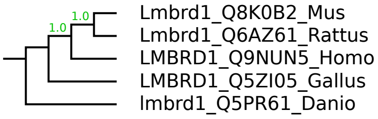

### pSLC.LMBR-B family

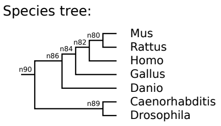


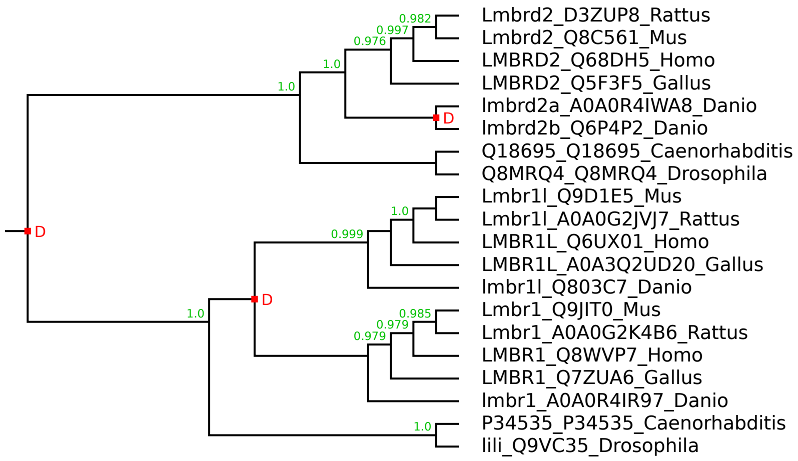

### pSLC.MFSD1 family

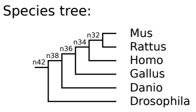


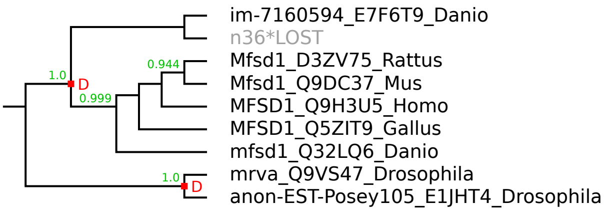

### pSLC.MFSD6 family

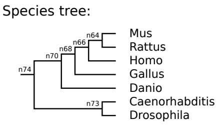


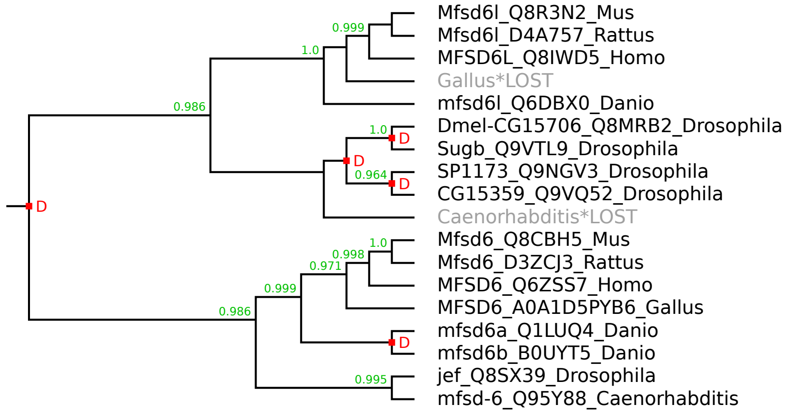

### pSLC.RFT1 family

Species tree:

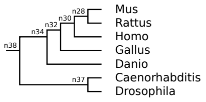


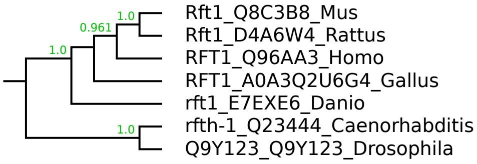

### pSLC.SIDT family

Species tree:

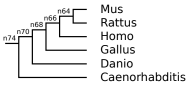


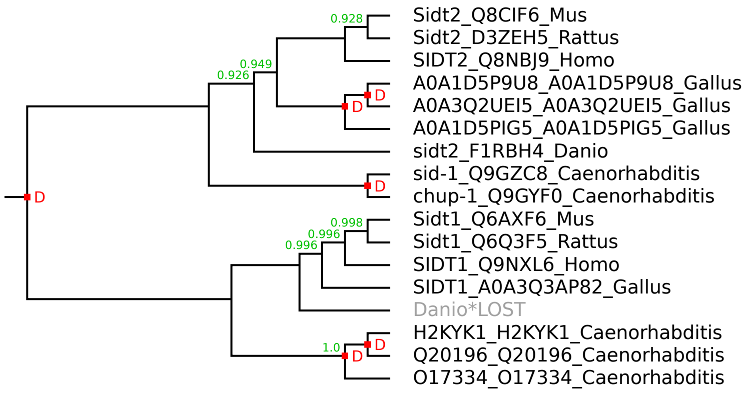

### pSLC.STAR family

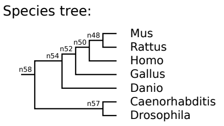


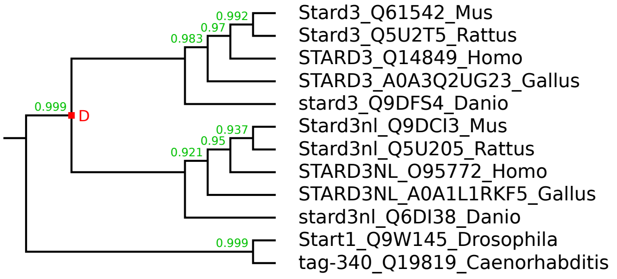

### pSLC.STRA family

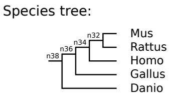


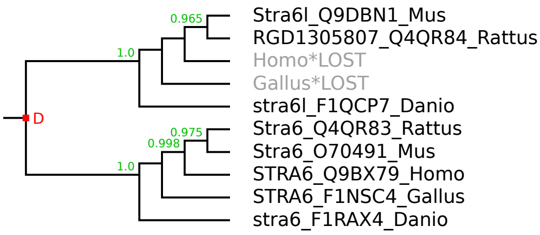

### pSLC.TMCO3 family

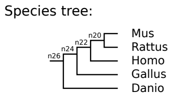


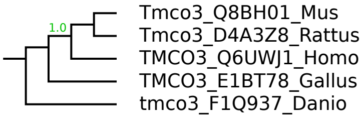

### pSLC.TMEM14 family

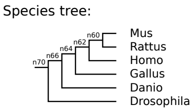


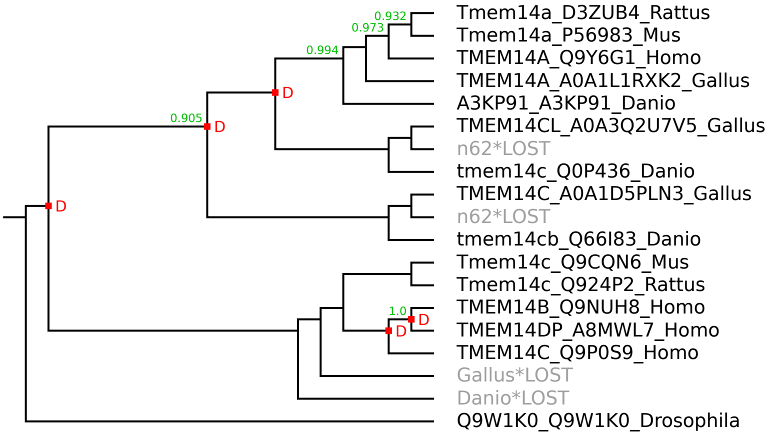

### pSLC.TMEM41-64 family

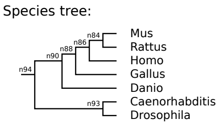


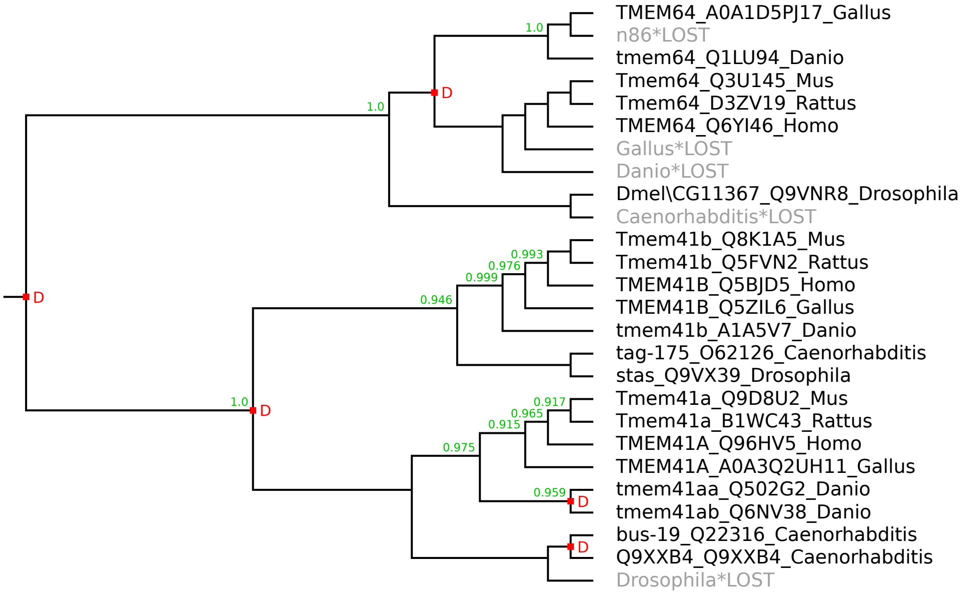

### pSLC.TMEM104 family

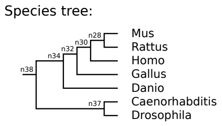


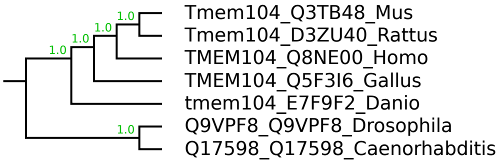

### pSLC.TMEM144 family

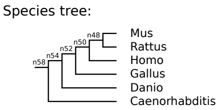


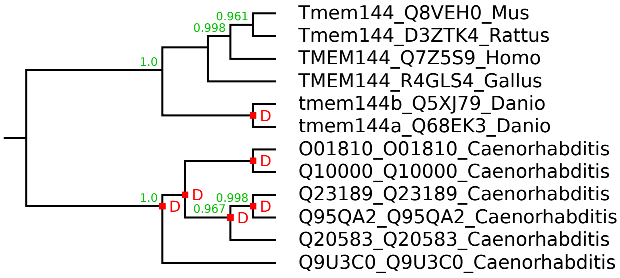

### pSLC.TMEM163 family

Species tree:

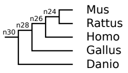


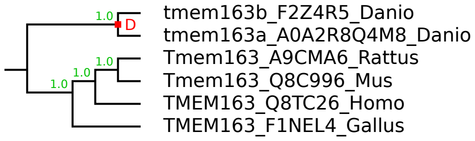

### pSLC.TMEM205 family

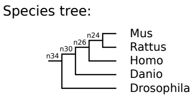


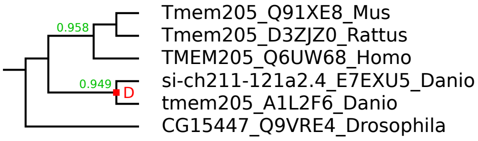

### pSLC.TMEM234 family

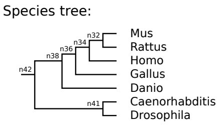


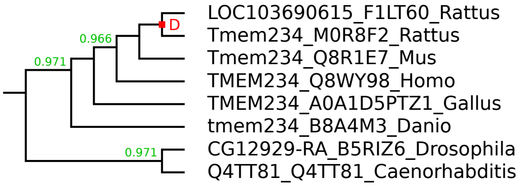

### pSLC.TMEM245 family

Species tree:

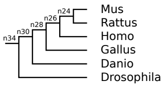


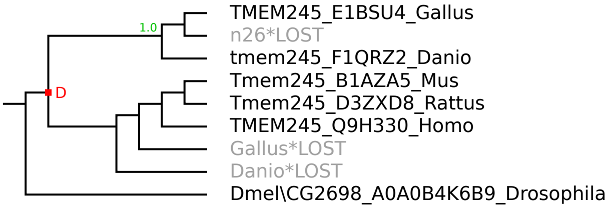

### pSLC.TSPO family

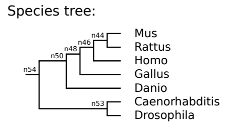


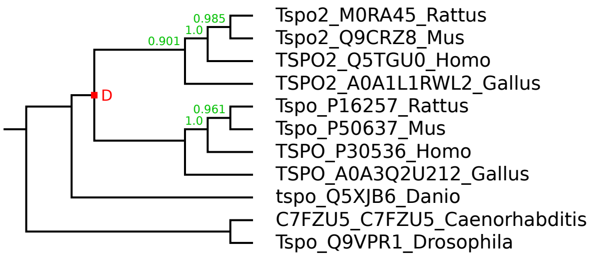

### pSLC.UNC93 family

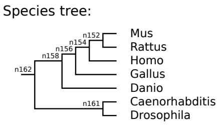


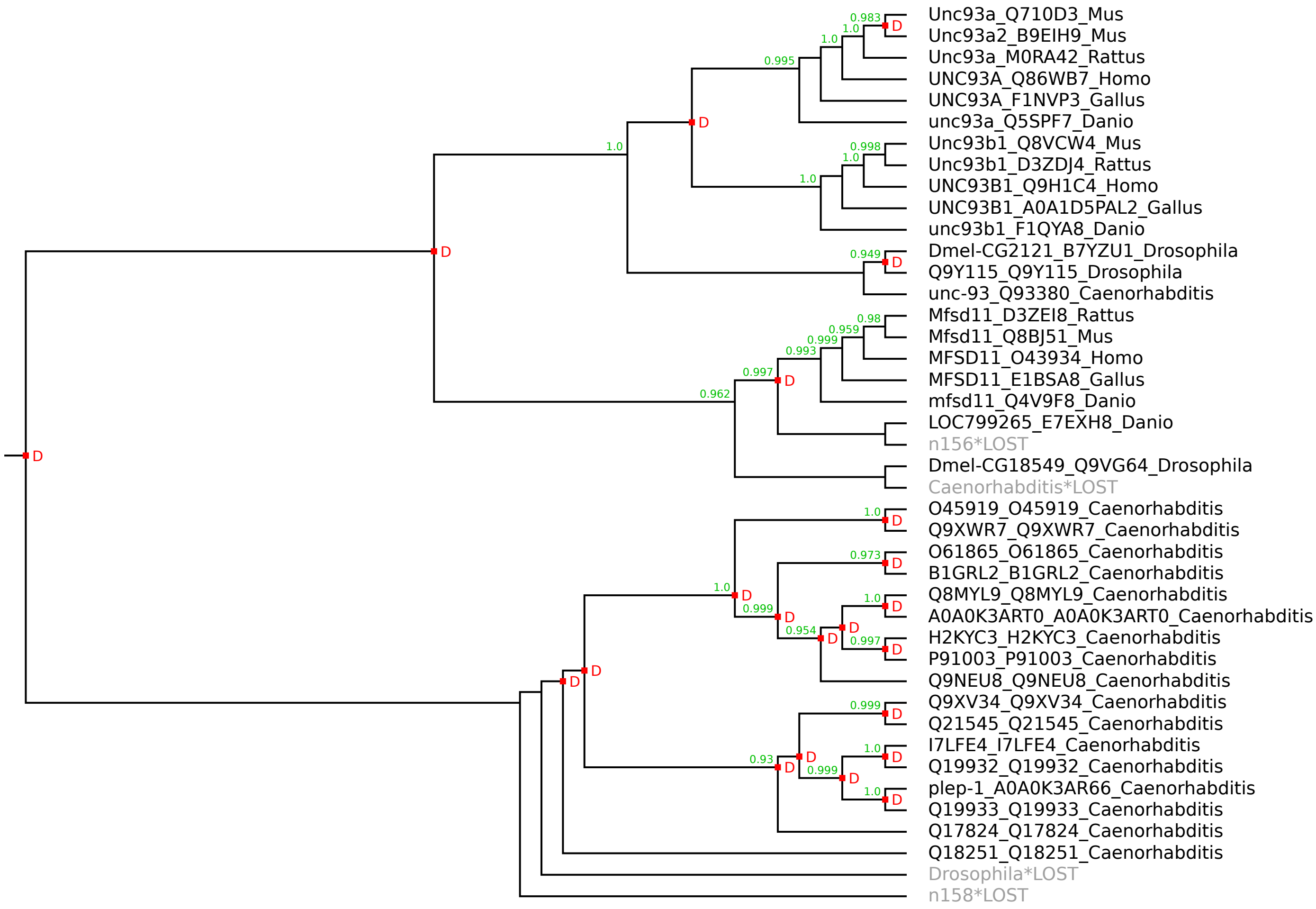

### pSLC.XK family

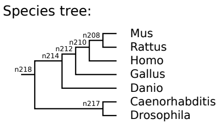


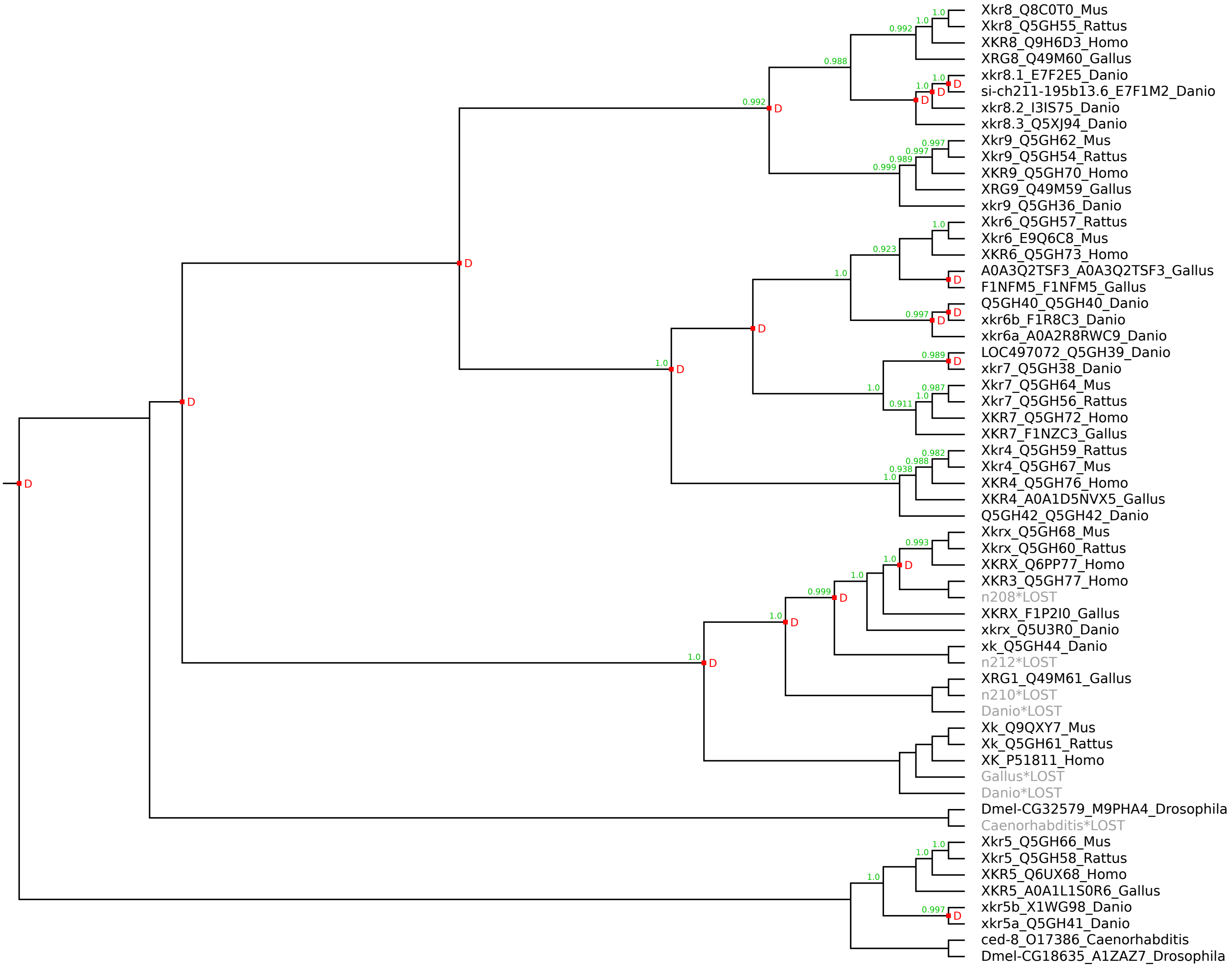
